# Generative modeling of spatially resolved perturbation responses with CONCERT

**DOI:** 10.1101/2025.11.08.686890

**Authors:** Xiang Lin, Zhenglun Kong, Boyang Fu, Soumya Ghosh, Manolis Kellis, Marinka Zitnik

## Abstract

Spatial perturbation transcriptomics measures gene expression responses to genetic or chemical perturbations in intact tissue. Each response depends on the intrinsic state of a cell and on its niche, the neighboring cells through which perturbation effects propagate. Here we present CONCERT, a niche-aware Gaussian process variational autoencoder that embeds each perturbation with the niche of the receiving cell and learns a spatial kernel for each perturbation, whose lengthscale sets how far the effect extends in tissue. We evaluate CONCERT in within-niche prediction, for held-out cells in the niche where a perturbation was measured, and in cross-slide prediction, for a slide from which the perturbation was withheld during training. On a SPAC-seq screen of hundreds of knockouts at single-cell resolution and on Perturb-map lung tumors at spot resolution, CONCERT predicts response magnitude and gene ranking more accurately than counterfactual models trained on dissociated cells, spatial perturbation models and spatial nearest neighbor aggregation. In dextran sodium sulfate colitis, where each mouse is sampled once, CONCERT reconstructs spatial gene expression at unmeasured time points, enabling longitudinal comparison that separates changes over time from differences between mice. The predictions recover the nonmonotonic course of injury, which peaks at day 9 and resolves by day 21. In ischemic stroke, CONCERT predicts responses to lesions of different sizes and locations, in single sections and across serial sections in three dimensions, and reproduces the expression patterns of the lesion core and perilesional tissue. Across lung tumor, colon and brain tissue, CONCERT predicts niche-aware counterfactual responses at locations where a perturbation was not measured and reconstructs missing spatial data.

## Main

The response of a cell to a perturbation depends on its intrinsic state and on its spatial context [1]. The niche of a cell, which comprises its neighboring cells and local microenvironment, shapes its transcriptional state through paracrine signals from neighboring cells [2], direct cell contacts [3], the local supply of nutrients [4] and immune surveillance [5]. In cancer immunotherapy, the tumor niche influences the response to treatment through the composition and activation state of infiltrating immune cells and through the local expression of checkpoint molecules such as PD-L1, which suppress T cell activity [6–8]. Dissociated single-cell assays remove cells from this context and cannot capture spatial effects that are important in development [9], immunity [10] and disease [11].

Spatial perturbation screens introduce genetic perturbations into cells in tissue, often *in vivo*, and read out perturbations and cellular states *in situ*, which preserves tissue context and the interactions between cells [12–14]. These assays measure non-cell-autonomous effects, in which a perturbation in one cell changes gene expression in its neighbors, and the spatial extent of these effects across niches (Supplementary Fig. 2). Dissociated single-cell assays cannot measure these effects because they remove cells from the tissue. Spatial profiling requires fixed or lysed tissue, so the same cell cannot be measured before and after perturbation. Counterfactual prediction, which infers gene expression under conditions that were not observed or cannot be tested experimentally, is therefore central to the analysis of these data.

Single-cell foundation models are pretrained on large cell atlases to learn cell and gene representations that transfer across tasks [15–18]. They are trained on dissociated scRNA-seq without spatial coordinates or neighborhood graphs, so their representations do not encode spatial context. Perturbation models such as scGEN [19], CPA [20], CellOT [21], GEARS [22], BioLORD [23] and PDGrapher [24] predict transcriptional responses to perturbations but treat each cell independently of its neighbors. In parallel, spatial transcriptomics methods deconvolve spots into cell types and identify spatial domains and expression patterns [25–33] but do not model responses to perturbations.

Methods for spatial perturbation data address parts of this problem. Vespucci [34], River [35] and Perturb-STNet [36] identify regions, genes, programs or regulators that respond to observed perturbations, but they cannot predict the effect of a perturbation at locations where it was not measured (limitation 1). Celcomen [37] separates regulation within and between cells and infers perturbation effects from learned gene–gene interactions, but these interactions are shared across the tissue, so the model assumes the same regulation at every location and cannot represent effects that change with the niche (limitation 2). stFormer [38], Morpheus [39] and CellAgentChat [40] perform *in silico* perturbations, each within a narrow scope. stFormer models ligand–receptor interactions with a transformer, Morpheus applies counterfactual optimization to spatial proteomic profiles for a single immune phenotype, T cell infiltration, and CellAgentChat simulates cell–cell communication with rule-based agents (limitation 3).

We introduce CONCERT, a generative model for spatial perturbation prediction built on a Gaussian process variational autoencoder (Fig. 1a). Given the gene expression profile, spatial coordinates and perturbation identity of an unperturbed cell, CONCERT predicts its response gene expression (rGEX), the expression profile of the cell after perturbation, conditioned on its niche. CONCERT consists of three modules (Fig. 1b). The disentanglement module separates spatial from intrinsic variation in expression and the perturbation effect from the basal state of the cell, so that a perturbation can be applied *in silico* to any cell at any location (addressing limitation 1). The perturbation-signal module conditions the perturbation effect on the niche and separates biological response from technical shifts caused by guide delivery and by variation between slides (addressing limitation 2). The generative module decodes these components into rGEX. CONCERT handles several perturbation types and learns their spatial effects in 2D and 3D tissue (addressing limitation 3). It can also impute missing cells and predict their responses in regions left unmeasured by sparse sampling or tissue damage.

**Figure 1:**
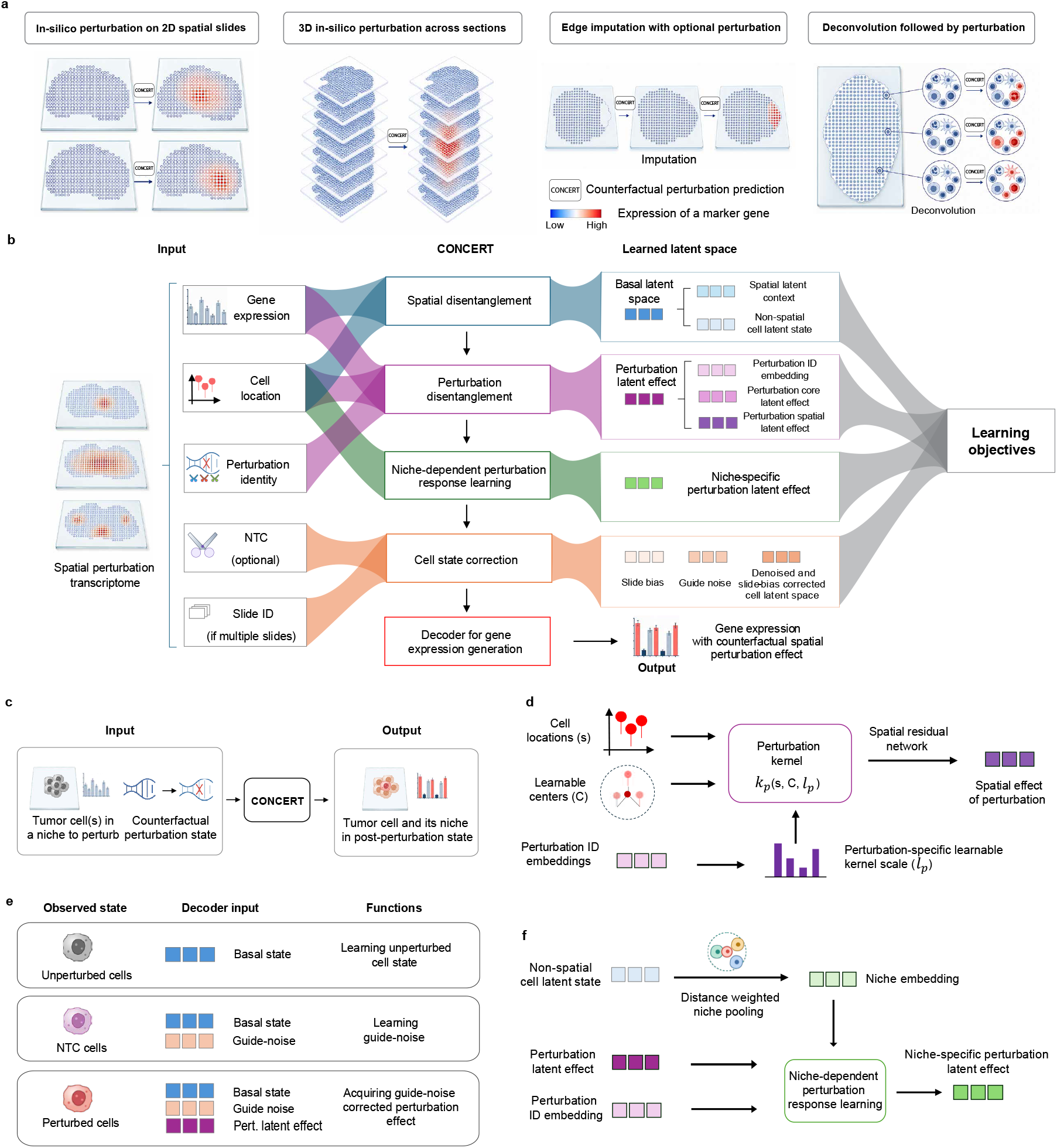
Overview of CONCERT functions, architecture, and key modeling components. **a**, CONCERT can conduct spatially aware counterfactual prediction in four scenarios: 1) at specified locations within a two-dimensional section, 2) across registered sections in three dimensions, 3) in imputed missing regions, and 4) in cell-level states deconvolved from mixed measurements. **b**, Architecture of CONCERT. From cells’ gene expression, coordinates, slide identity, guide status (denoted as NTC in panel b) and perturbation identity, CONCERT learns a perturbation-free basal representation, split into a non-spatial latent for cell identity and a spatial latent for tissue variation. The perturbation response is decomposed into a spatially invariant core effect and a location-dependent spatial effect, modulated by the states of nearby cells, and combined with basal expression with the guide and slide corrections. A negative-binomial model generates the output. **c**, Counterfactual inference. Parameters are fixed after training; the perturbation state of a control cell is replaced by a target identity while its cell state and spatial context are retained. Then CONCERT predict the response gene expression to the target perturbation in a specific location of tissue. **d**, Perturbation-specific spatial effects. Each perturbation embedding **e**_*p*_ (for a perturbation *p*) generates a core response and a length scale ℓ_*p*_. The coordinates *s* of a queried cell are compared with learned reference centers *C* by radial-basis functions, and the resulting features pass with **e**_*p*_ through a residual network to learn the spatial effect of perturbation. RBF is used as the default kernel function. **e**, Guide and slide corrections. Unassigned (unperturbed) cells identify basal expression; non-targeting-control (NTC) cells identify the technical shift from guide delivery, separating it from the biological response of knockout cells. **f**, Niche-dependent responses. The non-spatial latents of a cell’s nearest neighbours are pooled with distance weighting into *z*_niche_; Niche-FiLM combines it with **e**_*p*_ to rescale and shift the response to the perturbation *p*.

We evaluate CONCERT in settings that differ in which cells and slides are withheld from training (Fig. 1a and Supplementary Note 7). In *within-niche prediction*, a model predicts the response of held-out unperturbed cells to a perturbation measured in other cells of the same niche, which tests whether it can place a perturbation effect at an unmeasured location. In *cross-slide prediction*, the perturbation is withheld from an entire target slide and the model transfers the response learned on other slides, which tests whether the learned effect is a property of the perturbation and not of a single slide. We benchmark CONCERT on SPAC-seq, a screen of hundreds of knockouts profiled at single-cell resolution across five slides [41], and on Perturb-map, Visium profiles of mouse lung tumors carrying CRISPR knockouts of *Jak2, Tgfbr2* and *Ifngr2* [12]. In each setting, CONCERT predicts the magnitude and gene ranking of the response more accurately than competing models. We next apply CONCERT to a MERFISH atlas of dextran sodium sulfate colitis, in which each mouse is sampled once [42]. CONCERT reconstructs the spatial gene expression of each mouse at unmeasured time points, which separates changes over time from differences between mice and recovers the rise of inflammation to day 9 and its resolution by day 21. We also apply CONCERT to a mouse model of photothrombotic stroke [43]. CONCERT predicts rGEX for lesions of sizes and locations that cannot be controlled experimentally, in two and three dimensions, and its predictions reproduce the expression patterns of the lesion core and perilesional tissue. These results show that CONCERT performs niche-aware prediction across tissues, perturbation types and experimental designs.

## Results

### Generative model of spatial perturbations

CONCERT predicts the response gene expression (rGEX) of a cell from its expression profile, spatial location and niche (Fig. 1a). The model consists of a disentanglement module, a perturbation-signal module and a generative module, which we describe in turn (Fig. 1b). For counterfactual prediction, CONCERT changes the perturbation label of a cell, or a continuous attribute such as dose or time. The model then adapts the new perturbation effect to the cell and its niche and decodes the corresponding rGEX (Fig. 1b,c).

The disentanglement module separates information along two axes. Along the perturbation axis, CONCERT embeds the perturbation identity, such as a gene knockout or a control condition, with attributes such as dose or time, and keeps this perturbation embedding separate from the basal embedding, which describes the cell without perturbation. This separation allows the model to apply a perturbation to a cell that was never exposed to it.

Along the spatial axis, CONCERT splits the basal embedding into a non-spatial component for intrinsic cell identity and state and a spatial component for variation associated with location (Fig. 1b). CONCERT models tissue space with two fields. The basal spatial field describes smooth anatomical variation independent of perturbation by mapping cell coordinates to random Fourier features [44], sine and cosine patterns at several spatial scales that approximate a stationary Gaussian process kernel. The perturbation field describes how the effect of each perturbation varies across the tissue, as the sum of a core effect shared across locations and a spatial adjustment that varies with location. The adjustment depends on the distance from each cell to a set of learnable reference locations through a radial basis function kernel, whose perturbation-specific lengthscale sets how fast the influence of a reference location decays with distance. A short lengthscale describes a localized effect and a long lengthscale an effect that extends over a broad tissue region (Fig. 1d). The lengthscale is predicted from the perturbation embedding by a mapping shared across perturbations, so related perturbations learn similar spatial ranges and perturbations observed in few cells borrow their range from related perturbations.

The perturbation-signal module adjusts the perturbation effect to the niche of each cell and separates this biological effect from technical variation. First, the module pools the non-spatial basal states of the nearest neighbors of a cell, weighted by distance, into a niche embedding, a summary of the composition and state of the surrounding cells. A feature-wise linear modulation (FiLM) layer [45] uses this embedding to rescale and shift the perturbation effect, so the same perturbation can act more strongly, more weakly or in the opposite direction in different niches (Fig. 1f). Second, in screens with non-targeting controls, the module uses the control groups to separate biological from technical effects. Cells without an assigned guide define basal expression, cells with a non-targeting guide define the expression shift caused by guide delivery, and cells with a gene-targeting guide receive this shift together with the target-specific effect (Fig. 1e). A learned term for each slide absorbs residual technical variation between tissue sections.

The generative module combines these components to predict rGEX. A variational encoder maps the observed expression of a cell to a low-dimensional latent state, and a decoder maps this state to basal expression. The perturbation effect, the guide shift and the slide term act as shifts on the log scale of this basal expression, and the result defines the mean of a negative binomial distribution over gene counts. The decoder reconstructs measured expression during training and generates counterfactual expression at inference, after the perturbation identity, its location or a continuous attribute is changed. By varying the number and arrangement of cells perturbed *in silico*, the model predicts responses to lesion sizes and locations that were not measured, and by changing a continuous attribute, it predicts responses at unmeasured times or doses. The model can also infer the basal state at unmeasured locations from their coordinates and neighboring cells, which allows it to reconstruct damaged or missing tissue regions and predict how the reconstructed cells respond to a perturbation.

### A benchmark for spatial perturbation prediction

We benchmarked CONCERT on SPAC-seq, a spatial perturbation screen of hundreds of knockouts measured at single-cell resolution across five tissue slides [41]. Because the perturbations are introduced *in vivo*, the locations that receive each perturbation cannot be controlled, and many combinations of perturbation and location are not measured (Supplementary Fig. 1). We evaluated CONCERT in within-niche and cross-slide prediction (Supplementary Note 7). In each task, the model receives the location, unperturbed expression and target perturbation of a held-out unperturbed cell, which we call a *predicted cell*, and its prediction is compared with measured cells that have the target perturbation. We refer to the expression states before and after perturbation as the *source state* and *target state*. In *within-niche prediction*, a spatial patch of unperturbed cells is withheld from training and predicted into each target state measured in the same niche, which tests prediction at unmeasured locations. In *cross-slide prediction*, a perturbation is withheld from an entire target slide but observed on other slides, and its response is predicted for the unperturbed cells of the target slide, which tests whether the learned effect is a property of the perturbation independent of its slide. Cross-slide prediction tests spatial transfer, not the prediction of unseen perturbations.

We define niches by rescaling the coordinates of each slide to a common frame and dividing the slide into a regular 6 *×* 6 grid, which gives 36 niches per slide, each spanning one sixth of the slide along each axis. The grid is used only for evaluation and is not an input to the model. Because a knockout observed in few cells gives an unreliable reference for evaluation, we filter perturbations by the number of cells in which they are observed. For within-niche prediction, a knockout is eligible on a slide if it is observed in at least 200 cells on that slide and in at least 20 cells within one niche, and it is scored in a niche only when that niche contains at least 20 knockout cells and at least 20 held-out unperturbed cells, which gives 87 perturbations. For cross-slide prediction, all cells with the target knockout are withheld from the target slide, so a knockout is eligible if it is observed in at least 20 cells on the donor slide, which provide training examples of its effect, and in at least 100 cells on the target slide, which provide a reliable measured response for evaluation. All eligible knockouts are predicted, which gives 85 knockout–target-slide pairs.

We evaluate performance with the root-mean-square error (RMSE) and the Spearman rank correlation between predicted and measured log-fold changes relative to unperturbed control cells. RMSE measures the magnitude of the predicted response, and the rank correlation measures whether the model orders the responding genes correctly. We evaluate log-fold changes instead of total expression because the perturbation response is usually a small fraction of measured expression, whereas the basal state accounts for most of the signal. Metrics computed on total expression would be dominated by basal expression and could appear favorable even when the perturbation effect is predicted poorly.

We compared CONCERT with three groups of methods. Reference strategies make predictions without a fitted model, using the mean of measured cells in the target state, the mean of cells in the same niche, or the mean of the nearest measured cells in the target state, selected by spatial or transcriptomic distance (Supplementary Note 8 discusses their limitations). Perturbation models trained on dissociated cells (scGEN [19], BioLORD [23], GEARS [22] and STATE [46]) predict a response from expression alone and have no access to the location of a cell in tissue. Spatial perturbation models (Celcomen [37], CellAgentChat [40] and Steamboat [47]) represent spatial or cell–cell interaction structure and are the closest published comparators. Some perturbations are outside the scope of these methods, such as knockout targets missing from the gene panel of Celcomen or the ligand–receptor database of CellAgentChat, and each method is scored on the perturbations it covers.

Sensitivity analyses of the hyperparameters of the spatial components on SPAC-seq and Perturb-map show that the performance of CONCERT changes little across a broad range of settings and that the best settings differ between cell-level and spot-level data (Supplementary Fig. 3, Supplementary Fig. 4, Supplementary Fig. 5, Supplementary Fig. 6; Supplementary Note 9).

### Within-niche prediction

We first evaluated within-niche prediction (Fig. 2a). The rank correlation of log-fold changes separates the methods most clearly. Across the 87 knockouts, CONCERT reaches a rank correlation of 0.427, compared with 0.340 for the target-mean baseline, 0.336 for scGEN, 0.305 for GEARS and 0.301 for BioLORD, whereas the three spatial perturbation models and STATE reach correlations at or below zero (Fig. 2b). When scores are rescaled within each knockout, so that easy and difficult knockouts have equal weight, CONCERT has the highest score on most knockouts (Fig. 2c). Among the methods that scored each knockout, CONCERT has a mean rank of 1.54 and ranks first on 78% of knockouts (Fig. 2d). Its margin over each competing method is positive on most knockouts, with a median margin of 0.069 over the target-mean baseline (Fig. 2e). CONCERT also has the lowest mean log-fold-change RMSE among the twelve methods (4.97) and the lowest error on 57 of the 87 knockouts (Extended Data Fig. 1). The GEX kNN and mean niche baselines rank fourth and sixth by RMSE, but their rank correlations are close to zero (0.092 and 0.096). These baselines reproduce the dominant basal expression, which keeps their error low, but they do not capture the perturbation-specific response. CONCERT models the perturbation effect separately from the basal state and fits it to measured log-fold changes (Methods), so its predictions are not dominated by basal expression. Paired Wilcoxon signed-rank tests show that CONCERT improves significantly over every competing method in rank correlation and in RMSE (Supplementary Tables 1 and 2).

**Figure 2:**
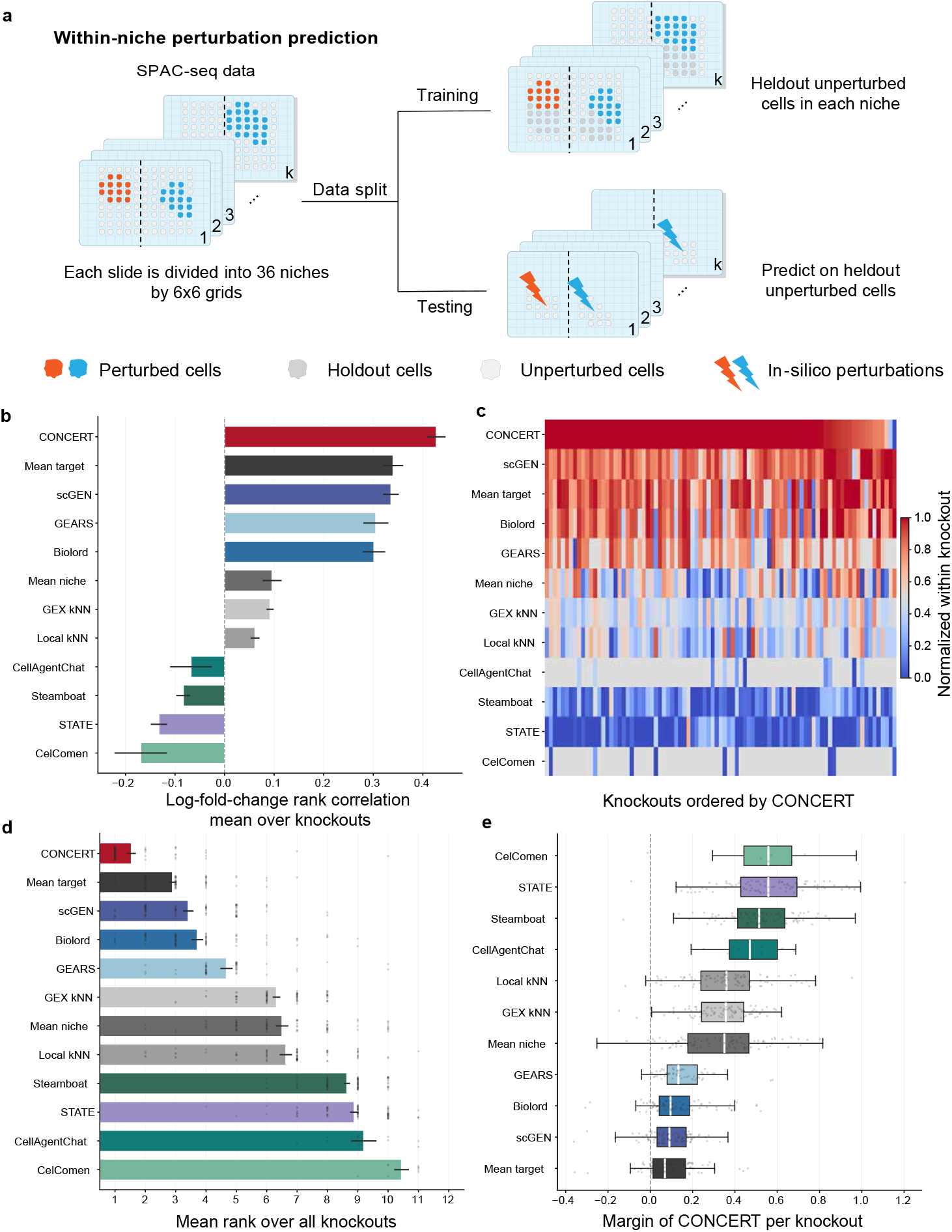
Within-niche prediction on SPAC-seq. A spatial patch of unperturbed cells is withheld from training and predicted into a knockout state carried by cells of the same local niche. Accuracy is the Spearman rank correlation between the predicted and the measured log-fold change against unperturbed control cells; higher is better. **a**, Task schematic. Each of the *k* slides is divided into *p* niches. A patch of unperturbed cells is held out of training (heldout cells in grey). At test time, those cells are perturbed *in silico* into each target perturbation state and the prediction is compared with the cells that actually carry that perturbation. Here, perturbed cells (red and blue) carry a perturbation in the data; unperturbed cells (white) are the remaining control cells. **b**, Rank correlation of each method, the mean over the 87 knockouts with the standard error across them. Methods are averaged over the knockouts for which they returned a prediction. **c**, Every knockout as one column, with the methods rescaled inside that column so that 1 marks the best method and 0 the worst. Grey marks a knockout a method left unpredicted, and the columns are ordered by the score of CONCERT. **d**, Mean rank of a method inside a knockout, taken over the methods that scored that knockout, with rank 1 the best and one dot per knockout. **e**, Margin of CONCERT over each competing method on every knockout both scored, positive where CONCERT leads. The box spans the quartiles and each dot is one knockout. Mean target takes the mean of 50 measured cells drawn at random from the target state and Mean niche the mean of the cells in the same niche; Local kNN and GEX kNN average the nearest cells selected by spatial and by transcriptomic distance. The corresponding RMSE is in Extended Data Fig. 1.

We next related the learned spatial parameters to the spatial distribution of each knockout in tissue. CONCERT learns two spatial parameters for each perturbation, the niche gain, which bounds how strongly the niche can modulate the effect, and the lengthscale, which sets how far the effect extends. Across the 87 knockouts, partial rank correlations controlled for the number of knockout cells show that the niche gain increases and the lengthscale decreases with the fraction of section area reached by the knockout cells (Extended Data Fig. 5 and Supplementary Note 5). CONCERT thus learns niche-dependent responses for knockouts spread across many microenvironments and spatially uniform responses for knockouts confined to a single cluster of cells. A model trained in the same configuration on a tumor section from Perturb-FISH, an imaging-based CRISPR screen [13], shows the same pattern, with a higher niche gain and a shorter lengthscale for knockouts whose cells reach a larger fraction of the section (Supplementary Fig. 28).

### Cross-slide prediction

We next evaluated cross-slide prediction, in which the cells of a knockout are masked on a target slide during training and the response is predicted on that slide (Fig. 3a). Across the 85 knockout– target-slide pairs, CONCERT reaches a rank correlation of 0.703 between predicted and measured log-fold changes, compared with 0.627 for the target-mean baseline, 0.602 for BioLORD, 0.532 for GEARS and 0.496 for scGEN and STATE (Fig. 3b). CONCERT ranks first on all 85 pairs (Fig. 3c,d), has a higher rank correlation in each of the 759 comparisons in which a competing method scored the same pair, and exceeds the target-mean baseline by a median of 0.076 (Fig. 3e). Baselines that average nearby cells perform poorly in this setting. Because no cell on the target slide has the target knockout, the nearest cells that local kNN averages are unperturbed, so its prediction reproduces basal expression and reaches a rank correlation at or below zero. The three spatial perturbation models also reach correlations at or below zero. GEX kNN, which averages the measured knockout cells most similar in expression to each predicted cell, reaches 0.323.

**Figure 3:**
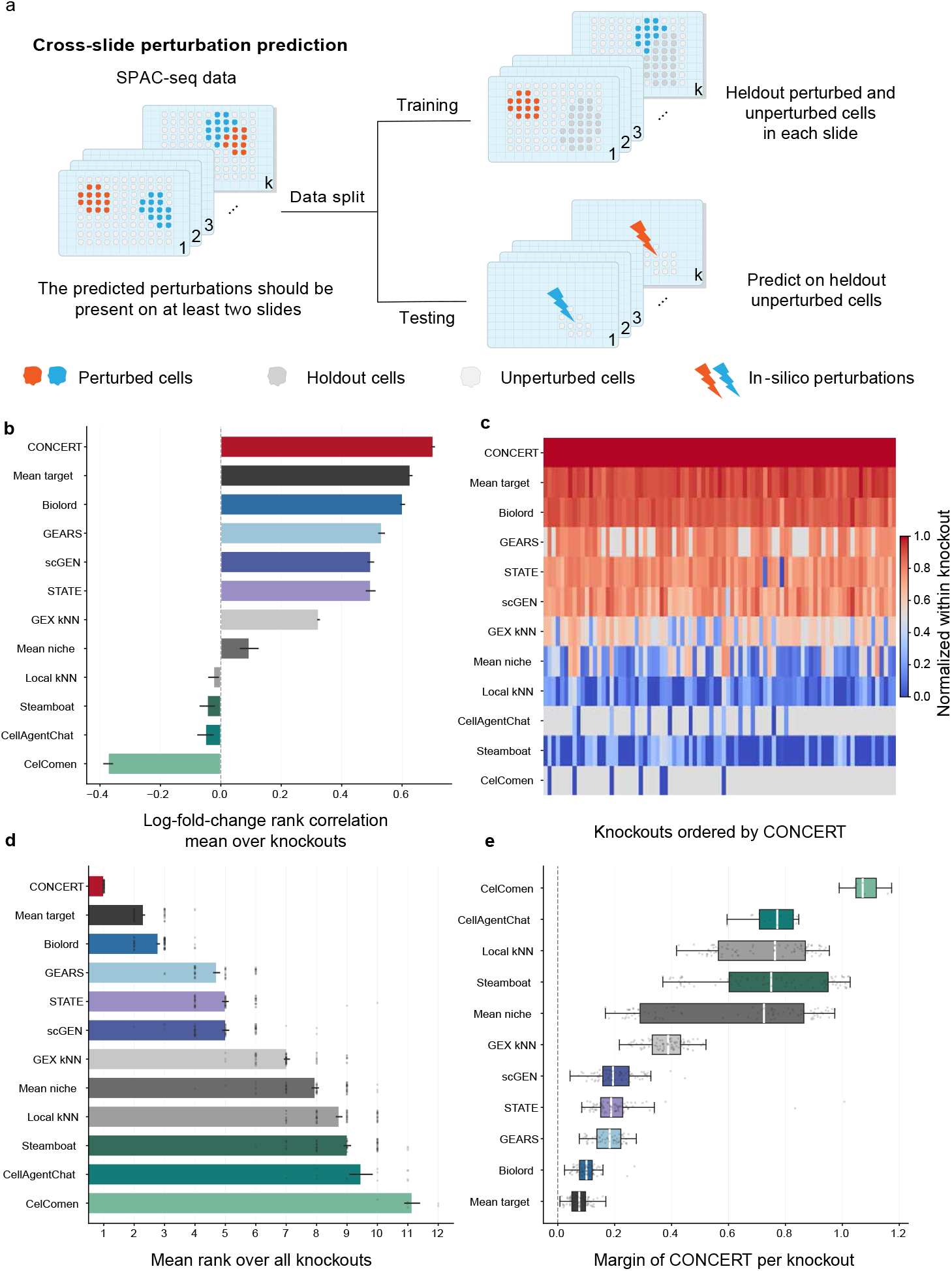
Cross-slide prediction on SPAC-seq. The perturbed cells of a knockout are masked on one target slide while that knockout remains observed on the donor slides, and the response is predicted on the target slide from its unperturbed cells. Evaluation metric, color scheme, and competing methods are the same as in Fig. 2. **a**, Task schematic. Each slide carries more than one perturbation. For a heldout perturbation, both its perturbed cells and a patch of unperturbed cells on the target slide are withheld from training (heldout cells in grey; perturbed cells in red and blue). At test time, the withheld unperturbed cells are perturbed *in silico* into that perturbation. The perturbation is observed on the donor slides in training, so this is a test of spatial transfer. **b**, Rank correlation of each method, the mean over the 85 knockout–target-slide pairs with the standard error across them. **c**, Every pair as one column, with the methods rescaled inside that column so that 1 marks the best method and 0 the worst. Grey marks a pair a method left unpredicted, and the columns are ordered by the score of CONCERT. **d**, Mean rank of a method inside a pair, taken over the methods that scored that pair, with rank 1 the best and one dot per pair. **e**, Margin of CONCERT over each competing method on every pair both scored, positive where CONCERT leads. The box spans the quartiles and each dot is one pair. The corresponding RMSE is in Extended Data Fig. 2.

For response magnitude, CONCERT has the lowest RMSE (3.81) and the lowest error on 64 of the 85 pairs (Extended Data Fig. 2). A control that predicts no change in expression has an RMSE of 4.47 on the same records, and six of the seven published models have a higher RMSE than this control. Paired Wilcoxon signed-rank tests show that CONCERT improves significantly over all eleven competing methods in rank correlation and RMSE (Supplementary Tables 1 and 2). Because the knockout was masked on the target slide during training, these results indicate that the effect learned by CONCERT is a property of the perturbation and not of a single slide.

### Deconvolution of mixed spots and imputation of missing regions

Many sequencing-based spatial assays, including Perturb-map, measure several cells at each spot, so each observed profile is a mixture of cell states [48]. Because CONCERT predicts the expression of a given cell state at a given position, it can deconvolve a mixed spot, that is, decompose it into its component states, and represent the recovered cells as sub-spots, which increases spatial resolution without additional experiments. We first tested deconvolution on pseudo-bulk spots formed by aggregating SPAC-seq cells, for which the true composition of every spot is known. Non-negative least squares (NNLS) with the location-specific state signatures of CONCERT reduced the mean absolute error of the estimated composition from 0.297, obtained with NNLS on average signatures, to 0.157, and increased the Pearson correlation with the true composition from 0.415 to 0.499. The error decreased in 79% of mixed spots, by 0.141 on average, and the predicted knockout fractions agreed with the true fractions across the full range of mixture proportions (Supplementary Fig. 9). We next applied CONCERT to Perturb-map, where spots around a knockout patch can contain knockout cells, unperturbed tumor cells and normal cells. These spots correlated only moderately with the profiles of each of the three populations (0.588, 0.632 and 0.576), which reflects this mixing. After deconvolution and imputation, CONCERT represented the mixed region as sub-spots, with each component placed at its own position between two observed spots (Supplementary Fig. 10, Supplementary Note 6).

CONCERT can also reconstruct tissue regions that were damaged or not measured. To this end, we withheld contiguous regions of SPAC-seq tissue, instead of individual cells, so that each region had to be reconstructed from its spatial and transcriptional surroundings. The withheld regions were five knockout patches, so reconstruction required inferring the basal state from the neighborhood and the perturbation effect from cells of the same knockout elsewhere on the slide. CONCERT achieved the highest accuracy among the tested methods, with a mean per-cell Spearman correlation of 0.23, compared with 0.20 for SpaIM [49], 0.19 for Tangram [27] and 0.07 for SpatialScope [50]. For the log-fold change, the mean Spearman correlation was 0.65 for CONCERT, compared with 0.61, 0.56 and 0.35, respectively, and ranged from 0.48 to 0.72 across patches (Supplementary Fig. 11). Unlike these imputation methods, CONCERT also models the perturbation effect, which allows it to predict the response of the reconstructed cells.

### Niche-dependent perturbation responses in lung tumors

We next used Perturb-map, in which each spot can contain several cells, to test CONCERT on spot-level data and to study how the niche shapes perturbation responses in mouse lung tumors [12]. In a within-niche benchmark against five published methods, CONCERT achieved the highest log-fold-change rank correlation and the lowest RMSE for each knockout (Fig. 4j). Its rank correlations were 0.475, 0.445 and 0.570 for *Tgfbr2, Ifngr2* and *Jak2*, with RMSEs of 3.301, 4.057 and 1.408. The strongest competing method by rank correlation, scGEN, reached 0.348, 0.339 and 0.256, with RMSEs of 4.932, 5.142 and 2.746. Paired Wilcoxon signed-rank tests show that CONCERT improves significantly over every competing method in rank correlation and RMSE (Supplementary Tables 1 and 2). These results extend the SPAC-seq benchmark from cell-level to spot-level data.

**Figure 4:**
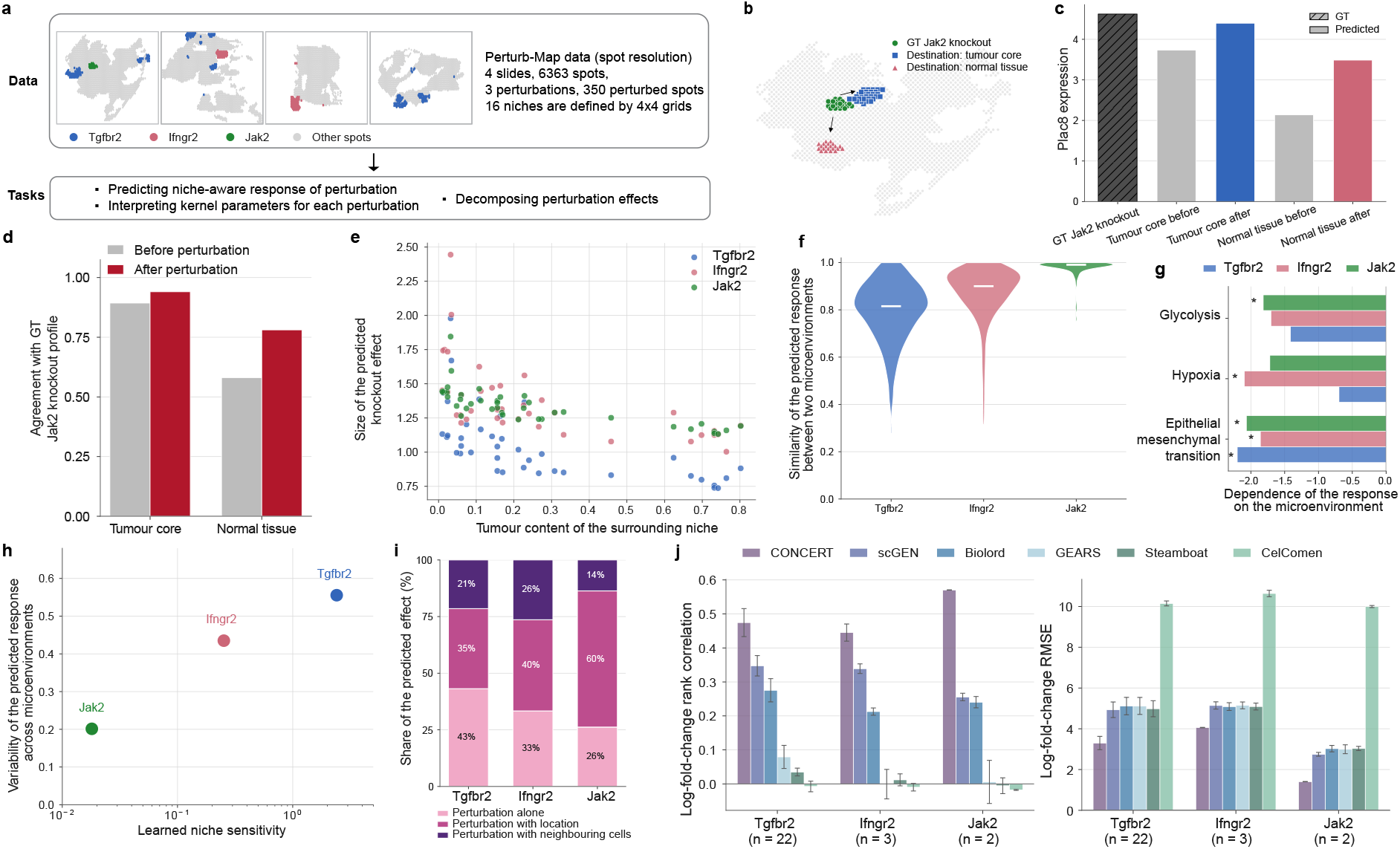
CONCERT predicts how a knockout’s response changes with the microenvironment. Perturb-map lung tumor slides [12]. GT is the observed ground truth; every prediction is shown beside it. **a**, The four slides and the knockouts each carries. Each slide carries one or two of the three knockouts, so every comparison between knockouts is across slides. **b–d**, *In silico* transfer of a knockout between niches. Jak2-knockout spots from a tumor core are moved *in silico* into another tumor core and into a tumor surface surrounded by normal tissue. **b**, the source spots and the two destinations. **c**, predicted Plac8 expression at each destination, beside the measured Jak2 value. **d**, agreement of the whole predicted profile with GT Jak2 at each destination. **e–g**, Each knockout placed into 40 sampled niches of 20 spots. **e**, the predicted effect, the mean absolute log-fold change against the same cells unperturbed, against the tumor content of the surrounding neighborhood. **f**, similarity of a knockout’s response between two niches. **g**, hallmark programmes whose response tracks niche composition, showing every set reaching *q <* 0.05 in at least one knockout; asterisks mark *q <* 0.05. **h**, the learned niche sensitivity of each knockout against how much its predicted response moves between niches. **i**, share of each knockout’s predicted effect contributed by the perturbation alone, by the perturbation with the cell’s location, and by the perturbation with the expression of neighboring cells; the three sum to 100% by construction and are mean absolute contributions. **j**, CONCERT and five published methods predict each knockout, scored against the measured perturbed spots by the rank correlation (left) and root mean squared error (right) of the log-fold-change vector. The observed log-fold change uses as reference the unperturbed tumor spots of the same niche that lie outside every held-out patch, or the slide-wide such spots when a niche holds fewer than five, so the reference and the predicted spots are disjoint. Bars are means over niche-patch records pooled across slides, error bars the s.e.m. across records, and *n* per knockout is on the axis. All methods are scored on an identical set of records.

We then used CONCERT to study how the niche changes perturbation responses, first by moving the same perturbed patch between niches and then by testing which transcriptional programs vary with niche composition.

#### In silico transfer of a knockout between niches

On slide GSM5808054, we applied the *Jak2* knockout, observed in spots of the tumor core, *in silico* to two destinations, a second tumor-core region of 31 spots and a tumor surface region of 21 spots surrounded by normal tissue (Fig. 4b). Although the perturbation was identical, CONCERT predicted different responses in the two niches. *Plac8*, which is higher in the observed *Jak2*-knockout spots, increased from 3.74 to 4.40 in the tumor-core destination, close to the measured knockout value of 4.62, and from 2.14 to 3.49 at the tumor surface (Fig. 4c). Across the full transcriptome, the predicted profile agreed more closely with the observed *Jak2*-knockout profile in the tumor-core destination than at the tumor surface (*R*^2^ = 0.940 and *R*^2^ = 0.781; Fig. 4d). The same perturbation therefore produced a response close to the observed knockout state in the tumor core and a distinct response at the tumor surface, where basal *Plac8* expression is lower.

#### Responses across many niches

Comparing two niches cannot show whether niche dependence is a general pattern, so we placed each knockout *in silico* into 40 niches of 20 spots sampled across the slide. For each knockout, the magnitude of the predicted response, measured as the mean absolute log-fold change relative to the same spots without perturbation, decreased as the fraction of tumor cells in the niche increased (Spearman *ρ* = −0.82 for *Tgfbr2*, −0.73 for *Ifngr2* and −0.85 for *Jak2*; Fig. 4e). Responses predicted in different niches were least similar for *Tgfbr2*, which indicates that its effect depends most on the niche (Fig. 4f). We then asked which programs in the predicted response change with the tumor content of the niche, after removing differences in the overall magnitude of the response. Gene set enrichment analysis against the Hallmark collection (Methods) identified epithelial–mesenchymal transition (EMT) as the program most consistently associated with niche composition, with a weaker EMT response in tumor-rich niches for all three knockouts (Fig. 4g). For *Tgfbr2*, this result is consistent with the established role of TGF-*β* signaling in EMT [51], and it points to EMT as a program through which the niche may shape perturbation responses. The niche gain learned for each knockout agreed with the variation of its predicted response across the 40 niches (Fig. 4h), and a decomposition of the predicted effect showed that the terms involving location and neighboring cells account for most of the response for each knockout (Fig. 4i).

### Time course of inflammation and recovery in colitis

We applied CONCERT to a MERFISH atlas of colitis induced by dextran sodium sulfate (DSS), in which colon tissue was collected at day 0 (healthy) and at days 3, 9 and 21 [42]. Sampling is terminal, so each of the 15 mice was measured at a single time point, with 3, 4, 6 and 2 mice sampled at days 0, 3, 9 and 21 (Fig. 5a). Changes over time are therefore confounded with differences between mice, especially when the same tissue region is compared across days.

**Figure 5:**
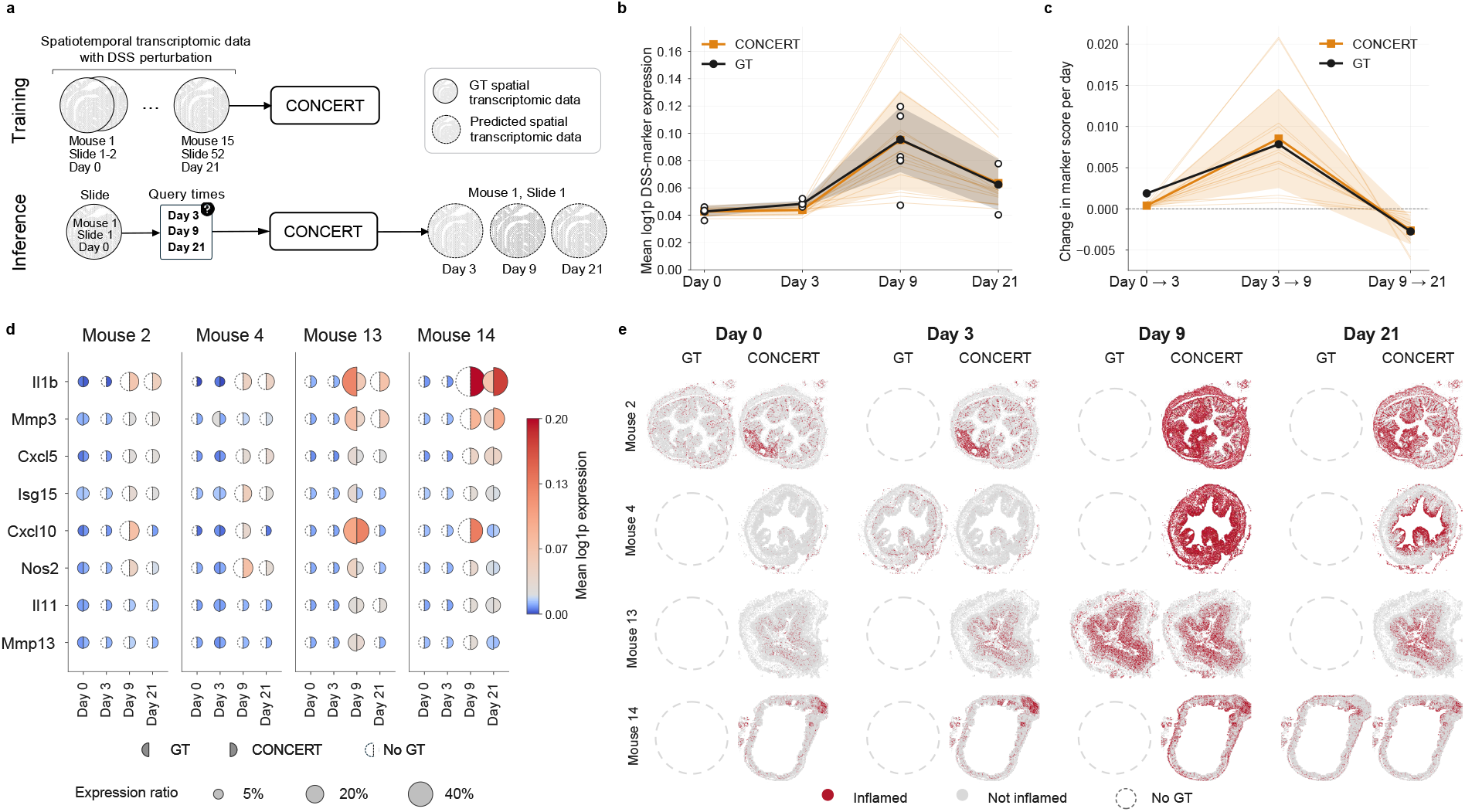
CONCERT reconstructs the time course of DSS (dextran sodium sulfate) colitis from cross-sectional snapshots. GT indicates the observed ground truth and No GT marks a day at which that animal was never sampled. **a**, Experimental design. The overall samples are stratified by mice, the tissue slices of each mouse, and sampling day of each mouse, with DSS administered on day 3. Sampling is terminal, so each of the 15 mice is measured at exactly one day and no animal’s trajectory is ever observed. This is the issue that we want to address with CONCERT. **b**, Population trajectory of the inflammation score: the trend of the mean log1p expression of the 17 inflammatory marker genes and the trend of that for each mouse. Black stands for GT and orange stands for CONCERT. The thin lines indicate the prediction of individual mice, and the bands indicate the 1 s.d. across replicates. **c**, Velocity of the same score, the change per elapsed day within each interval; the dashed line marks zero. **d**, Marker-level profiles for four representative mice, one per sampled day. Rows are the eight panel markers with a GT fold change. The columns are the four virtual days. Every dot is split at one position: left half GT, right half CONCERT. Colour is mean log1p expression on a shared scale; area is the fraction of expressing cells. For CONCERT, it is the detection probability *P* (count *>* 0) = 1 −*e*^−*µ*^. **e**, Spatial reconstruction of the inflamed region for the same mice in panel d. A cell is called inflamed (red) when its marker score exceeds a healthy-tissue cutoff. The value *r* indicates the inflamed fraction of that section. The representative animals and sections were chosen for structural completeness and never for agreement with the prediction.

Time enters CONCERT as a continuous attribute, so the model can predict the cells of each mouse at every time point by changing this attribute while keeping the basal state and position of each cell. This gives a predicted time course for each mouse that preserves its cell states and tissue structure. We scored inflammation with a panel of 17 DSS damage and regeneration markers from the original study [42]. The predicted course rose to a peak at day 9 and declined by day 21, as observed, and the ratios of predicted to observed daily means were 1.00, 0.91, 1.00 and 1.02 for days 0, 3, 9 and 21 (Fig. 5b). The predicted rate of change captured the switch from increasing to decreasing inflammation after day 9 (Fig. 5c). Because each mouse is measured once, such a per-mouse trajectory cannot be obtained by interpolating its own measurements.

We next examined the eight markers whose observed expression increased at least 1.5-fold from day 0 to day 9. For each marker, the predicted profiles agreed with the measured profiles across time points and individual mice (Fig. 5d). CONCERT also recovered the fraction of cells expressing each marker. With the detection probability computed under a Poisson approximation, *P* (count *>* 0) = 1 − *e*^−*µ*^, the predicted fractions differed from the observed fractions by 1.79 percentage points, whereas a quantile cutoff on the predicted mean gave an error of 8.58 percentage points (Fig. 5d). CONCERT thus recovers the expression level of individual inflammation markers and the fraction of cells that express them.

Predicted marker expression agreed with the measured profiles of each mouse in magnitude and spatial pattern. We used the predicted profiles to call a cell inflamed when its marker score exceeded a cutoff derived from healthy tissue, and mapped the inflamed regions of each section. The predicted fraction of inflamed cells followed the observed trend, low at early time points, highest at day 9 and lower by day 21 (Fig. 5e and Extended Data Fig. 3). Results for eight mice, two from each time point, are shown in Supplementary Fig. 12 to Supplementary Fig. 27.

### Responses to simulated lesions in ischemic stroke

We applied CONCERT to spatial transcriptomics of the mouse brain 3 days after photothrombosis, which induces a focal ischemic stroke in the cortex [43]. The dataset contains four coronal sections from a stroke mouse and four from a sham control mouse, collected at 1.2 mm intervals from 0.4 mm anterior to 3.2 mm posterior to bregma. We registered the sections of each mouse in a common in-plane frame by pairwise partial alignment with PASTE2 [52] and, because the sections are equally spaced, used the ordinal index of each section as the third coordinate (Extended Data Fig. 4a and Methods). All counterfactual predictions were made on the sham sections, which contain no lesion. CONCERT allows the user to specify the location and size of a lesion and predicts the response within and across sections. The genes analyzed here, Lgals1, Lcn2, and Spp1, are all associated with ischemic stroke [43, 53–56].

#### Lesion location and size

Photothrombosis offers limited control over the location and size of a lesion, so we placed simulated lesions on a sham section. The predicted response depended on location. For the same 200-spot lesion, CONCERT predicted a larger increase of the injury marker *Lcn2* in the cortex than in the thalamus (Extended Data Fig. 4b,c). When we enlarged a contiguous cortical lesion from 25 to 400 spots, predicted *Lcn2* expression within each lesion spot increased from 0.57 to 0.74 (Extended Data Fig. 4d,e). CONCERT thus predicts that a larger lesion produces a stronger response within each affected spot, as effects from neighboring lesion spots accumulate, and not only a larger affected area. CONCERT also predicted expression of the perilesional marker *Spp1* in a rim around the lesion and not in its core, as measured in the stroke brain (Extended Data Fig. 4f).

#### Lesions across serial sections

We next fitted a three-dimensional version of CONCERT, in which the perturbation field also uses the index of each section, so that a lesion effect can extend across sections (Supplementary Note 4). We examined *Lgals1*, one of the ischemia-induced genes whose expression decreases with distance from the infarct core in the original study [43]. Predicted *Lgals1* expression in the lesion region was highest in the first sham section, lower in the second and third and lowest in the fourth (Extended Data Fig. 4h), and the perilesional marker *Spp1* was concentrated around the lesion (Extended Data Fig. 4g), as observed in the stroke brain. The fractions of spots called as lesion from the predicted markers were 0.243, 0.188, 0.115 and 0.086 across the four sections, compared with 0.267, 0.208, 0.113 and 0.017 observed in the stroke brain. The predicted and observed fractions decreased from the first to the fourth section and agreed closely in the first three sections, whereas the predicted fraction in the fourth section was higher. In the stroke brain, *Lgals1* expression is higher at the border of the lesion than at its center, consistent with the accumulation of microglia and macrophages, the main sources of galectins, in the proximal peri-infarct area [43]. Predictions on intact sham tissue show *Lgals1* across the whole lesion region and do not resolve this difference between border and center. The 3D model predicted a stronger response than the 2D model, as expected when the effect can extend and accumulate across sections (Extended Data Fig. 4i).

## Discussion

We present CONCERT, a generative model for counterfactual prediction of perturbation responses in tissue. We evaluated CONCERT in within-niche prediction, which asks a model to predict a measured perturbation in a new niche, and in cross-slide prediction, which asks it to predict a perturbation on a slide where it was masked during training (Fig. 1a). In each setting, CONCERT predicted perturbation responses more accurately than perturbation models trained on dissociated cells, spatial perturbation models and baselines that average neighboring cells, with the largest margin in the ranking of perturbation-responsive genes. Benchmarks on cell-level SPAC-seq and spot-level Perturb-map data showed that this advantage holds across spatial resolutions.

CONCERT separates the basal state of a cell from the biological perturbation effect and from technical variation, and it separates the core perturbation effect from the part that varies with location. A perturbation-specific lengthscale sets the spatial range of the effect, and the niche gain adapts its strength to the surrounding cells. These components allow the same perturbation to produce different responses across tissue regions while sharing a common core effect. The learned parameters are also interpretable, as the niche gain and lengthscale of each knockout relate to the spatial distribution of its cells in tissue.

In Perturb-map lung tumors, CONCERT predicted how the response to the same knockout differs between tumor and normal niches and identified epithelial–mesenchymal transition as a program associated with niche composition. In DSS colitis, it reconstructed unmeasured time points for each mouse, and the predicted trajectories recovered the rise in injury to day 9 and its resolution by day 21, although each mouse was measured once [42]. In photothrombotic stroke, CONCERT predicted responses to lesions of chosen location and size, extended these predictions across serial brain sections in three dimensions and reproduced the expression patterns of the lesion core and perilesional tissue [43]. These studies show that CONCERT applies beyond tumor models and genetic perturbations, to chemical injury in the colon and ischemic injury in the brain.

This study focuses on perturbations observed during training. CONCERT predicts their effects in new cells, niches, locations and slides, but it is not designed to predict a perturbation that has never been measured, a setting addressed by models that use gene networks or other biological priors [22, 24]. Our evaluation therefore tests spatial transfer of known perturbations, including transfer to a slide on which the target perturbation was masked. All benchmarks use held-out cells, niches or slides within each dataset. Evaluation on two tumor models at different resolutions, together with applications in colon and brain, shows that CONCERT applies to several tissues, and larger spatial perturbation screens will allow evaluation across datasets.

CONCERT provides testable hypotheses about how a perturbation changes gene expression across tissue, but they do not directly predict functional outcomes, and future work could link them to cellular phenotypes and tissue-level measurements. Biological priors could extend the model to unseen perturbations, as drug structures, gene regulatory networks and pathway information have improved prediction in single-cell perturbation studies [22, 24, 57]. Histology and anatomical annotations offer another extension. CONCERT currently learns how responses vary across tissue from cell coordinates and gene expression alone, and tissue boundaries, lesion shapes and resection margins from histology, or drug penetration maps, could serve as additional inputs in tissues with complex structure. The stroke analysis used four sections spaced 1.2 mm apart, so the three-dimensional predictions were evaluated at few positions along the *z*-axis. Serial sections at finer spacing, or volumetric spatial transcriptomics, would allow CONCERT to model responses throughout the tissue volume and to test its three-dimensional predictions more completely.

CONCERT brings counterfactual perturbation prediction to spatial transcriptomics, where the response of a cell depends on its intrinsic state, its location and its niche. By modeling these dependencies, CONCERT predicts perturbation responses at tissue locations and under conditions that were not measured. These predictions can help prioritize spatial perturbation experiments and interventions that target specific tissue regions.

## Supporting information

all supp contents

## Data availability

The raw Perturb-map dataset is available via GEO under accession number GSE193460. The processed data is available via Figshare at https://figshare.com/articles/dataset/Datasets-Perturb-Map/29198468. The raw and processed SPAC-seq dataset is available in the original paper [41] from https://spac.pku-genomics.org. The processed mouse colitis MERFISH dataset used in the time-course case study is available from the original paper [42] from https://datadryad.org/dataset/doi:10.5061/dryad.rjdfn2zh3. The raw mouse brain stroke dataset is available from the authors of the original paper [43] upon request. The processed data is available via Figshare at https://figshare.com/articles/dataset/Mousebrain_stroke_dataset/29882900. The raw and processed Perturb-Fish dataset is available in the original paper [13] from https://doi.org/10.35077/ace-gem-get.

## Code availability

Project website is at http://zitniklab.hms.harvard.edu/projects/CONCERT. Python implementation of CONCERT is at https://github.com/mims-harvard/CONCERT.

## Acknowledgements

We thank Yuchang Su and Yepeng Huang for helpful discussions on the manuscript. We thank Yang Li for designing the CONCERT logo and Ruthie Johnson for assistance with figure design and generation.

## Authors contribution

X.L. retrieved, processed, and analyzed the spatial perturbation datasets. X.L. implemented, benchmarked CONCERT, and together with Z.K. B.F., and S.M. performed detailed analyses of CONCERT. X.L., M.L., and M.Z. designed the study with M.Z. leading the overall research. All authors discussed the research and contributed to the manuscript.

## Methods

We first describe the datasets used for benchmarking and for biological applications. We then define prediction problems, describe the architecture, objective and training of CONCERT, and explain how the model performs counterfactual prediction and imputation. The final sections describe the benchmark design and evaluation, the analyses for each application and the statistical tests.

### Datasets

#### SPAC-seq datasets

We evaluated CONCERT primarily on SPAC-seq, a spatial CRISPR screen assay that captures sgRNA identity *in situ* alongside the whole transcriptome [41]. We used the *in vivo* screen in which 1,520 sgRNAs were introduced into MC38-Cas9 cells and five mouse replicates were profiled, yielding five tissue slides with 632,032, 454,825, 304,478, 356,519 and 453,638 spots, respectively. Each spot is assigned by its captured sgRNA to one of three classes, a gene knockout, a non-targeting control, or unassigned when no sgRNA is recovered. In total, this dataset contains 2,201,492 spots over 20,579 genes, of which 472,667 are gene-knockout spots, 4,625 are non-targeting control spots and 1,724,200 are unassigned, and 546 knockout targets are supported by at least 25 spots. We retained the 2,000 highly variable genes shared across the five slides as the model input. Because this dataset covers hundreds of perturbations measured in the same tissue, we used it to evaluate CONCERT across many perturbations, which the three perturbations in Perturb-map do not allow.

#### Perturb-map datasets

We also used the Perturb-map data of mouse lung adenocarcinoma (GEO GSE193460), which comprise four slides. Spot annotations were obtained in the original study by imaging cytometry [12]. The GSM5808054 and GSM5808055 samples contain 1,903 and 1,872 spots, respectively, and each carries two genetic perturbations, *Jak2* knockout (*Jak2*-KO) and *Tgfbr2*-KO in GSM5808054, and *Ifngr2*-KO and *Tgfbr2*-KO in GSM5808055. The GSM5808056 and GSM5808057 samples contain 1,233 and 1,355 spots, respectively, and each carries one perturbation, *Ifngr2*-KO and *Tgfbr2*-KO. All slides also have annotated unperturbed tumor spots, tumor periphery spots and normal spots. We preprocessed the data downloaded from GEO with Seurat [58] and converted them into H5 or H5AD files as input for the models. In total, this dataset contains 6,363 spots, with 21 *Jak2*-KO spots, 224 *Tgfbr2*-KO spots, 105 *Ifngr2*-KO spots and 6,013 unperturbed spots.

#### Mouse colitis MERFISH datasets

To study a perturbation whose response develops over time, we used the MERFISH atlas of colitis from Cadinu et al. [42]. The authors induced colitis in mice with dextran sodium sulfate (DSS) and collected colon tissue at day 0 (healthy), day 3, day 9 and day 21, so that the same injury is observed at four points along its course. The processed data contain 1,348,582 cells from 52 tissue slices of 15 mice over a 942-gene panel, with 239,450 cells from three mice (10 slices) at day 0, 299,274 cells from four mice (13 slices) at day 3, 660,977 cells from six mice (25 slices) at day 9 and 148,881 cells from two mice (4 slices) at day 21. Cell-type annotations at three levels of granularity and neighborhood labels are provided in the original study. This dataset resolves single cells and replicates each time point across several mice, which allows us to test whether CONCERT recovers the time course of a perturbation and not only a single snapshot.

#### Mouse stroke datasets

We used the spatial transcriptomics dataset of mouse ischemic stroke from Han et al. [43]. This dataset includes brain sections from two mice, one subjected to photothrombosis (PT) to induce ischemic stroke in the cortex and one sham control without PT. For each mouse, four coronal sections were collected at 1.2 mm intervals, from 0.4 mm anterior to 3.2 mm posterior to bregma [43]. The PT sections contain 2,397, 2,650, 2,781 and 2,755 spots, and the sham sections contain 1,752, 2,031, 2,618 and 2,793 spots, respectively. To construct a 3D coordinate system, we registered the four stroke sections into a common in-plane frame by pair-wise partial alignment with PASTE2, applied the same procedure independently to the four sham sections, and used the ordinal index of each section as the third coordinate, which is proportional to its anteroposterior position because the sections are equally spaced. In total, this dataset contains 10,583 PT spots and 9,194 sham spots. Of the PT spots, 511 lie in the ischemic core and 1,303 in the perilesional region, and the remaining spots are largely unaffected by PT.

#### Perturb-FISH dataset

We used the tumor section of Perturb-FISH, an image-based CRISPR screen that reads out sgRNA identity and a targeted transcriptome in the same cells [13]. The section contains 9,432 cells measured with the 500-gene ImmunOncology MERFISH panel. Each cell has either a knockout of one of 35 Toll-like receptor and NF-*κ*B pathway genes or the control label, which covers 4,264 cells, and 31 knockout targets are observed in at least 25 cells. Cell centroids are rescaled so that one coordinate unit corresponds to one cell spacing. Because the panel covers a limited number of genes, we used this dataset only to replicate the parameter interpretation analysis performed on SPAC-seq.

For all datasets, we normalized the count matrices with SCANPY [59] before model training.

### CONCERT model

#### Model overview

CONCERT predicts the response of a cell to a perturbation from its gene expression profile, spatial coordinates and perturbation state, and it can also impute missing cells and predict their responses. The model has three modules. The disentanglement module separates spatial perturbation transcriptomics (SPT) data in latent space along two axes. Along the spatial axis, it separates the smooth anatomical variation of the tissue from non-spatial variation. Along the perturbation axis, it separates the perturbation effect from the perturbation-free basal state and splits this effect into a core component, shared wherever the perturbation is present, and a spatial adjustment that varies with location. The perturbation-signal module adapts the effect to the niche of each cell and separates it from the technical shifts caused by guide delivery and by variation between slides. The generative module decodes the basal state and applies these shifts to reconstruct gene expression.

For a cell *k*, the attribute embeddings map the expression profile **x**^*k*^ to a basal embedding 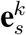 and the attribute labels to a perturbation embedding 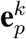 and one embedding for each further attribute. The basal state is built from two separate inputs. The basal spatial field maps the coordinates **s**^*k*^ to the spatial basal latent **z**^sp^ without reading expression, and the non-spatial encoder maps the non-perturbation embedding 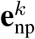, which concatenates 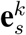 with the embeddings of further attributes, to the non-spatial basal latent **z**^G^ without reading coordinates. The decoder maps **z** = ⊕ (**z**^sp^, **z**^G^) to the perturbation-free mean ***µ***_basal_. The perturbation follows a separate path. The perturbation field combines 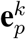 with **s**^*k*^ into the gene-space shift Δ_*p*_(**s**^*k*^) of Eq. (19), the perturbation-signal module adapts this shift to the niche of the cell, giving 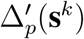, and the predicted post-perturbation mean is ***µ***_basal_ multiplied by the exponentiated shift in Eq. (31). Because only this path receives the perturbation, a perturbation can be applied to a cell without changing its basal state. We first state the problem formulation, then describe the modules and the training objective, and finally explain counterfactual prediction, imputation and the implementation for each dataset.

#### Preliminaries

Uppercase **X** denotes a matrix and lowercase **x** denotes a vector, and uppercase *X* denotes a set of observations or variables. We denote *p*(**X**) as a probability distribution over a set of random variables **X** and *p*(**X** = **x**) as the probability of **X** that is equal to the value of **x** under the distribution *p*(**X**). This section uses terminology and concepts from variational inference [60]. Neural networks are written *f*_•_, learned or fixed linear maps **W**_•_ and **b**_•_, and loss terms *L*_•_ with weights *λ*_•_, with the subscript naming the component. Latent variables and embeddings are named along two axes, basal or perturbation, and spatial or non-spatial. A name that carries only one of the two, or neither, refers to the combined quantity, so the spatial basal latent and the non-spatial basal latent are the two halves of the basal latent.

#### Problem formulation: Disentangling perturbation effects

The input consists of three quantities per cell. The gene expression profile is **X** ∈ ℝ^*N×G*^ for *N* cells and *G* genes, and the coordinates in tissue space are **S** ∈ ℝ^*N×D*^, with *D* = 2 for a single tissue section and *D* = 3 when serial sections are registered into a stack. The perturbation state is the categorical label **l**_**p**_ ∈ *{*1, …, *N*_*p*_*}* over *N*_*p*_ perturbations 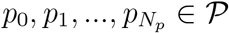, of which *p*_0_ is the state without perturbation. Further cell attributes **V** can be supplied, categorical such as disease state or continuous such as time since perturbation (Supplementary Note 10). The goal is to disentangle **X** into **E**_**s**_, the embedding of the cell state independent of the perturbation (the basal state), and **E**_**p**_, the embedding of the perturbation effects,

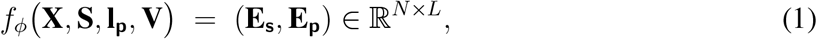

where *f*_*ϕ*_ is a function with learnable parameters *ϕ* and *L* is the dimension of the embeddings. A function *f*_*θ*_ with learnable parameters *θ* reconstructs the gene expression profiles from the disentangled embeddings,

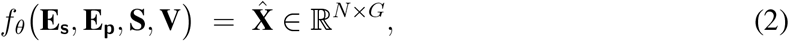

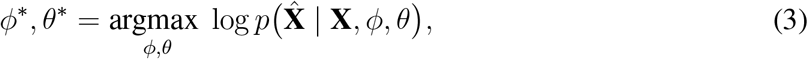

where *p* is an appropriate density function. This objective ensures that **E**_**s**_ and **E**_**p**_ preserve the information in **X** needed for reconstruction. Each attribute in **V** is disentangled from **E**_**s**_ in the same way and has its own embedding, so that it can be changed independently.

We split each embedding further, so that a perturbation effect can be placed in tissue space. Along the *spatial axis*, the basal state is split into a spatial component, a function of the coordinates alone, and a non-spatial component,

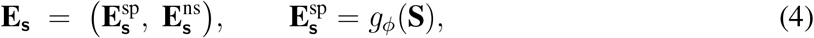

which separates the smooth anatomical variation of the tissue from the variation intrinsic to a cell. 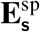 depends on **S** and not on the measured expression of any neighbors. Along the same axis, the perturbation embedding is split into a *core* effect, shared at every location where the perturbation is present, and a *peripheral* term, the spatial adjustment that varies with location,

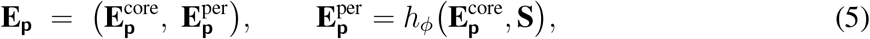

where the peripheral term is confined to a neighborhood of the perturbed cells whose extent is learned for each perturbation. The same perturbation can therefore reach a different distance, and act with a different magnitude or sign, in different parts of the tissue. This second split distinguishes CONCERT from non-spatial perturbation models. Setting 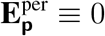 leaves an effect that is identical wherever the perturbation is present, as in a model without spatial context. The maps *g*_*ϕ*_ and *h*_*ϕ*_ are implemented by the basal spatial field *f*_sp_ and the perturbation field *f*_field_, defined below.

#### Problem formulation: Spatial perturbation prediction

Given a set *C* ⊆ *{*1, …, *N}*, |*C*| = *M*, of cells without perturbation (**l**_**p**_ is *p*_0_), we obtain their disentangled embeddings 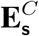 and 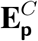. With *P* the set of perturbations observed in **X**, the goal is to predict the counterfactual response gene expression of the cells in *C* to a perturbation *p*_1_ ∈ *P*,

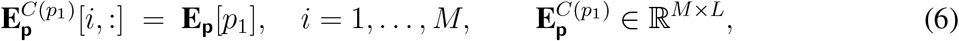

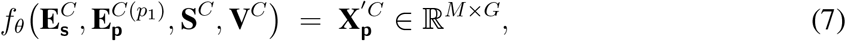

where every cell in *C* receives the row of the learned codebook **E**_**p**_ that belongs to *p*_1_, and the indicators that switch on the guide term and the perturbation shift are set to those of *p*_1_. 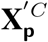 is the predicted post-perturbation expression of the cells in *C*, conditioned on their coordinates **S**^*C*^ and attributes **V**^*C*^. The core and peripheral terms are read from the embedding of *p*_1_, but only the core term is the same at every location, and the peripheral term is evaluated at **S**^*C*^. Applying *p*_1_ to two cells with the same basal state at different locations therefore gives different predictions.

#### Problem formulation: Cell imputation with perturbation prediction

A set of observed cells has the embedding **E** = [**E**_**s**_||**E**_**p**_], where || denotes row-wise concatenation. A further set *U* holds *M*_*u*_ unseen cells whose expression profiles are missing and whose spatial coordinates 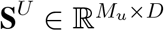 are known. A function *f*_*π*_ first imputes the embeddings of the missing cells,

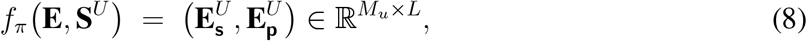

where 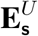 and 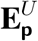 are the imputed embeddings of the cells in *U* . We then perturb 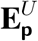 *in silico* with a perturbation *p*_1_ ∈ *P*,

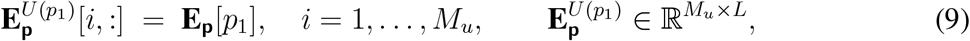

and generate the perturbed gene expression profiles 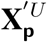 of the cells in *U*,

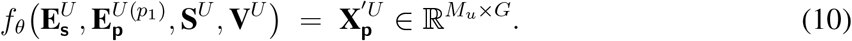

#### Attribute embeddings

CONCERT disentangles gene expression profiles in latent space following LORD, a method for latent optimization for representation disentanglement [61], which BioLORD applied to single-cell perturbation data [23]. CONCERT decomposes each profile into embeddings of cell attributes, which can be categorical, such as perturbation or disease state, or continuous, such as time after perturbation or tumor size. A regularized basal embedding captures the remaining cell-specific heterogeneity. In CONCERT, this embedding is amortized, that is, an encoder produces it from the expression profile of the cell, instead of storing it as a free parameter for each training cell as in BioLORD. Variation between slides has no attribute embedding and is handled in the decoder (Eq. (31)). CONCERT applies to cell-level and spot-level data. We describe the model for single cells, as in the SPAC-seq and MERFISH datasets, and for spot-based assays such as Perturb-map each data entry is a spot.

We use SPAC-seq as the running example, in which the encoded attribute is the knockout target read out from the captured sgRNA of each cell. The attributes encoded for each dataset are listed in Supplementary Note 10. The basal embedding of cell *k* is amortized from its log-normalized expression profile **x**^*k*^ by an MLP 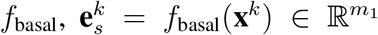. Each categorical attribute has a randomly initialized codebook, including the perturbation codebook 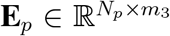 over 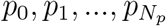, where *p*_0_ is the state without perturbation and *N*_*p*_ is the number of perturbation states, and, for a further categorical attribute with *N*_*d*_ levels, a codebook 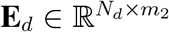. A continuous attribute **c** is projected to dimension *m*_4_ by an MLP 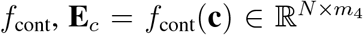, which is how time enters the colitis model (Supplementary Note 3). The dimensions *m*_1_, *m*_2_ and *m*_3_ are set to 256 by default. As in CPA [20], *f*_basal_ learns the heterogeneity of gene expression that is independent of the other attributes. The basal embedding 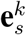 differs from the basal state **E**_**s**_ of the problem formulation. The encoder infers the non-spatial component 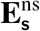 from 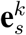, the basal spatial field produces the spatial component 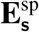 from the coordinates, and neither component contains the perturbation.

To prevent 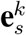 from learning information that belongs to the perturbation embedding 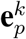 or to the embeddings of other attributes, we regularize it with Gaussian noise,

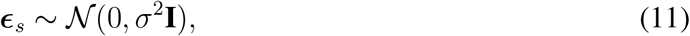

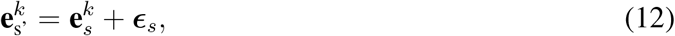

with *σ* = 0.1 for all datasets. The embeddings of cell *k* are concatenated into its latent representation (Fig. 1b),

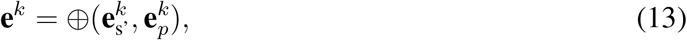

where ⊕ denotes row-wise concatenation. Because 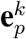 is a known attribute embedding and 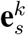 is inferred, the concatenation contains the information needed to reconstruct the spatial perturbation transcriptomic data after training. We write **e** for **e**^*k*^ where the cell is clear from context.

#### Basal state disentanglement

The basal state of a cell has a spatial component **z**^sp^, a function of the location of the cell, and a non-spatial component **z**^G^, each produced by a separate path. Variational autoencoders (VAEs) [62] assume that latent representations are independent and identically distributed across samples. The GP-VAE [63–65] relaxes this assumption with a Gaussian process (GP) prior that models correlations between latent variables. For input data **x ∈ X**, the latent distribution *z*(**x**) follows

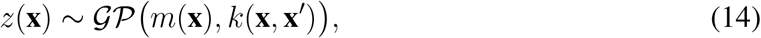

where *m*(**x**) = *f*^*µ*^(**x**) and *k*(**x, x**^*′*^) = *f*^*σ*^(**x**) *f*^*σ*^(**x**^*′*^)^⊤^, *f*^*µ*^ and *f*^*σ*^ are neural network encoders for the mean and variance, and **x** and **x**^*′*^ are two cells in **X**. spaVAE [26] applied the GP-VAE to spatially resolved transcriptomics, with kernel values computed from the spatial locations of cells. CONCERT follows this principle for the spatial component of its basal latent and approximates the GP with random Fourier features, which learns smooth dependencies between cells without a fixed number of neighbors.

#### Spatial basal component

The basal spatial field maps coordinates directly to the spatial basal latent. For a cell at **s** ∈ ℝ^2^, we draw a fixed random projection **W**_rff_ ∈ ℝ^2*×F*^ and offset **b**_rff_ ∈ ℝ^*F*^ once at initialization and form random Fourier features [44] of the rescaled coordinates,

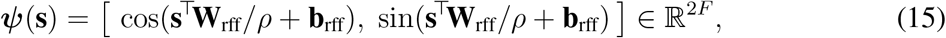

where *ρ* rescales coordinates to a common range across slides and the bandwidth of **W**_rff_ sets the spatial resolution of the features. The features pass through a shared MLP *f*_sp_ and a meanfield variational linear head [66] that returns the mean of **z**^sp^, with a learned variance for each latent dimension. Random Fourier features with a Bayesian linear output approximate a GP with a stationary kernel [44], and the MLP between them makes the kernel a learned function of these features. Because **W**_rff_ and **b**_rff_ are fixed, the basal spatial field has no kernel scale that can shrink during training, so the spatial component cannot collapse to a constant. The variational head regularizes the weights of this mapping and contributes a closed-form KL term.

#### Non-spatial basal component

MLP encoders 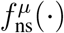 and 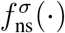 map the non-perturbation embedding **e**_np_ of a cell to the mean and variance of the non-spatial basal latent **z**^G^,

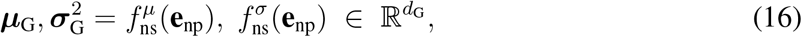

where **z**^G^ is the non-spatial component 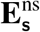 of the basal state and follows a standard normal prior,

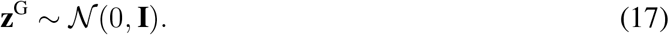

The encoder reads no coordinates and the basal spatial field reads no expression, so each half of the basal latent has one source, and the approximate variational posterior factorizes as

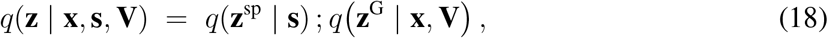

where **s** is the spatial location of a cell and **V** the cell attributes other than the perturbation. The latent dimensions are *d*_sp_ for **z**^sp^ and *d*_G_ for **z**^G^, and the corresponding matrices over all cells are 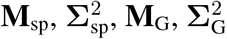 and **Z**.

#### Perturbation effect disentanglement

Unlike the basal state, the perturbation effect acts in gene space, as a shift on the log scale of the perturbation-free mean. We write Δ_*p*_(**s**) ∈ ℝ^*G*^ for the shift that perturbation *p* imposes at location **s**, Eq. (19), and 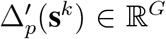 for the same shift after the niche of cell *k* has modulated and gated it, Eq. (22). 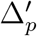 enters the decoder in Eq. (31). Both shifts are zero for a cell without a gene-targeting guide.

For perturbation *p*, the non-spatial component is a shift *f*_core_(**e**_*p*_) ∈ ℝ^*G*^, a linear readout of the perturbation embedding **e**_*p*_ that is the same at every location. The spatial component is a residual conditioned on coordinates, so that the same perturbation can act differently in different regions of the tissue. It depends on position through *M*_*r*_ *reference locations* 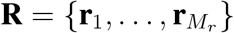 spread across the rescaled coordinate frame, initialized on a regular grid and free to move during training. A cell at **s** is described by its proximity to each reference location, 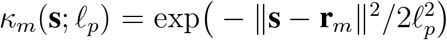, collected into a vector 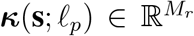, where the lengthscale ℓ_*p*_ sets how far the influence of a reference location reaches and is learned for each perturbation. The two components are read from the same perturbation embedding and sum to the perturbation shift at that location,

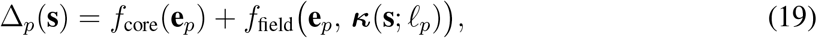

where *f*_field_ is an MLP shared across perturbations. The residual is initialized near zero, so training starts from a spatially uniform effect and departs from it only where a spatial component improves the fit.

The lengthscale makes the spatial extent of each perturbation interpretable. A single linear head shared across perturbations predicts it from the perturbation embedding,

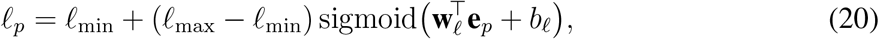

where **w**_ℓ_ and *b*_ℓ_ are the weight vector and bias of the head, and ℓ_min_ and ℓ_max_ are fixed bounds on the range of distances an effect can span. This construction has two purposes. Because sigmoid( ·) lies between 0 and 1, Eq. (20) always returns a value between ℓ_min_ and ℓ_max_, and a squared-hinge penalty on the same interval in the objective provides a second guard. Because **w**_ℓ_ and *b*_ℓ_ are shared across perturbations, perturbations with similar embeddings receive similar lengthscales, so a knockout observed in few cells inherits its spatial extent from the perturbations it resembles instead of being fitted in isolation. The exact form of each field and all hyperparameter values are given in Supplementary Note 1.

#### Constraints enforcing the disentanglement

Three constraints separate the perturbation effect from the basal state. First, the parameterization is asymmetric. The perturbation embedding is shared, one row of **E**_*p*_ for all cells with the same perturbation, while the basal path is a function of the profile of each cell, so any variation that co-varies with perturbation identity is represented once in **E**_*p*_ at a far lower cost than relearning it for each cell. Second, the capacity of the basal path is limited, because the Gaussian noise of Eq. (12) and the *L*_2_ term of *L*_latent_ penalize a basal representation that stores attribute-specific detail. Third, the perturbation embedding is withheld from the variational encoder. The latent **z** therefore contains no perturbation information by construction, and a perturbation can reach gene expression only through the niche-modulated shift 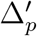 of Eq. (22), which enters the decoded mean in Eq. (31).

#### Perturbation-signal module: niche-specific learning and noise correction

This module modulates the magnitude of the effect separately from its spatial extent. For a cell *k*, we collect the set *N*_*k*_ of its *k*_nn_ nearest cells in tissue space within the training batch and summarize their non-spatial basal latents **z**^G^ into a niche descriptor 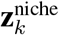,

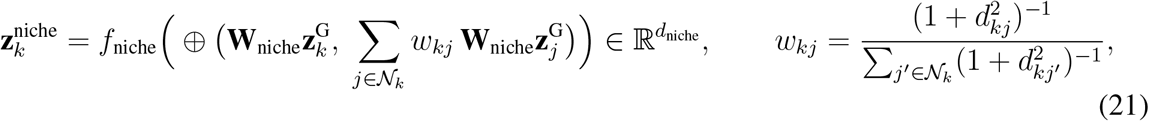

where *d*_*kj*_ is the distance between cells *k* and *j*, **W**_niche_ is a linear projection and *f*_niche_ is a two-layer MLP. The descriptor depends on the states of the surrounding cells and on their distances, so it can be evaluated on a slide held out from training. Neighbors are taken within the batch, so one distance computation per batch replaces a precomputed neighborhood graph, and **z**^G^ is detached, so the descriptor acts only as a conditioning signal. The descriptor turns Δ_*p*_(**s**^*k*^) into the modulated shift 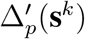 through a signed scalar gain *γ*_*k*_, a per-gene residual ***β***_*k*_ and a scalar gate *a*_*k*_,

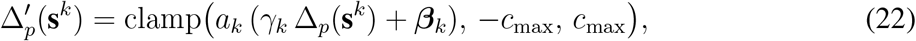

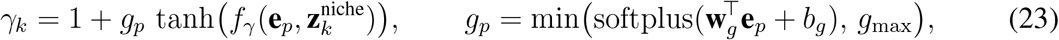

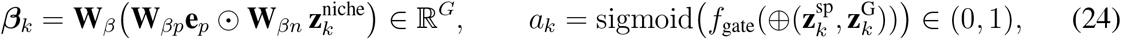

where *f*_*γ*_ and *f*_gate_ are MLPs, ⊙ is the elementwise product, and **W**_*β*_, **W**_*βp*_ and **W**_*βn*_ are the factors of a rank-*r*_*β*_ bilinear map. The gain is a feature-wise linear modulation (FiLM) layer [45] with a bounded depth *g*_*p*_, the niche gain of each perturbation. A head shared across perturbations reads *g*_*p*_ from the perturbation embedding, so a rarely observed perturbation inherits it from the perturbations it resembles. Because *γ*_*k*_ can cross zero once *g*_*p*_ exceeds one, the same perturbation can act with a different strength, or in the opposite direction, depending on its niche. The residual ***β***_*k*_ adds a niche-specific term for each gene, and the gate *a*_*k*_, read from both halves of the basal latent, sets how much of the shift reaches a given cell. Every head is initialized so that the module starts at the identity, 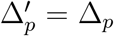, with zero weight and bias in the tanh head, *b*_*g*_ = −4, **W**_*β*_ = 0, and zero weight and bias 3 in the last layer of *f*_gate_. These four values were not tuned against the reported metrics. All heads are fitted jointly with the rest of the model through the reconstruction term and the log-fold-change term *L*_LFC_ of *L*_pert_, and no niche label, clustering or annotation is supplied. The penalty *L*_*g*_, also part of *L*_pert_, acts on the mean gain and the squared entries of **W**_*β*_, so a departure from the identity is kept only where it reduces the reconstruction and log-fold-change terms. The shift is clamped to [ −*c*_max_, *c*_max_] with *c*_max_ = 5. Values of *k*_nn_, *d*_niche_, *r*_*β*_, *g*_max_ and the widths of *f*_*γ*_, *f*_gate_ and *f*_niche_ are listed in Supplementary Note 1.

The second part of this module separates the biological response from technical noise. Cells are grouped by guide status into cells with no assigned guide, cells with a non-targeting control guide and cells whose guide targets a gene. A per-gene guide term ***µ***_guide_ ∈ ℝ^*G*^ applies to every cell that received a guide, and the perturbation shift 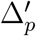 applies only to cells whose guide targets a gene. The three classes make the two terms identifiable, because unassigned cells fix the perturbation-free baseline, non-targeting controls identify the shift produced by guide delivery alone, and targeting guides identify 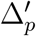 on top of this shift. The same classes also route the basal path. Only unassigned cells have a free non-spatial posterior, and the non-spatial basal latent of every cell that received a guide is set to its prior. The basal state of a perturbed cell is therefore given by the spatial field at its coordinates alone, and its perturbed expression cannot enter **z**^G^. A learned bias 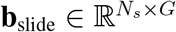, with one row per slide and 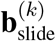 the row of the slide of cell *k*, absorbs residual variation between the *N*_*s*_ sections. ***µ***_guide_ and **b**_slide_ are free parameters initialized at zero, each fitted through the reconstruction term from the cells its indicator selects, and ***µ***_guide_ is clamped to [−5, 5]. Only models trained on more than one section include **b**_slide_. The guide term, the slide bias and the perturbation shift act as shifts on the log scale of the perturbation-free mean 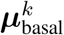, as defined in Eq. (31).

Guide delivery differs between SPAC-seq sections, and most sections contain few non-targeting controls, so the guide shift also has a slide-specific component. In the multi-section SPAC-seq models, a cell with a guide on slide *s* receives 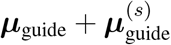, where 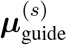 is the row of slide *s* in a free matrix of shape *N*_*s*_ *× G*, initialized at zero and clamped to [− 5, 5]. It is fitted from all cells with a guide on that slide, so knockout cells inform the delivery shift on sections where non-targeting controls are sparse. A prediction on slide *s* applies the rows of **b**_slide_ and 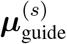 of that slide, as in training. The other multi-section models generate predictions in a common frame that does not depend on the section.

#### Clonality gate

In a pooled *in vivo* screen, a guide label marks a cell as part of a transduced clone. On sections with sparse transduction, cells in small clones lie apart from other cells with a guide, and their expression reflects a weaker share of the response than cells in large clones. The SPAC-seq models therefore weight each cell with a guide by clone size. A clone is the set of cells with one perturbation on one slide, and all non-targeting control cells of a slide form a single clone. Cells of clones with at least *n*_clone_ = 100 training cells keep weight one, and cells of smaller clones receive weight zero in the reconstruction likelihood and in the log-fold-change targets of *L*_pert_. Unassigned cells always keep weight one, and a knockout keeps the signal of its large clones on the other slides.

#### Generative module: decoder for reconstructing post-perturbation gene expression

This module decodes the basal latent into the perturbation-free mean ***µ***_basal_ and applies the perturbation-signal terms to it. The spatial and non-spatial basal latents are concatenated into the basal latent, with posterior

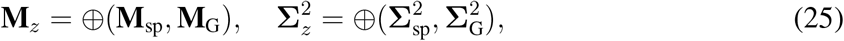

where **M**_sp_ collects the means returned by the basal spatial field at the coordinates of each cell and 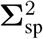 its learned variance for each dimension, shared across cells, and **M**_*z*_ and 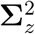 are the combined posterior mean and variance. The latent distribution is

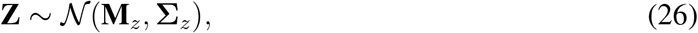

and we use the reparameterization trick to backpropagate through it, drawing latent samples for a cell *k* as

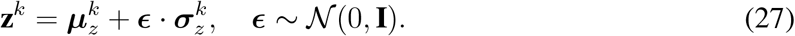

We draw *n*_*z*_ samples, with *n*_*z*_ = 1 during training and *n*_*z*_ = 10 at inference for stable predictions. Each sample **z**^*k*^ passes through the MLP decoder *f*_dec_,

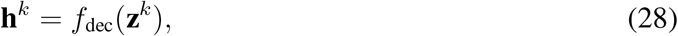

and a mean head returns the *perturbation-free* mean of cell *k*, while a separate parameter gives the dispersion of the negative binomial (NB) distribution,

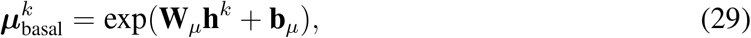

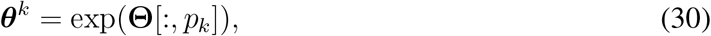

where **W**_*µ*_ and **b**_*µ*_ are the weights and bias of the mean head, and 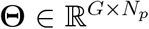 is a free parameter matrix with one dispersion vector per perturbation, from which the column of the perturbation *p*_*k*_ of cell *k* is selected. The dispersion is therefore learned for each gene and perturbation and does not depend on **h**^*k*^. Because the perturbation embedding is withheld from the encoder, Eq. (29) gives the profile that cell *k* would show without a guide. The niche-modulated shift 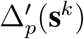 of Eq. (22), the guide shift ***µ***_guide_ and the slide bias 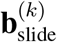 enter the model only here, each switched on by the indicators of cell *k*,

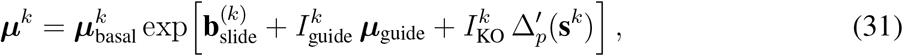

where 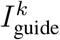 and 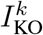 indicate whether cell *k* has a guide and whether that guide targets a gene, and ***µ***_guide_ includes the slide row 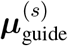 in the SPAC-seq models. For datasets without non-targeting controls, we disable the guide term. The mean of the negative binomial distribution of cell *k* is the shifted ***µ***^*k*^ of Eq. (31), not 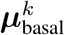, and the dispersion ***θ***^*k*^ does not depend on the shift.

At prediction, the shifted mean averaged over the *n*_*z*_ latent samples, 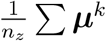, is the predicted response gene expression of cell *k*. Setting 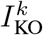 to zero in Eq. (31) gives the prediction for the same cell without the perturbation, and the two predictions give the log-fold change reported by counterfactual queries. In the SPAC-seq benchmark, the predicted mean is converted to counts at the measured depth of the predicted cell, the same step applied to every compared method (Evaluation).

#### Reconstruction loss

For each cell *k*, the negative binomial loss uses the raw counts 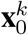, the predicted mean ***µ***^*k*^ of Eq. (31), the dispersion ***θ***^*k*^ and the size factor 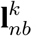,

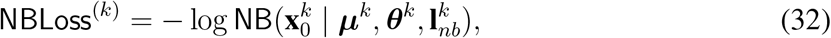

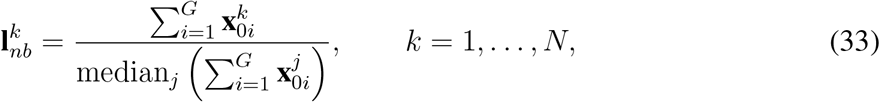

where *G* is the number of genes in **X**_0_ and *j* indexes all cells. The reconstruction loss is summed over the genes and the cells of a mini-batch *B*,

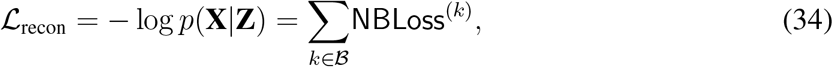

and measures the discrepancy between observed and reconstructed data.

#### Overall training objective

CONCERT is trained by minimizing a single objective with three parts. The first measures how well the decoded mean matches the observed counts, the second limits the capacity of the latent variables, and the third constrains where the perturbation effect is placed,

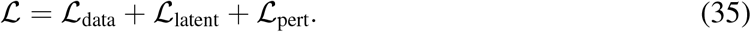

The data term *L*_data_ combines the negative binomial reconstruction loss of Eq. (34) with a mean squared error between the predicted mean ***µ***^*k*^ and the raw counts of the same cell. The latent term *L*_latent_ penalizes capacity. It contains the KL divergence of the non-spatial basal latent, the closed-form KL divergence of the variational head of the basal spatial field, and an *L*_2_ penalty on the basal embedding. Because the features of the basal spatial field are deterministic, its variational head carries the uncertainty of the spatial basal latent. The *L*_2_ penalty prevents the basal embedding from absorbing information that belongs to the attribute embeddings. A free-bits floor allows each latent dimension a fixed budget of information without penalty, and the weight of the two KL divergences is increased from zero over the first epochs, so that reconstruction is established before the latents are compressed.

The perturbation term *L* _pert_ acts on the shift instead of the reconstruction and supervises 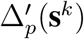 directly on the quantity we evaluate. Knockout cells are grouped by niche and perturbation, the group mean of 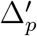 is compared with the measured log-fold change between knockout and control cells of that group by squared error, and the result is averaged over groups. Because ***µ***_guide_ is shared with the non-targeting controls and cancels in this contrast, the group mean of 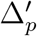 equals the predicted log-fold change of the model, so the supervision reaches the perturbation embedding, the lengthscale head and the niche heads together. The same term penalizes spatial autocorrelation of the non-spatial basal latent **z**^G^ through a Moran’s *I* statistic, so that the smooth spatial structure of the tissue is represented by the spatial path. Its pair weights are Gaussian, 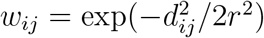, so only autocorrelation at about the scale *r* of the perturbation effects is penalized and finer cell-type structure is preserved. Finally, *L*_pert_ keeps each lengthscale within its bounds by a squared hinge and pulls the niche gain and the modulation weights toward the identity, so that the model represents dependence on the niche only where it improves the fit. The KL weight follows the warm-up schedule, and the weights of the other contributions are given with their explicit forms, the free-bits floor and the warm-up schedule in Supplementary Note 2. Module hyperparameters are listed in Supplementary Note 1.

#### Counterfactual prediction and imputation for observed and unobserved cells

As shown in Fig. 1c, CONCERT performs *in silico* perturbation and places the perturbation effect across tissue space through the peripheral term of the perturbation embedding, without passing messages between cells. Continuing with SPAC-seq as the example, an unperturbed cell *k* has the embedding 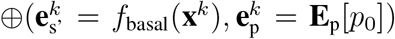. If cell *k* is observed on the slide, we replace its embedding of the unperturbed state *p*_0_ with the embedding of the target state of perturbation 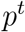,

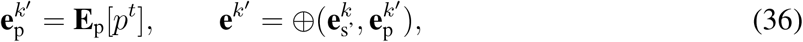

and the basal embedding and the embeddings of any further attributes are unchanged.

If cell *k* is not observed on the slide, its expression profile is missing, so its non-spatial basal latent cannot be encoded and is taken from neighboring cells. If the target perturbation *p*^*t*^ of cell *k* is known, we assign its embedding from **E**_p_ as in Eq. (36) and collect the neighborhood *N* (*k*) of cells with *p*^*t*^. If we only impute the expression of cell *k* without perturbation, so that none of its attributes is known, *N* (*k*) contains the neighboring cells regardless of their perturbation state. In both cases, the non-spatial basal latent of *k* is the mean posterior over *N* (*k*),

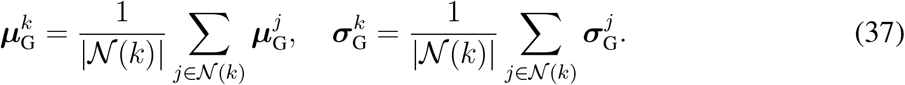

The spatial basal latent needs no averaging. The basal spatial field maps coordinates to **z**^sp^ and has no free parameters for individual cells, so it is evaluated directly at the coordinates **s**^*k*^ of the unobserved cell,

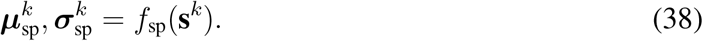

Because the spatial component of the basal state is a function of the coordinates alone, it is defined at any location, including locations where expression was not measured. The spatial and non-spatial basal latents of cell *k* are concatenated and decoded into its imputed expression.

#### Multi-slide coordinate frame

Multi-slide input is placed in one coordinate frame. The coordinates of each slide are rescaled to the target range, and the slides are tiled into disjoint regions separated by a gap, so that no cell has a neighbor on another slide while the field and kernel parameters are shared across slides. The per-slide bias **b**_slide_ of Eq. (31), created only when more than one slide is present, absorbs residual technical differences between sections. CONCERT assumes that the spatial extent of a perturbation is the same on every slide once this bias is applied, so the lengthscale of Eq. (20) is learned for each perturbation and not for each slide.

#### Parameterization and training

The embeddings of the cell attributes have 256 dimensions by default. The mini-batch size is 2,048. We use Adam [67] with a learning rate of 1 *×* 10^−4^ and a weight decay of 1 *×* 10^−6^, and the learning rate of the kernel scale parameters is ten times higher so that they converge within the same training budget. For the sequencing-based datasets (SPAC-seq, Perturb-map and mouse stroke), we selected the 2,000 most highly variable genes for model training, and for the imaging-based datasets, we used the full gene panels (942 genes for colitis and 500 genes for Perturb-FISH). The MLP encoder and decoder have layer sizes [128, 64] and [128], respectively. The spatial and non-spatial basal latents have 2 and 8 dimensions by default, so the basal latent **z** has 10 dimensions. The basal spatial field uses 128 random Fourier features at bandwidth 0.15 with a hidden width of 128, and the perturbation field uses 16 reference locations with a hidden width of 256. The KL weight (*λ*_KL_) increases linearly from 0 to 1 over the first 20 epochs. Five percent of the training set is held out for validation and used for early stopping, with a patience of 200 epochs and a minimum improvement of 0. For the large multi-slide datasets, we require the validation loss to improve by at least 0.05 before the patience counter is reset and reduce the patience to 50 epochs. All experiments were run on NVIDIA A100 GPUs.

#### Implementation on SPAC-seq data

We train CONCERT on the five SPAC-seq slides jointly, encoding the knockout target in the disentanglement module, and define niches with a 6 × 6 grid on each slide. Knockout targets observed in fewer than 25 cells are excluded, which removes 147 targets and leaves 546 knockout targets for training. Each perturbation has its own lengthscale, so its effect can extend to a different distance in tissue, which makes the learned kernel parameters interpretable for each perturbation (Supplementary Note 5).

#### Implementation on Perturb-map data

On the Perturb-map slides, we encode the perturbation (*Jak2*-KO, *Tgfbr2*-KO, *Ifngr2*-KO or control) and the cell type (tumor, normal or periphery), and define niches with a 4 *×* 4 grid on each slide. We run the counterfactual prediction tasks on this dataset, as well as a joint analysis in which a contiguous knockout neighborhood is deconvolved and imputed. The full list of encoded attributes and the patch sampling used for the counterfactual tasks are given in Supplementary Note 6,7,10.

#### Implementation on the colitis time course

We train CONCERT on all 52 slices from the 15 mice, with perturbation identity (DSS or control) and day (0, 3, 9 and 21) as attributes, and impute the spatial transcriptomic profile of each mouse and each slice at the unmeasured days. Because day enters as a continuous attribute, the model can be queried at days that were not sampled. Supplementary Note 3 describes the kernel that models variation over time.

#### Implementation on mouse stroke PT data

We train CONCERT on eight sections, encoding perturbation identity and the tissue attribute. The raw coordinate frames of the eight sections are not comparable, so before any three-dimensional model is fitted, we register them into a common frame by pairwise partial alignment with PASTE2 [52] (Supplementary Note 4). The lesion is defined as the infarct core together with the perilesional area, using the region labels of the original study [43]. We perform counterfactual prediction of ischemic regions in 2D, within each section, and in 3D, across sections.

### Benchmarking methods

We consider four types of competing methods to benchmark CONCERT:

1. **Reference strategies**: Four simple strategies that predict the perturbed profile of a spot without learning a model. *Mean target* takes the mean expression of 50 measured spots drawn at random from the target perturbation state, a number on the scale of the perturbed spots each model receives in training. *Mean niche* takes the mean expression of the spots in the same niche. *Spatial kNN* and *GEX kNN* average the *k* nearest measured spots in the target state, the first selecting neighbors by spatial distance and the second by transcriptomic distance. These models are not performance ceilings. They average over spots, track the whole gene expression well, but suppress the perturbation-specific response.
2. **Non-spatial perturbation models**: scGEN [19], Biolord [23], GEARS [22] and STATE [46]. These methods predict a perturbation response from expression alone and have no access to where a spot sits in the tissue, so comparing against them demonstrates how much of CONCERT’s performance comes from spatial context.
3. **Spatial perturbation models**: CelComen [37], CellAgentChat [40] and Steamboat [47]. These methods are context-aware perturbation prediction model, and are therefore the closest published comparators to CONCERT. CelComen learns two gene-by-gene interaction matrices, one acting within a cell and one between neighbouring cells, and simulates a knockout by setting the target gene to zero and propagating that change through the graph. Each matrix carries one weight per ordered pair of genes, so the model is fitted on a panel of 500 genes, and a knockout target outside that panel cannot be perturbed. CellAgentChat is an agent-based model of cell-cell communication built on a ligand-receptor database, in which an *in-silico* perturbation blocks a ligand or a receptor. Only targets listed in that database can be perturbed, and most of the knockout targets in the SPAC-seq screen are not listed. Steamboat is an attention-based factor model that reconstructs each cell from itself and from its spatial neighbourhood. It carries no perturbation term of its own, so the perturbed state has to be supplied by substituting measured cells that already carry the target knockout.
4. **Spatial imputation and deconvolution models**: Tangram [27], SpatialScope [50] and SpaIM [49] are used to recover gene expression at locations where it was not measured, which is what the patch imputation task asks for. They are used as comparators for this task only. For deconvolution, the comparator is non-negative least squares over empirical signatures (NNLS).

All published methods are implemented following their official documentation, with all parameters kept at their default values.

#### STATE fine-tuning for cross-slide prediction

STATE has two components [46]. A State Embedding model (SE) maps each cell to an embedding, and a State Transition model (ST) maps a set of control cells to their perturbed state. The released ST checkpoints are trained on human cell lines. The perturbation vocabulary of the ST-SE-Replogle checkpoint covers only a few perturbations that we plan to predict. We therefore fine-tune STATE on the SPAC-seq training spots through the pretrain-then-fine-tune route of the official implementation. Each spot is embedded with the pretrained SE-600M model, whose weights are kept fixed. The ST model is initialized from the ST-SE-Replogle checkpoint and fine-tuned on the same training spots as every other method.

### Evaluation

The cells carrying the target state, against which a prediction is scored, are selected differently in the two tasks. In within-niche prediction, we sample patches of unperturbed cells for prediction at locations where measured cells of the target perturbation are present in the same local niche, so that the comparison is against what that perturbation actually does in that microenvironment. In cross-slide prediction, the cells of the target perturbation are removed from the predicted slide before training, serving as the ground truth. The prediction is scored against the masked cells on this slide, so the ground truth lies in the same niche with the predicted cells, but the perturbations are learned from a different slide. The log-fold change is always taken between the perturbed and unperturbed cells within each slide, so that slide-level differences are canceled. In both tasks, predictions are made only on the masked unperturbed cells.

We score a prediction by how well it reproduces the response a perturbation induces, rather than the absolute expression level. This design is essential because a model may achieve strong overall performance simply by reproducing basal gene expression profiles, even if it entirely fails to capture perturbation-driven signals. For a given perturbation, we therefore summarize both the predicted and the measured profiles as a log-fold change (LFC) against the unperturbed control spots,

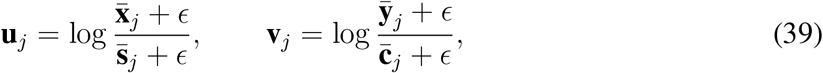

where 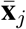 is the mean predicted expression of gene *j* over the predicted cells, 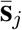 the mean measured expression of the same cells before perturbation, 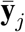 the mean measured expression of the cells carrying the target perturbation, 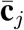 the mean measured expression of unperturbed control cells disjoint from the predicted cells, *ϵ* = 10^−6^, and *j* runs over the *g* evaluated genes. Expression is log-normalized to 10,000 counts per cell. The predicted response is thus the change a method applies to the measured cells it is given, and every method is scored against the same measured cells. The evaluated genes are the highly variable genes of the slide that fall in the 2,000-gene panel of CONCERT, excluding the knocked-out gene, and they are identical for all methods.

Measured expression is a sparse count, and a model that returns an expected expression profile differs from it in noise character alone. On SPAC-seq, every method’s prediction for a cell is therefore renormalized to a relative profile, drawn once as Poisson counts at that cell’s measured library size, and log-normalized like the measurements. The same transform is applied to all methods. We then report two complementary metrics on these LFC vectors:

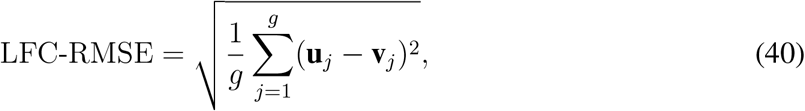

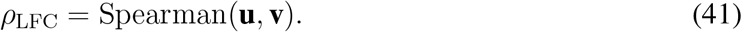

LFC-RMSE measures how close the predicted response is to the measured one in magnitude, and the Spearman correlation *ρ*_LFC_ measures whether the model ranks the responding genes in the right order. We report both metrics to achieve a comprehensive evaluation. An averaging strategy can achieve a competitive LFC-RMSE while its *ρ*_LFC_ is near zero, the failure mode a magnitude-only metric would hide. Both metrics are computed per evaluation record, averaged over the records of each knockout (within-niche) or knockout–target-slide pair (cross-slide), and then averaged over knockouts or pairs; the error we report is the standard error of the mean across knockouts or pairs. Each method is averaged over the knockouts it scored.

We summarize each benchmark condition with three further views computed over all scored knockouts. For the per-knockout scale, the scores of the methods that scored a knockout are rescaled to [0, 1] inside that knockout, with 1 on the best and 0 on the worst method, so that easy and hard knockouts weigh the same. For the rank view, the methods that scored a knockout are ranked inside it, with rank 1 the best, and a method’s ranks are averaged over its knockouts. For the margin view, we take the difference between CONCERT and each competing method on every knockout both scored, signed so that a positive value favours CONCERT.

### Within-niche and cross-slide prediction on SPAC-seq

We conduct these tasks on SPAC-seq data, which measures hundreds of knockouts on the same tissue. We evaluated CONCERT in two settings: within-niche prediction and cross-slide prediction. In both settings, the model is given a cell’s location, its measured unperturbed expression and a target perturbation. It then predicts that cell’s perturbed expression. Predictions are always made on unperturbed cells and scored against the measured perturbed cells. The settings differ only in what is withheld during training. In within-niche prediction, spatial patches of unperturbed cells are excluded from training. The model predicts their response to each of 87 perturbations, using measured cells of the target perturbation that lie in the same local niche. In cross-slide prediction, the perturbed cells of a perturbation are masked on one target slide while that perturbation remains observed on the donor slides. The model must predict the response on the unperturbed cells from the target slide. 85 knockout–target-slide pairs are held out this way. We note that cross-slide prediction tests spatial transfer, not unseen perturbation prediction: the perturbation is still observed in the training data but in a different slide.

#### Definition of a niche

A niche is a fixed spatial unit, not a transcriptomic cluster. This ensures that the same partition applies to every method and every slide. We rescale the coordinates of each slide per axis to a common frame and partition each slide into a regular 6 *×* 6 grid. This gives 36 niches per slide, each covering one sixth of the slide extent along each axis. The grid is an evaluation construct. It defines the local neighbourhood, the niche, over which log-fold changes are computed, so that a prediction is always compared with the measured cells of the target perturbation in the same part of the tissue. In the same way, each Perturb-map slide is divided into a regular 4 *×* 4 grid since the smaller spot numbers.

#### Held-out patches within a niche

Within each niche, we cluster the unperturbed cells by *k*-means on their coordinates into *k* = min(4, max(2, *n/*15)) contiguous sub-patches, where *n* is the number of unperturbed cells in the niche. Niches with fewer than 40 unperturbed cells are not split. Sub-patches, drawn at random under a fixed seed, are withheld from training for every method. The unselected patches in the niche provide the observed reference against which log-fold changes are taken. This reference is therefore disjoint from the cells being predicted.

#### Filtering perturbations by cell number

For both tasks, we only consider perturbations that appear in sufficient numbers of cells to estimate the target state reliably. A knockout captured in only a handful of cells yields a noisy reference against which no perturbation signals can be captured. For within-niche prediction, we evaluate a niche only if its withheld patch contains at least 20 unperturbed cells. Similarly, the most abundant perturbation in that niche must be carried by at least 20 cells. Across the five slides, this admits 34, 25, 24, 23 and 24 niches and 25, 20, 19, 19 and 17 perturbations respectively, giving 100 perturbations-slide pairs over 87 distinct perturbations. For cross-slide prediction, a knockout must be carried by at least 20 cells on the donor slide 1-1. This slide is always used as the donor slide, because it contains most perturbations and perturbed cells, ensuring enough training sample size for each perturbation. The perturbation must also be carried by at least 100 cells on the target slide, ensuring a reliable reference for evaluation. Every knockout meeting both thresholds is evaluated, giving 85 knockout–target-slide pairs: 33 on slide 1-2, 19 on 1-3, 16 on 1-4 and 17 on 1-5. Every method predicts the same 5,000 unperturbed cells per target slide, and the measured control cells exclude them. We apply the same thresholds to every method, so all methods are scored on the same perturbations, the same niches and the same cells.

### Patch imputation and counterfactual prediction

Spatial transcriptomics slides contain regions where cells are missing because of technical dropout or physical damage to the tissue. When such a region lies within a perturbed area, a model that fills the gap must also predict the perturbed state of the missing cells. We tested this with a masking experiment on SPAC-seq slide 2-2, which comes from a different donor than the slides used in the benchmark. Before training, we removed half of the cells of each of the five largest knockout patches on the slide (H2-Ea, Iqgap1, Cttn, Ccn1 and Phgr1). We trained CONCERT on the remaining cells only, so no masked cell contributed to any embedding. At inference, we used CONCERT to reconstruct the masked cells at their original coordinates, one knockout patch at a time, and compared the predictions with the measured profiles of the masked cells. Because the model never observed a cell of the masked knockout at those locations, reconstructing a masked patch requires imputation and counterfactual prediction together. CONCERT infers the basal state of the masked cells from their spatial neighborhood and the perturbation effect from cells of the same knockout elsewhere on the slide.

We compared CONCERT with three spatial imputation methods, Tangram, SpatialScope and SpaIM, which impute missing expression but do not model perturbations. To keep the comparison fair, we scored all methods on the same masked cells and on the genes shared by their panels. We report two metrics. The first is the mean per-cell Spearman correlation between imputed and measured expression. The second is the Spearman correlation between the imputed and measured log fold change of each patch, where the measured log fold change is the mean expression of the knockout cells in the patch minus the mean expression of nearby unassigned cells on the same slide. For the imputation methods, we computed the imputed log fold change against the same nearby unassigned cells. For CONCERT, we computed it against the predicted unperturbed expression of the same cells.

### Pseudo-bulk deconvolution with known perturbation composition

To test whether CONCERT can resolve a measurement that mixes several cells, we constructed pseudo-bulk spots from SPAC-seq slide 2-1, for which the perturbation identity of every cell is known. We partitioned the slide into spatially compact groups of about nine cells with a greedy nearest-neighbor procedure and averaged the counts of the cells in each group. This yielded 34,921 spots from 273,886 cells, with a mean of 7.8 cells per spot. Spots at the edge of a knockout patch mixed knockout cells with the surrounding unperturbed cells, and the perturbation composition of every spot, defined as the fraction of its cells in each perturbation state, was known by construction. We evaluated the ten knockouts that formed the largest patches on the slide.

We estimated perturbation composition of each spot with non-negative least squares (NNLS), the standard approach for deconvolving a mixed measurement, which models the expression profile of a spot as a non-negative weighted sum of reference signatures. As a baseline, we used empirical signatures, defined as the average expression profile of the measured cells in each state. We then repeated NNLS with the location-specific signatures of CONCERT, which are the expression profiles that CONCERT predicts for each state at the location of the spot. The two settings differed only in the signatures, so the comparison isolated the contribution of the signatures learned by CONCERT.

We scored the deconvolution against ground truth in two ways. First, we compared the estimated composition with the true fraction of each state in each spot, using the mean absolute error and the Pearson correlation. Second, we compared the predicted signatures with the measured profiles of the individual cells that made up each spot, which tested whether the deconvolution recovers the expression of the constituent cells in addition to their proportions.

### Temporal imputation on the colitis time course

The colitis MERFISH data follow one perturbation, DSS-induced injury, across four time points (days 0, 3, 9 and 21). Each mouse was sampled at a single time point for spatial profiling, so the data contain no longitudinal measurements from the same animal, and the response of a given tissue region over time cannot be observed directly. We used CONCERT to impute the unmeasured time points for each mouse and each tissue section and to reconstruct a complete time course from cross-sectional data.

We trained one model on all 52 sections from 15 mice. The model was used to predict the expression of the cells of each section at the time points at which that mouse was not sampled, so that every mouse has a predicted profile at all four time points and not only at the day on which it was sampled. Time enters the model as a continuous attribute, which allows predictions at any day. Because no ground truth exists at days that were not sampled in the cohort, we restricted all predictions and evaluations to the four sampled time points.

### Interpretation of learned kernel parameters

The niche gain *g*_*p*_ is a scalar learned for each perturbation *p* by the niche-conditioned modulation head (Supplementary Note 1). It is computed from a linear projection of the perturbation embedding **e**_*p*_, passed through a softplus function and capped at *g*_max_ = 3, so that *g*_*p*_ = min(softplus 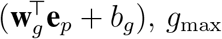). The head applies a multiplicative gate *γ* = 1 + *g*_*p*_ tanh(· ) to the perturbation shift, where the argument of tanh depends on the niche embedding of the receiving cell. Because tanh takes values between −1 and 1, the gate lies between 1 − *g*_*p*_ and 1 + *g*_*p*_, and *g*_*p*_ sets how strongly the niche can modulate the predicted response. A perturbation with *g*_*p*_ near zero is predicted to have the same response wherever it occurs, whereas a perturbation with a large *g*_*p*_ can have a response that changes with the composition of the surrounding neighborhood.

We extracted *g*_*p*_ from the trained SPAC-seq model in the within-niche benchmark, together with the footprint lengthscale ℓ_*p*_, which sets how far the spatial effect field of the same perturbation extends. We tested whether these two parameters track the spatial distribution of the perturbed cells in tissue. We described the distribution of each perturbation with layout descriptors computed from the measured cell coordinates. These descriptors are the fraction of the section area reached by the perturbed cells, the nearest-neighbor index, which is the mean distance from a perturbed cell to the nearest cell carrying the same perturbation divided by the distance expected under random placement of the same number of cells, and four descriptors of focus structure obtained by spatial clustering. We related each parameter to each descriptor with partial Spearman correlations controlled for the number of perturbed cells and the norm of the perturbation embedding (Statistical analyses). We repeated the analysis on a model trained in the same configuration on a Perturb-FISH tumor section, using the 31 knockouts carried by at least 25 cells. Extended Data Fig. 5 shows the results, and Supplementary Note 5 gives the descriptor definitions, the correlation values and the corrected *p*-values.

### Sensitivity analyses

We assessed the sensitivity of CONCERT to ten hyperparameters of its spatial components by varying one hyperparameter at a time and holding all others at their reported values. Four hyperparameters control the perturbation footprint field: the range of its lengthscale, the number of inducing points, the scale of the spatial coordinates and the weight of the log-fold-change penalty. Four control the spatial smoothness penalty and the basal spatial field: the weight of the penalty, the radius at which it is evaluated, the number of random features and the bandwidth of the field. Two control the niche-conditioned modulation head: the dimension of the niche code and the size of the neighborhood from which it is computed. We trained three models with different random seeds for each setting and scored each model by the rank correlation and the RMSE of predicted against measured log fold changes. We report the mean across the three seeds with the standard error across seeds. To set the scale of variation expected from random initialization alone, we also trained five models of the reported configuration that differ only in their random seed. Supplementary Note 9 lists the tested values and describes the results, which are shown in Figs. Supplementary Fig. 3–Supplementary Fig. 8.

### Statistical analyses

In the benchmarking experiments, we compute each metric for every evaluation record, average it over the records of each knockout in within-niche prediction or of each knockout–target-slide pair in cross-slide prediction, and report the mean across knockouts or pairs. The uncertainty on each metric is the standard error of the mean across knockouts or pairs. We apply the same procedure to every method and average each method over the knockouts or pairs for which it returns predictions. We compare CONCERT with each competing method using a two-sided paired Wilcoxon signed-rank test. This nonparametric test asks whether the paired differences between two methods are centered at zero, without assuming that the differences follow a normal distribution. We pair the two methods on matched evaluation units. A unit is one (perturbation, slide, niche) record scored by the two methods, so the comparison always uses the same held-out cells and the same measured response. Some methods return predictions for only part of the perturbation panel, and we test these methods on the units they share with CONCERT. We test the rank correlation of log fold changes and the log-fold-change RMSE separately.

Each table of comparisons contains one test for each competing method and metric. We correct the *p*-values within a table using the Benjamini-Hochberg procedure, which controls the false discovery rate, the expected fraction of false positives among the comparisons called significant, at 0.05. For each comparison, we report the adjusted *p*-value, the mean of each metric for the two methods, and the fraction of matched units on which CONCERT is closer to the measured response, that is, has a higher rank correlation or a lower RMSE than the competing method.

To relate the learned spatial parameters to the observed spatial layout of each perturbation, we compute partial Spearman correlations between each learned parameter, the niche gain and the lengthscale, and each layout descriptor of the perturbed cells, such as the fraction of the section that they reach and the nearest-neighbor index. A partial correlation measures the association between two quantities after removing the variation that covariates explain in each of them. We use the number of perturbed cells and the norm of the perturbation embedding as covariates, because the spread of a perturbation in tissue and the size of its learned parameters can increase with the number of cells that carry it. We report 95% confidence intervals computed on the Fisher scale and correct the *p*-values across descriptors using the Benjamini-Hochberg procedure at a false discovery rate of 0.05.

In the pathway analysis shown in Fig. 4g, we test which biological programs in the predicted response change with the composition of the niche. We place each knockout in silico into 40 niches sampled across the slide, rank genes by how their predicted response changes with the tumor content of the niche after removing differences in the overall magnitude of the response, and run gene set enrichment analysis (GSEA) on this ranking against the Hallmark collection of MSigDB [68]. GSEA tests whether the genes of a set concentrate at the top or the bottom of the ranking. We report normalized enrichment scores, which adjust the enrichment score for the size of the gene set, with *q*-values corrected using the Benjamini-Hochberg procedure, and we consider a gene set significant at *q <* 0.05.

## Extended Data Figures

**Extended Data Fig. 1:**
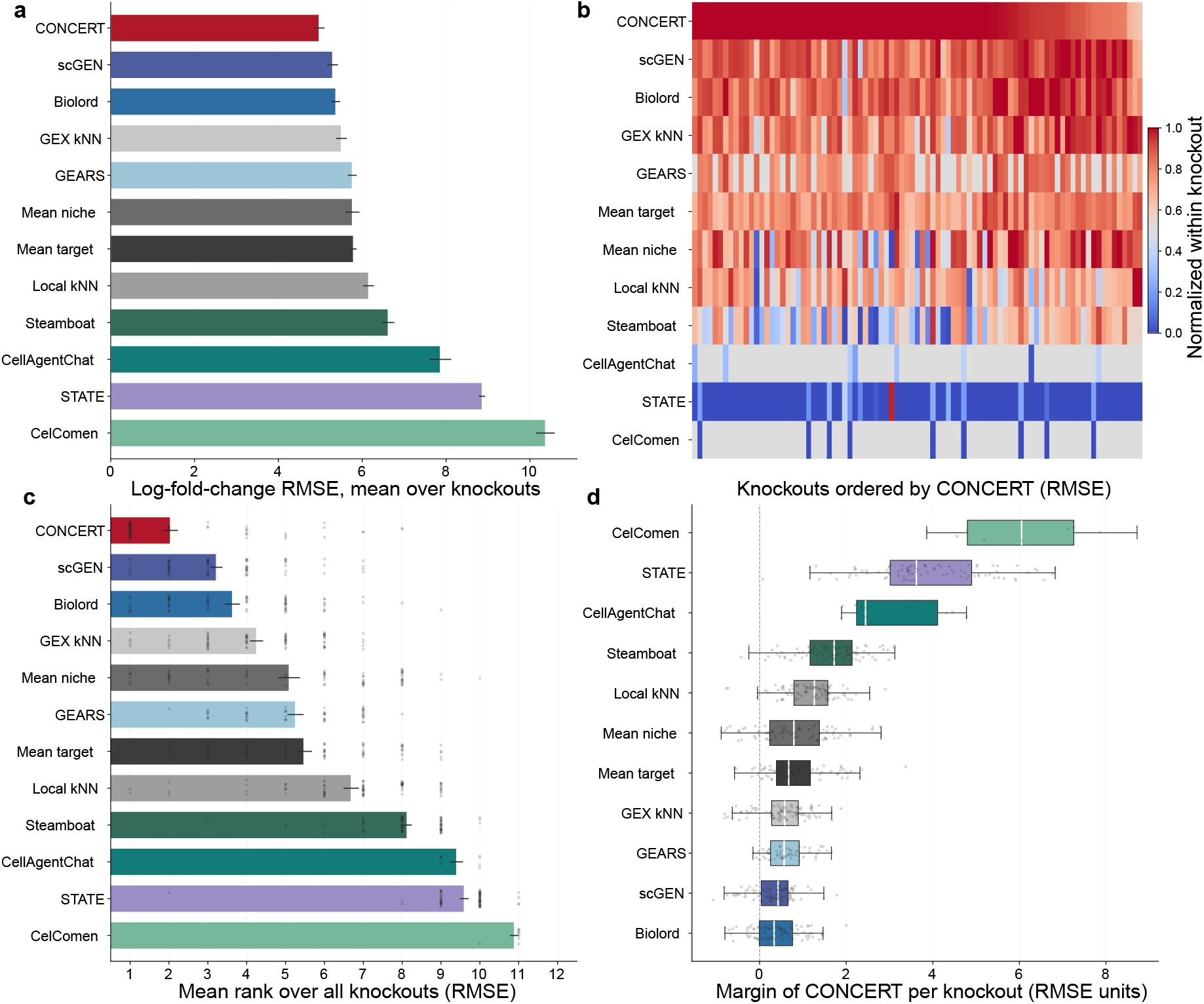
Response magnitude in within-niche prediction. Companion to Fig. 2, which reports the gene ordering of the same predictions. Accuracy is the root-mean-square error between the predicted and the measured log-fold change against unperturbed control cells; lower is better. **a**, RMSE of each method, the mean over the 87 knockouts with the standard error across them. **b**, Every knockout as one column, rescaled inside the column so that 1 marks the best method and 0 the worst; grey marks a knockout a method left unpredicted. **c**, Mean rank of a method inside a knockout, taken over the methods that scored it, with rank 1 the best and one dot per knockout. **d**, Margin of CONCERT over each competing method on every knockout both scored, positive where CONCERT leads. Method colors are as in Fig. 2.

**Extended Data Fig. 2:**
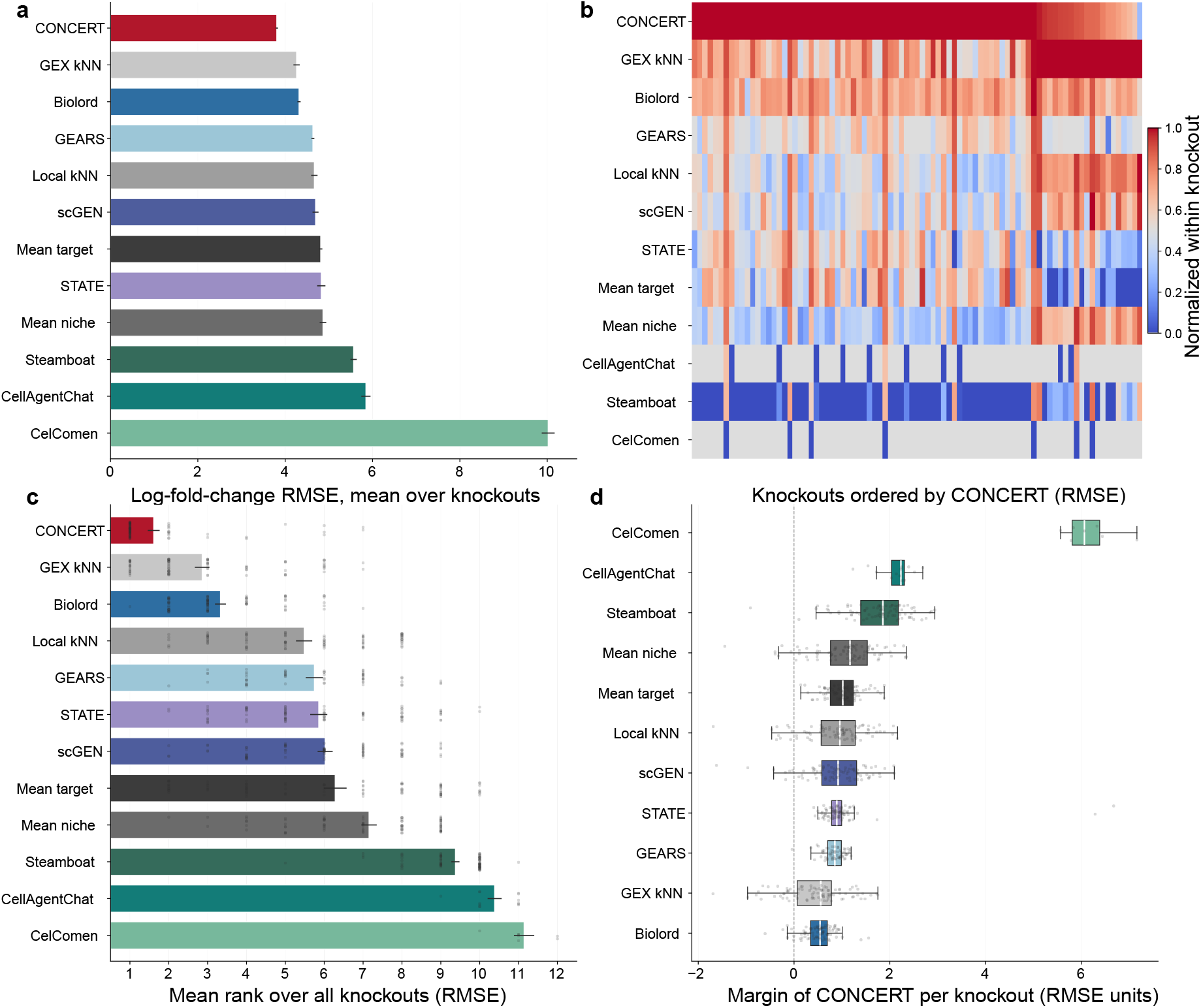
Response magnitude in cross-slide prediction. Companion to Fig. 3, which reports the gene ordering of the same predictions, and scored as in Extended Data Fig. 1. **a**, RMSE of each method, the mean over the 85 knockout– target-slide pairs with the standard error across them. **b**, Every pair as one column, rescaled inside the column so that 1 marks the best method and 0 the worst; grey marks a pair a method left unpredicted. **c**, Mean rank of a method inside a pair, taken over the methods that scored it, with rank 1 the best and one dot per pair. **d**, Margin of CONCERT over each competing method on every pair both scored, positive where CONCERT leads. Method colors are as in Fig. 2.

**Extended Data Fig. 3:**
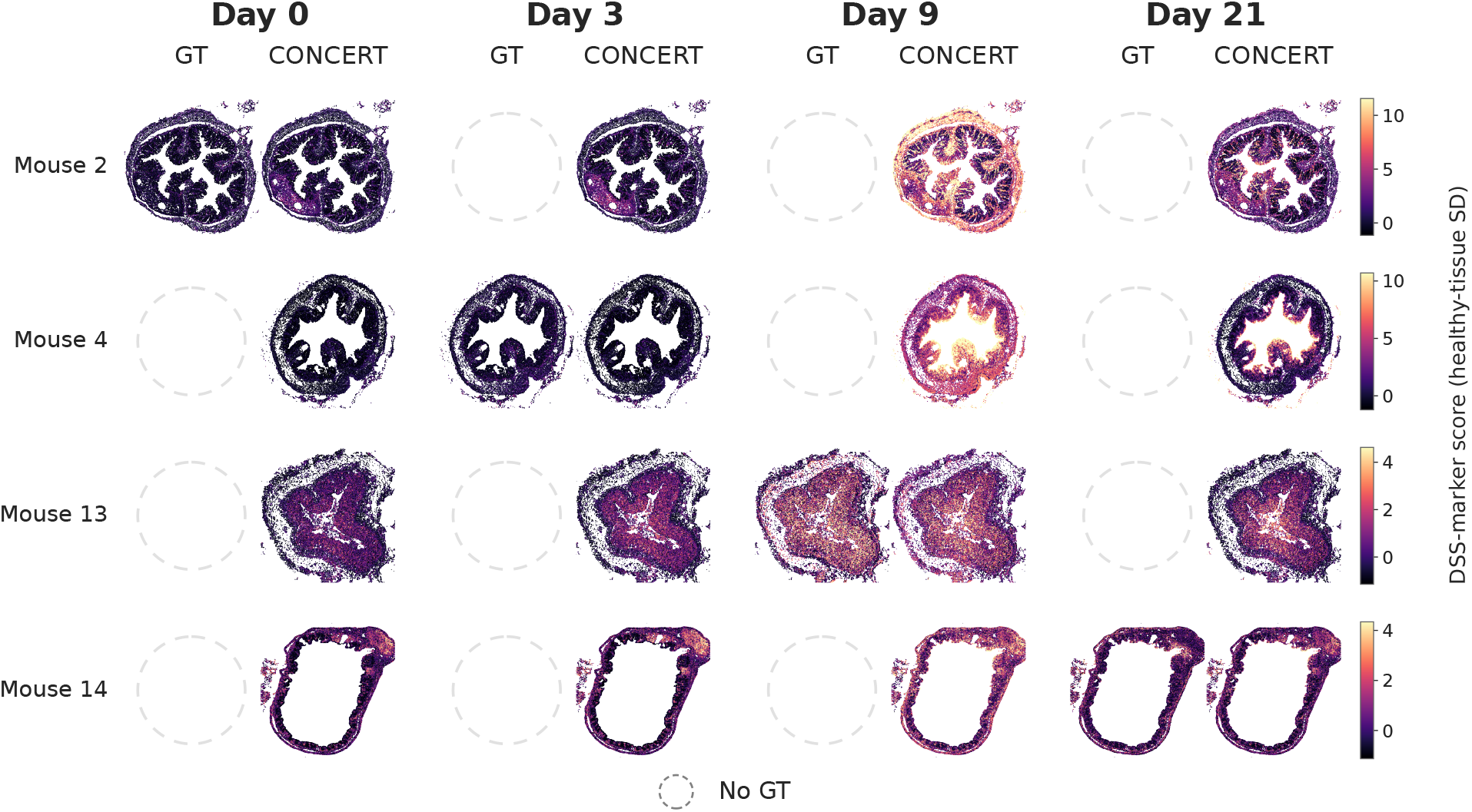
Reconstructed inflammation field of individual animals at days other than their sampling day. Companion to Fig. 5e, which thresholds this field into inflamed regions. Rows are four representative mice from the DSS colitis MERFISH atlas [42], one per sampled day, each shown as its largest colon section. Sections were chosen for structural completeness alone. Columns are the four virtual days, each split into observed ground truth (GT) and CONCERT. Color is the DSS-marker score in healthy-tissue s.d. units, scaled per row, so panels are comparable within each row. Because sampling is terminal, GT exists only on the diagonal; every other GT slot is drawn as an empty dashed circle beside the CONCERT prediction.

**Extended Data Fig. 4:**
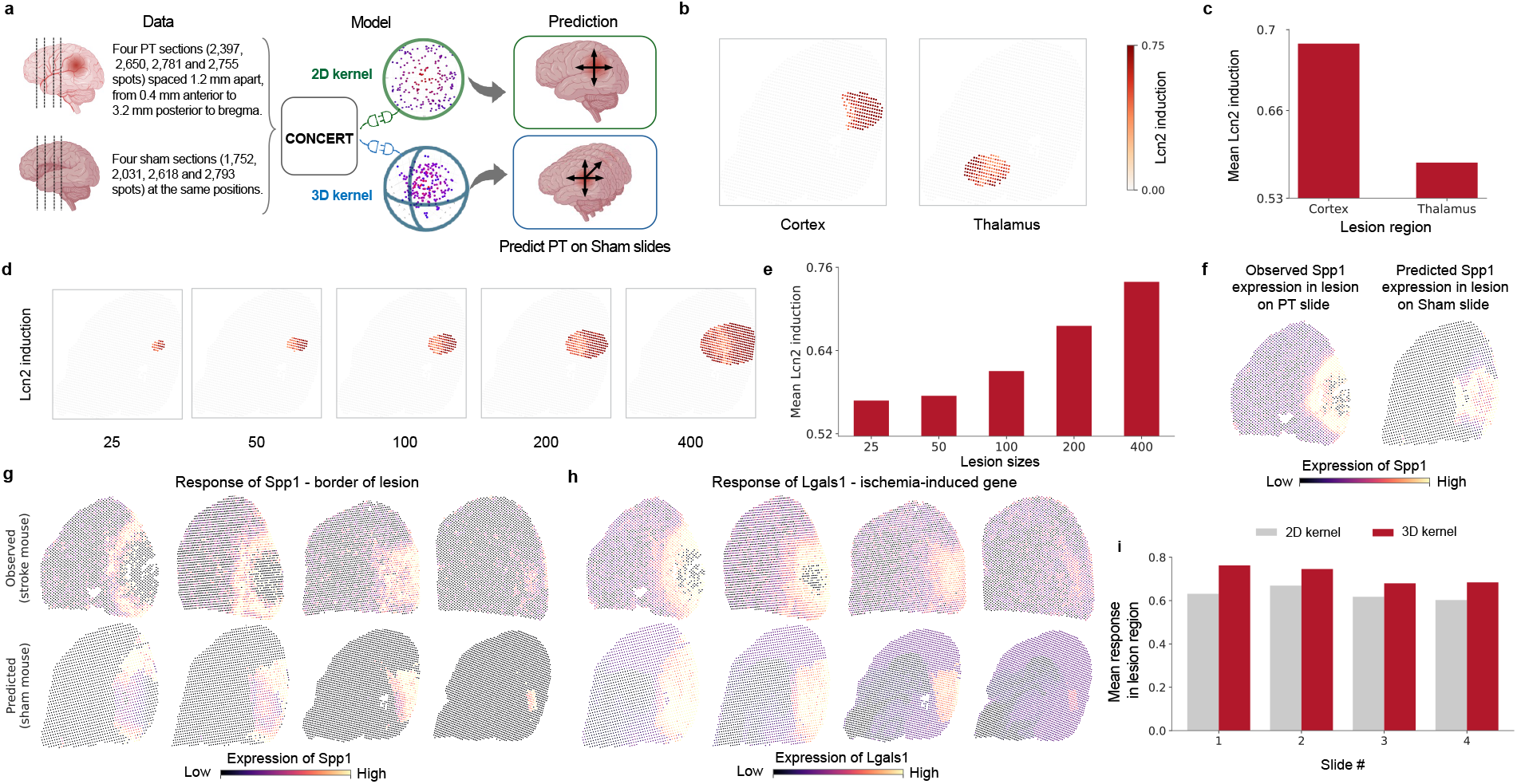
CONCERT predicts region-specific, dose-integrated and three-dimensional responses to ischemic injury. **a**, The photothrombotic stroke dataset: four coronal sections from a stroke (PT) brain and four from a sham brain. CONCERT is fitted on the four stroke and four sham sections, and every counterfactual below is imposed on intact sham tissue. **b**, Predicted response to an identical 200-spot lesion placed in cortex and in thalamus, shown on the slide, and **c**, quantified inside the lesion. **d**, Predicted response to a contiguous cortical lesion grown from 25 to 400 spots, and **e**, quantified inside the lesion. **f**, Observed (left) and predicted (right) expression of the peri-infarct marker *Spp1*. **g, h**, The same comparison across the four sections in three dimensions, for the perilesional marker *Spp1* (**g**) and the ischemia-induced gene *Lgals1* (**h**); upper row, observed in the stroke brain; lower row, predicted on the sham brain. A separate color scale was independently defined for each section. **i**, Level of the predicted response across sections under a two-dimensional and a three-dimensional footprint field.

**Extended Data Fig. 5:**
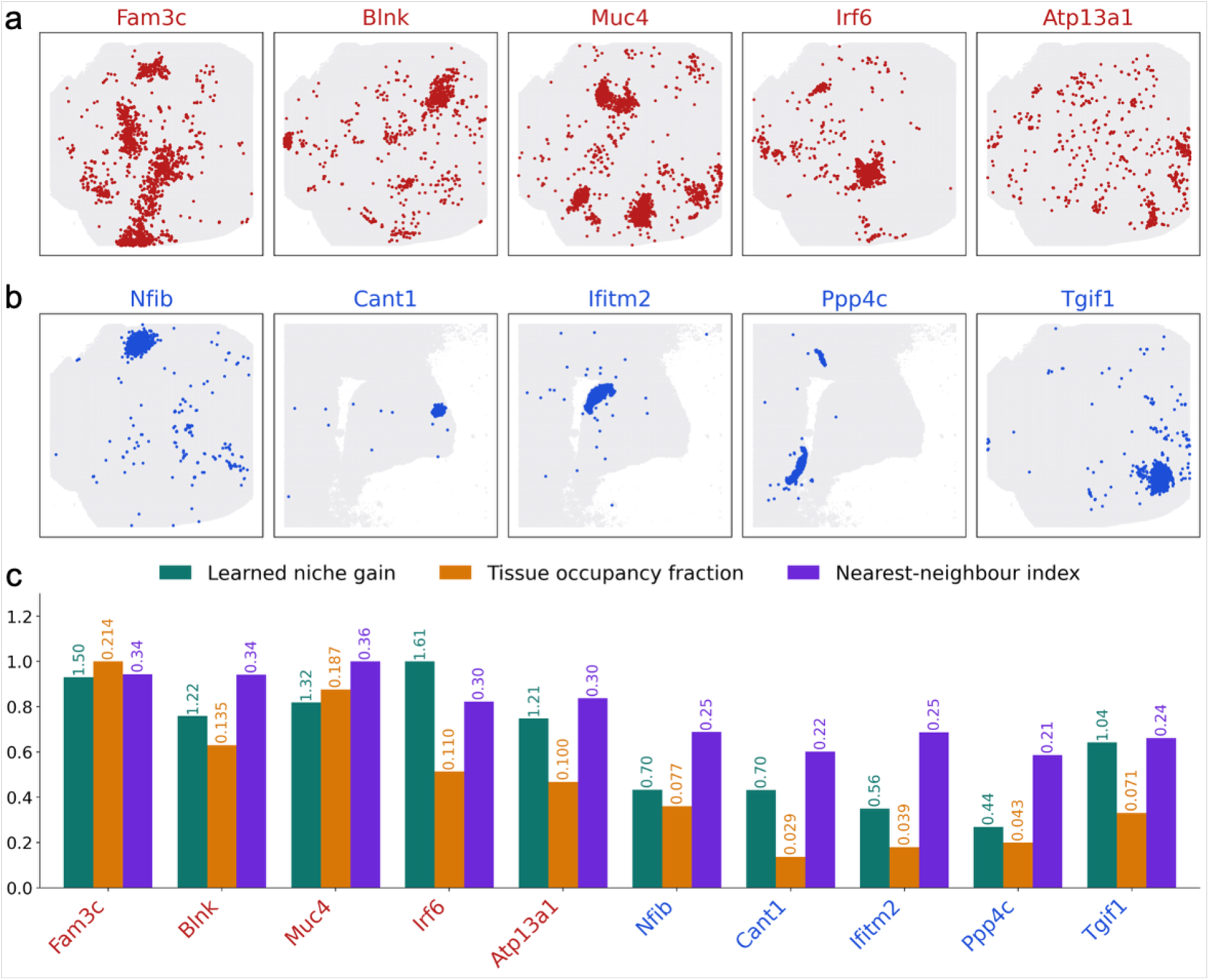
The learned niche gain tracks how widely a perturbation is spread across the tissue. **a, b**, Observed layout of the perturbed cells for ten perturbations spanning the range of the learned niche gain, the per-perturbation scalar that sets how strongly CONCERT allows a perturbation’s predicted response to be modulated by its local microenvironment. The ten were drawn from a matched band of perturbed-cell numbers so that all panels carry comparable numbers of marks. Gray points are the other cells on that section, drawn as a one-in-four subsample for legibility; colored points are all cells carrying the indicated perturbation, in measured array coordinates min–max scaled per section. **a**, the five perturbations with the highest niche gain (red). **b**, the five with the lowest (blue). Each perturbation is shown on the section carrying the majority of its cells. **c**, Three quantities for the same ten perturbations, one color per quantity. They are the learned niche gain, the fraction of tissue area the perturbation reaches, and the nearest-neighbor index. The nearest-neighbor index is the mean distance from a perturbed cell to the nearest cell carrying the same perturbation, divided by the distance expected under a random scatter of the same number of cells. Bar height is each quantity divided by its largest value among the ten, and the value is printed on every bar. Perturbation names are colored by group, as in a and b.

