## Supplementary material for "Generative modeling of spatially resolved perturbation responses with CONCERT": all supp contents

#### Supplementary Notes

- **Supplementary Note 1**. Spatial fields and niche modulation.
- **Supplementary Note 2**. Loss functions.
- **Supplementary Note 3**. Continuous-time conditioning in CONCERT.
- **Supplementary Note 4**. Adapting CONCERT to a three-dimensional tissue stack.
- **Supplementary Note 5**. Reading the learned kernel parameters.
- **Supplementary Note 6**. Deconvolution and patch imputation.
- **Supplementary Note 7**. Tasks designed for spatial perturbation data.
- **Supplementary Note 8**. Neighbor averaging on spatial perturbation tasks.
- **Supplementary Note 9**. Sensitivity analyses of CONCERT.
- **Supplementary Note 10**. Attributes encoded and implementation details per dataset.
- **Supplementary Note 11**. Marker availability and counterfactual protocol in the stroke case study.

#### Supplementary Tables (provided in a separate Excel file)

- **Supplementary Table 1**. Significance of the comparisons between CONCERT and every competing method on the log-fold-change rank correlation.
- **Supplementary Table 2**. Significance of the comparisons between CONCERT and every competing method on the log-fold-change RMSE.

#### Supplementary Figures

- **Supplementary Figure 1**. Problems in spatial perturbation technologies.
- **Supplementary Figure 2**. Mechanisms of the dispersion of perturbation effects in tissue.
- **Supplementary Figure 3**. Sensitivity of CONCERT to the hyper-parameters of its spatial components on Perturb-map (log-fold-change rank correlation).
- **Supplementary Figure 4**. Sensitivity of CONCERT to the hyper-parameters of its spatial components on Perturb-map (log-fold-change RMSE).
- **Supplementary Figure 5**. Sensitivity of CONCERT to the hyper-parameters of its spatial components on SPAC-seq (log-fold-change rank correlation).
- **Supplementary Figure 6**. Sensitivity of CONCERT to the hyper-parameters of its spatial components on SPAC-seq (log-fold-change RMSE).
- **Supplementary Figure 7**. Sensitivity of CONCERT to the hyper-parameters of its spatial components on Perturb-map and SPAC-seq, shown on a common standardized scale (log-fold-change rank correlation).
- **Supplementary Figure 8**. Sensitivity of CONCERT to the hyper-parameters of its spatial components on Perturb-map and SPAC-seq, shown on a common standardized scale (log-fold-change RMSE).
- **Supplementary Figure 9**. Deconvolution of pseudo-bulk spots on SPAC-seq.
- **Supplementary Figure 10**. Deconvolution of periphery spots on Perturb-map.
- **Supplementary Figure 11**. Imputation of held-out knockout patches on SPAC-seq.
- **Supplementary Figure 12**. Observed and predicted DSS-marker score on every slide of Mouse 1 (`062921_D0_m3a`), sampled at day 0.
- **Supplementary Figure 13**. Observed and predicted inflamed cells on every slide of Mouse 1 (`062921_D0_m3a`), sampled at day 0.
- **Supplementary Figure 14**. Observed and predicted DSS-marker score on every slide of Mouse 2 (`082421_D0_m6`), sampled at day 0.
- **Supplementary Figure 15**. Observed and predicted inflamed cells on every slide of Mouse 2 (`082421_D0_m6`), sampled at day 0.
- **Supplementary Figure 16**. Observed and predicted DSS-marker score on every slide of Mouse 4 (`092421_D3_m1`), sampled at day 3.

- **Supplementary Figure 17**. Observed and predicted inflamed cells on every slide of Mouse 4 (`092421_D3_m1`), sampled at day 3.
- **Supplementary Figure 18**. Observed and predicted DSS-marker score on every slide of Mouse 7 (`092421_D3_m4`), sampled at day 3.
- **Supplementary Figure 19**. Observed and predicted inflamed cells on every slide of Mouse 7 (`092421_D3_m4`), sampled at day 3.
- **Supplementary Figure 20**. Observed and predicted DSS-marker score on every slide of Mouse 8 (`062221_D9_m3`), sampled at day 9.
- **Supplementary Figure 21**. Observed and predicted inflamed cells on every slide of Mouse 8 (`062221_D9_m3`), sampled at day 9.
- **Supplementary Figure 22**. Observed and predicted DSS-marker score on every slide of Mouse 13 (`100221_D9_m5`), sampled at day 9.
- **Supplementary Figure 23**. Observed and predicted inflamed cells on every slide of Mouse 13 (`100221_D9_m5`), sampled at day 9.
- **Supplementary Figure 24**. Observed and predicted DSS-marker score on every slide of Mouse 14 (`082421_D21_m1`), sampled at day 21.
- **Supplementary Figure 25**. Observed and predicted inflamed cells on every slide of Mouse 14 (`082421_D21_m1`), sampled at day 21.
- **Supplementary Figure 26**. Observed and predicted DSS-marker score on every slide of Mouse 15 (`082421_D21_m2`), sampled at day 21.
- **Supplementary Figure 27**. Observed and predicted inflamed cells on every slide of Mouse 15 (`082421_D21_m2`), sampled at day 21.
- **Supplementary Figure 28**. Learned spatial parameters against the observed layout of each perturbation on Perturb-FISH and SPAC-seq.

### Supplementary Notes

#### Note 1: Spatial fields and niche modulation

This note gives the exact form of the components summarized in the Methods, together with the hyper-parameter values used on SPAC-seq. The settings that differ on the other datasets are listed in Note 10.

**Basal spatial field.** The basal spatial field maps a cell's coordinates to the mean and variance of the spatial basal latent. A projection matrix $\mathbf{W}_{\text{rff}} \in \mathbb{R}^{2\times F}$ is drawn once from $\mathcal{N}(0, \omega_{\text{rff}}^2)$ and an offset $\mathbf{b}_{\text{rff}} \in \mathbb{R}^F$ once from $\mathcal{U}(0, 2\pi)$; both are registered as fixed buffers. The coordinates of each slide are min–max scaled to $[0, 20]$ and, when several slides are modelled jointly, each slide is offset into its own non-overlapping region so that the neighbors of every cell lie on its own slide. For a cell at these coordinates $\mathbf{s}$, the random Fourier features are $\boldsymbol{\psi}(\mathbf{s}) = [\cos(\mathbf{s}^\top \mathbf{W}_{\text{rff}}/\rho + \mathbf{b}_{\text{rff}}), \sin(\mathbf{s}^\top \mathbf{W}_{\text{rff}}/\rho + \mathbf{b}_{\text{rff}})] \in \mathbb{R}^{2F}$. These pass through a single hidden layer with ReLU activation and then through a mean-field variational linear head, which returns the mean of the spatial basal latent. Its variance is a separate learnable parameter per latent dimension, with log-variance initialized at $-6$. The head places an independent Gaussian $\mathcal{N}(0, \sigma_0^2)$ prior on each weight and contributes the closed-form KL divergence $\text{KL}_{\text{basal}} = \text{KL}(q(\mathbf{W}_h) \,\|\, \mathcal{N}(0, \sigma_0^2))$ to the objective, amortized by the ratio of batch size to training-set size so that it is commensurate with the batch-summed latent KL terms. We use $F = 128$ features, bandwidth $\omega_{\text{rff}} = 0.15$, hidden width 128, $\rho = 20$, prior $\sigma_0 = 1$, and a spatial basal latent of dimension 2. On a dataset that encodes more than one tissue class, the tissue embedding is concatenated to $\boldsymbol{\psi}(\mathbf{s})$ before the hidden layer. The field is then conditioned on region identity, so two cells at the same coordinate but in different regions receive different spatial latents. This applies to the Perturb-map and stroke data. SPAC-seq carries a single class, and the field reduces to a function of the coordinates alone. Because $\mathbf{W}_{\text{rff}}$ and $\mathbf{b}_{\text{rff}}$ are fixed, the bandwidth of the field stays at its set value throughout training.

**Perturbation spatial field.** The perturbation spatial field produces a gene-space log-fold-change shift for each cell. It has a spatially constant core $f_{\text{core}}(\mathbf{e}_p) \in \mathbb{R}^G$, a bias-free linear map from the perturbation embedding to gene space whose weights are initialized from $\mathcal{N}(0, 0.01^2)$, plus a coordinate-conditioned residual. The residual is produced by an MLP with hidden width 256 applied to the concatenation of the perturbation embedding $\mathbf{e}_p$ and radial basis features of position. The radial basis features are $\kappa_m(\mathbf{s}; \ell_p) = \exp\!\big(-\|\mathbf{s}/\rho - \mathbf{r}_m\|^2/(2\ell_p^2)\big)$ for $m = 1, \ldots, M_r$, where the reference locations $\mathbf{r}_m$ are initialized on a regular $\sqrt{M_r} \times \sqrt{M_r}$ grid over the unit square and are learnable. We use $M_r = 16$ on SPAC-seq and $M_r = 64$ on Perturb-map, with $\rho = 20$. The final layer of the MLP is initialized with weights of standard deviation $10^{-3}$ and zero bias, so the residual starts close to zero and training begins from a nearly uniform effect $f_{\text{core}}(\mathbf{e}_p)$.

**Per-perturbation lengthscale.** The lengthscale is predicted from the perturbation embedding by a single linear head shared across perturbations, and squashed into a fixed band, $\ell_p = \ell_{\min} + (\ell_{\max} - \ell_{\min})\,\text{sigmoid}(\mathbf{w}_\ell^\top \mathbf{e}_p + b_\ell)$. The head weights are initialized at zero and the bias is set so that $\ell_p$ starts at a chosen value $\ell_0$, that is $b_\ell = \log\!\Big(\frac{\ell_0 - \ell_{\min}}{\ell_{\max} - \ell_0}\Big)$. On SPAC-seq we use $\ell_{\min} = 0.15$, $\ell_{\max} = 0.90$ and $\ell_0 = 0.25$, in the same normalized units as the rescaled coordinates. Multiplying by the coordinate scale $\rho$ gives the spatial units used in Notes 9 and 10, so this band is 3–18 spatial units. On Perturb-map the band is 60–360 spatial units, with $\ell_0 = 100$.

**Niche modulation and gate.** For each cell we take the $k_{\text{nn}}$ nearest cells in space within the training batch. The non-spatial basal latent of the cell and of each neighbor is passed through one shared linear projection. On SPAC-seq, where control guides are present, the non-spatial basal latent of every cell carrying a guide (control or knockout) is set to zero. The basal state of such a cell is then given by the spatial basal field alone, and its contribution to a descriptor is constant. The neighbor projections are averaged with weights proportional to $1/(1 + d_{ij}^2)$ and normalized over the neighborhood, and this average is concatenated with the cell's own projection and passed through a two-layer MLP to give the niche descriptor $\mathbf{z}^{\text{niche}} \in \mathbb{R}^{d_{\text{niche}}}$. The latent entering the descriptor is detached, so the descriptor conditions the modulation without changing the latent encoder. The descriptor modulates the perturbation shift in two ways. A scalar per-cell factor rescales the whole shift,

$$\gamma = 1 + g_p \cdot \tanh\big(f_\gamma(\mathbf{e}_p, \mathbf{z}^{\text{niche}})\big), \qquad g_p = \min\big(\text{softplus}(\mathbf{w}_g^\top \mathbf{e}_p + b_g),\ g_{\max}\big), \tag{1}$$

where $f_\gamma$ is an MLP of the perturbation embedding and the descriptor jointly. Because $\tanh$ is signed and $g_p$ may exceed one, $\gamma$ can cross zero, so the same perturbation can act in opposite directions in different neighborhoods, as well as with different strength. The gain $g_p$ is per perturbation and bounded by $g_{\max}$, and its head is initialized with a bias of $-4$ so that $g_p \approx 0$ and the layer starts at the identity. Second, a per-gene additive residual is formed as a low-rank bilinear interaction between the perturbation embedding and the descriptor, $\boldsymbol{\beta} = \mathbf{W}_\beta\big(\mathbf{W}_{\beta p}\mathbf{e}_p \odot \mathbf{W}_{\beta n}\mathbf{z}^{\text{niche}}\big)$ with rank $r_\beta$ and $\mathbf{W}_\beta$ initialized at zero. The residual is added with unit weight, independently of $g_p$. The modulated shift is $\gamma\,\Delta_p + \boldsymbol{\beta}$. We use $k_{\text{nn}} = 25$, $d_{\text{niche}} = 8$, $r_\beta = 8$ and $g_{\max} = 3$. The penalty $\mathcal{L}_g$ that holds this layer near the identity is given in Note 2.

A gate then controls how much of the shift reaches a given cell. An MLP $f_{\text{gate}}$ of hidden width 64 with ELU activation takes the two halves of the basal latent, the spatial and the non-spatial component, with gradients stopped at its input. It outputs a scalar, which a sigmoid maps to $a$, giving the gated shift $\Delta_p' = a\,(\gamma\,\Delta_p + \boldsymbol{\beta})$ defined in the Methods. Its final layer is initialized with zero weight and bias 3, so the gate starts at $\approx 0.95$. The result is clamped to $[-5, 5]$ and applied to the cells carrying a real knockout.

**Expression encoder and attribute correction.** The basal embedding is produced by an encoder applied to the cell's expression profile, with hidden widths 512 and 256, batch normalization, ELU activation and dropout 0.1. On SPAC-seq, a learned per-gene shift in log-mean expression, clamped to $[-5, 5]$, is added for every cell carrying a guide, control and knockout alike. It absorbs the offset between guide-carrying and unassigned cells, so that the knockout shift describes the effect of the knockout beyond this offset. Note 10 lists the datasets on which it is used.

#### Note 2: Loss functions

This note gives the explicit form of the three parts of the objective of the Methods. Throughout, $b$ is the mini-batch size and $N$ the number of cells in the dataset.

**Data term.** $\mathcal{L}_{\text{data}} = \mathcal{L}_{\text{recon}} + \mathcal{L}_{\text{mse}}$. The first contribution is the negative binomial loss on raw counts, evaluated with the decoded mean, the per-perturbation dispersion and the per-cell size factor, summed over the genes and the cells of the mini-batch. The second is the mean squared error between the decoded mean $\boldsymbol{\mu}^k$ and the raw counts of the same cell, averaged over cells and genes. The decoder also carries a per-slide gene-wise bias.

**Latent term.** $\mathcal{L}_{\text{latent}} = \lambda_{\text{KL}}(\text{KL}_{\text{G}} + \text{KL}_{\text{basal}}) + \mathcal{L}_2$. The non-spatial basal latent is a diagonal Gaussian posterior against a standard normal prior,

$$\text{KL}_{\text{G}} = \frac{1}{2} \sum_l \left[ \tilde{\varsigma}_l^2 + \tilde{m}_l^2 - \log \tilde{\varsigma}_l^2 - 1 \right],$$

where $\tilde{m}_l$ and $\tilde{\varsigma}_l^2$ are the posterior mean and variance of latent dimension $l$. A free-bits floor is applied per dimension before summing, so that each dimension may carry up to $\eta$ nats free of penalty; we use $\eta = 0.5$. This discourages dimensions from collapsing onto the prior early in training.

The basal spatial field is deterministic in its features, and its uncertainty and KL term come from the mean-field variational head, whose weights $\mathbf{W}_h$ have factorized Gaussian posteriors $q(w_{ij}) = \mathcal{N}(\mu_{ij}, \varsigma_{ij}^2)$ against a shared prior $\mathcal{N}(0, \sigma_0^2)$. The KL is available in closed form,

$$\text{KL}_{\text{basal}} = \sum_{ij} \left[ \log \frac{\sigma_0}{\varsigma_{ij}} + \frac{\varsigma_{ij}^2 + \mu_{ij}^2}{2\sigma_0^2} - \frac{1}{2} \right],$$

and is amortized by the factor $b/N$ so that it is commensurate with the batch-summed latent KL. We use $\sigma_0 = 1$. The weight $\lambda_{\text{KL}}$ shared by the two divergences is warmed linearly from 0 to 1 over the first 20 epochs, so that reconstruction is established before the latents are compressed. The remaining contribution $\mathcal{L}_2$ is the squared norm of the basal embedding, averaged over cells, which keeps the basal embedding from absorbing information belonging to the attribute embeddings.

**Perturbation term.** $\mathcal{L}_{\text{pert}} = \mathcal{L}_{\text{LFC}} + \mathcal{L}_M + \mathcal{L}_\ell + \lambda_g \mathcal{L}_g$. For each pair of a niche $n$ and a perturbation $p$ observed in that niche, we average the model's predicted shift over the cells carrying $p$ in $n$ and compare it with the measured knockout-versus-control log-fold change $\mathbf{t}_{n,p}$ of the same pair,

$$\mathcal{L}_{\text{LFC}} = \frac{1}{|\mathcal{Q}|} \sum_{(n,p) \in \mathcal{Q}} \left\| \bar{\Delta}_{n,p} - \mathbf{t}_{n,p} \right\|_{/G}^2,$$

where $\mathcal{Q}$ is the set of pairs in the current mini-batch for which a measured target exists and $\| \cdot \|_{/G}^2$ is the mean over genes. This supervises the perturbation spatial field directly on the quantity used for evaluation, in addition to the reconstruction.

$\mathcal{L}_M$ is a Gaussian-weighted Moran's $I$ of the non-spatial basal latent, computed with pair weights $w_{ij} = \exp(-d_{ij}^2/2r^2)$ so that the penalty acts on autocorrelation at approximately the footprint scale $r$. We sum the squared statistic over latent dimensions and use $r = 4$ in the per-slide coordinate units (range 0–20).

$\mathcal{L}_\ell = \mathbb{E}[\max(0, \ell_p - \ell_{\max})^2 + \max(0, \ell_{\min} - \ell_p)^2]$ is a squared-hinge penalty on the per-perturbation lengthscale. The sigmoid parameterization keeps $\ell_p$ within the specified band in the reported models.

The remaining term, $\mathcal{L}_g = \mathbb{E}_p[g_p] + \mathbb{E}[\mathbf{W}_\beta^2]$, penalizes the mean niche gain and the squared modulation weights. How fitted gain relates to spatial layout is analyzed in Note 5.

**Weights.** Every contribution above enters the objective with weight one, apart from $\mathcal{L}_g$, for which $\lambda_g = 10^{-3}$. The one schedule is the warm-up of $\lambda_{\text{KL}}$ described above. All parameters are optimized with Adam at a learning rate of $10^{-4}$ and weight decay $10^{-6}$.

#### Note 3: Continuous-time conditioning in CONCERT

For a perturbation time course, we query the model at unobserved days within the sampled interval. We therefore encode time as a continuous variable, which defines an embedding for every day of the course.

Let $t \in [0, 1]$ be the normalized day, obtained by dividing the collection day by the last sampled day of the course. We map $t$ to a time embedding $\mathbf{e}_t \in \mathbb{R}^{d_t}$ by first expanding it into Fourier features on a geometric band of frequencies and then passing them through a small MLP:

$$\boldsymbol{\psi}_t(t) = \big[\, t,\ \sin(\omega_1 t), \cos(\omega_1 t),\ \ldots,\ \sin(\omega_{N_B} t), \cos(\omega_{N_B} t) \,\big], \qquad \omega_b = 2^{b-1}\pi, \tag{2}$$

$$\mathbf{e}_t = \mathbf{W}_2 \, \mathrm{SiLU}(\mathbf{W}_1 \boldsymbol{\psi}_t(t)) \,, \tag{3}$$

where $N_B$ is the number of Fourier bands and $\mathbf{W}_1, \mathbf{W}_2$ are learnable. We use $N_B = 6$ and $d_t = 16$. The Fourier expansion is smooth in $t$, allowing the fitted time course to vary nonlinearly between sampled days.

The time embedding enters the perturbation spatial field in two places. First, $\mathbf{e}_t$ is concatenated onto the perturbation embedding before the lengthscale head, so the lengthscale becomes a function of time, $\ell(t)$, squashed by a sigmoid into a fixed interval $(\ell_{\min}, \ell_{\max})$. On this dataset the interval is wide relative to the rescaled slide, so the change of the spatial pattern over time is carried mainly by the second path. Second, $\mathbf{e}_t$ is supplied to the MLP that produces the spatial field, so that time can change the spatial pattern of the predicted shift itself.

Imputing an unsampled day is therefore a forward pass of the fitted model at that value of $t$, evaluated jointly with the cell's location, its perturbation state and the per-slide gene-wise bias of the decoder. The imputed profile follows the fitted time function across the whole course.

#### Note 4: Adapting CONCERT to a three-dimensional tissue stack

The eight stroke sections are separate slices whose raw coordinate frames are unrelated. A common frame within each brain is therefore built first by pairwise partial alignment with PASTE2, as described below. The remainder of this note describes what CONCERT does once that frame exists.

**Only the perturbation spatial field becomes three-dimensional.** Depth is added to the perturbation footprint field alone. The basal spatial field and the niche descriptor remain functions of $(x, y)$ on the registered frame shared by the four sections of a brain. The basal field is therefore shared across the sections of a brain, apart from a per-section gene-wise bias in the decoder, and the nearest neighbors used by the niche descriptor can come from any section. The section index enters the footprint field alone. The footprint field is evaluated at $(x, y, z)$, and every other spatial component at $(x, y)$.

**Reference locations are replicated per section.** The footprint field is a radial basis expansion about learnable reference locations. With four ordinal levels, the two-dimensional grid is replicated at each section level, and the replicas move freely during training. A field with $M$ in-plane locations therefore carries $4M$ locations in the three-dimensional arm. The radial basis and the per-perturbation lengthscale are otherwise exactly those of the two-dimensional model, so setting the number of levels to one recovers the two-dimensional model identically.

**Building the three-dimensional coordinate frame.** Consecutive sections within each mouse are registered pairwise with PASTE2 [1] (version 1.0.1), which solves a partial optimal-transport problem. Consecutive sections of the same brain overlap partially, and partial transport matches the overlapping tissue. Six pairs are registered, PT1–PT2–PT3–PT4 and Sham1–Sham2–Sham3–Sham4. The transport cost between two sections is the GLM-PCA expression distance with 50 latent dimensions on the 2,000 shared genes with the highest total counts (of 2,003). The overlap fraction is chosen per pair by PASTE2's own model selection over a ladder from 0.99 down to 0.05. The selected overlap differs substantially between pairs, with 0.8 for PT1–PT2 and PT3–PT4, 0.5 for PT2–PT3, 0.6 for Sham1–Sham2 and Sham3–Sham4, and 0.3 for Sham2–Sham3. Two settings extend the stock implementation and are recorded with the released coordinates. First, the GLM-PCA fit is seeded, which makes the result reproducible. Second, its penalty is raised along the ladder 1, 10, 100, 1,000 until the fit converges; a penalty above the default value of 1 was required on five of the six pairs. CONCERT rescales each registered brain to the common coordinate range and uses the registered coordinates directly. The stroke and sham brains are registered separately.

**Ordinal section coordinate.** For the 3D stroke analysis, the third coordinate is the ordinal section index, $z \in \{0, 1, 2, 3\}$. The four sections were collected at 1.2-mm intervals, at 0.4 mm anterior and 0.8, 2.0, and 3.2 mm posterior to bregma. We therefore represent section position by order. Within each brain, the index is rescaled to $[0, 1]$ and multiplied by $\rho_z = 0.25$, placing adjacent sections $0.25/3 \approx 0.083$ normalized units apart and the full four-section stack across 0.25 units. This scaling allows the perturbation-footprint field to use one isotropic lengthscale across the registered in-plane coordinates and the ordinal section axis. Lesion depth (rim-to-core distance) is calculated only post hoc for evaluation and is not provided to the model.

#### Note 5: Reading the learned kernel parameters

For each perturbation $p$, the trained model yields two summary parameters: the footprint lengthscale $\ell_p$, which sets the spatial scale over which its effect can vary, and the niche gain $g_p$, which scales the niche-conditioned modulation. Both are estimated jointly with the rest of the model through the objective in Note 2. We then relate these fitted parameters to descriptors of the spatial layout of perturbed cells.

We quantify spatial layout using descriptors computed from the cell coordinates after training. The occupancy fraction is the area of the $20 \times 20$ coordinate bins containing at least one perturbed cell divided by the area of the slide's bounding box, averaged over slides with weights proportional to cell number. The nearest-neighbor (Clark–Evans) index is the observed mean nearest-neighbor distance among perturbed cells divided by the value expected under a random layout of the same density; values below one indicate clustering and values above one dispersion. We also characterize focus structure by DBSCAN (radius 2 coordinate units; minimum of 3 cells per focus), recording the number of foci, their mean size, the fraction of cells in the largest focus, and the fraction labelled as noise.

Because the number of perturbed cells is related to tissue coverage, we estimate partial Spearman correlations conditional on the number of perturbed cells and the $L_2$ norm of the learned perturbation embedding. We correct $P$-values with the Benjamini–Hochberg procedure across the 15 layout and exposure covariates evaluated for each parameter. The analysis includes the 87 knockouts scored in the within-niche benchmark; the threshold defining perturbation carriage is fixed at 0.5 throughout.

After adjustment, niche gain increased with the fraction of the section reached (partial $\rho = 0.36$, corrected $P = 0.009$) and with the nearest-neighbor index (partial $\rho = 0.32$, corrected $P = 0.020$). Footprint lengthscale decreased with the same two descriptors (partial $\rho = -0.30$, corrected $P = 0.037$; partial $\rho = -0.28$, corrected $P = 0.049$). None of the focus-structure descriptors was associated with either parameter after correction. These results describe associations between fitted parameters and tissue layout; they do not establish a causal effect of layout. Extended Data Fig. 5 illustrates the result for ten perturbations selected from a matched range of perturbed-cell numbers.

**Independent analysis on Perturb-FISH.** We applied the same experimental analysis to the Perturb-FISH tumor section, an image-based assay. Of the 35 knockout targets in the section, 31 were carried by at least 25 cells and were included. We calculated the same spatial descriptors and partial Spearman correlations as for SPAC-seq, controlling for the number of perturbed cells and the $L_2$ norm of the learned perturbation embedding, with multiple-testing correction across the 12 covariates available for this dataset. The effect estimates followed the same directions as in SPAC-seq: niche gain increased with occupancy fraction (partial $\rho = 0.42$, corrected $P = 0.11$) and nearest-neighbor index (partial $\rho = 0.31$, corrected $P = 0.26$), whereas footprint lengthscale decreased with these descriptors (partial $\rho = -0.21$ and $-0.22$, respectively; corrected $P = 0.63$ for both). The Perturb-FISH model configuration is given in Note 10.

#### Note 6: Deconvolution and patch imputation

Both analyses apply a trained CONCERT model at inference, with the coordinate framing, normalization, gene order and perturbation codes used in training. Deconvolution is evaluated on SPAC-seq slide 2-1 (Supplementary Fig. 9) and on Perturb-map slide 0 (Supplementary Fig. 10). Patch imputation is evaluated on SPAC-seq slide 2-2 (Supplementary Fig. 11). Each SPAC-seq analysis uses a model trained on its own slide.

**Pseudo-bulk spots on SPAC-seq.** The cells of slide 2-1 are grouped into 34,921 non-overlapping pseudo-bulk spots of about nine spatially nearest cells, and the expression of a spot is the mean of the counts of its cells. The composition of a spot is the fraction of its cells in each perturbation state, which gives the ground truth. Ten knockouts are analyzed. A spot is a mixture spot when exactly one of the ten knockouts is present, at a fraction between 0.15 and 0.85, and at least 10% of its cells are unassigned; 3,775 mixture spots are scored. A spot is a pure spot when at least 80% of its cells carry a single state.

**Signatures.** For each mixture spot, CONCERT supplies location-matched signatures, the expected expression of a knockout cell and of an unassigned cell at the location of the spot. To generate the signature of a state $p$, both the spatial group and the perturbation shift are set to $p$, the basal embedding is obtained by encoding the spot profile with the expression encoder, and the decoded mean is averaged over 25 posterior samples. Each knockout is assigned a region of interest of radius 1,500 raw coordinate units around its patch. The empirical signature of a state is the mean profile of the pure spots of that state on the slide. All profiles are compared in log counts per ten thousand on 1,500 highly variable genes.

**Unmixing.** The quantity estimated for each mixture spot is its knockout fraction $w$, with the remainder assigned to the unassigned state, and it is estimated in four ways. Non-negative least squares (NNLS) unmixes the spot profile against the two signatures, once with empirical and once with CONCERT signatures, and the weights are rescaled to sum to one. A learned unmixing head, a multilayer perceptron that maps the spot profile and the two signatures to $w$, is applied once with empirical and once with CONCERT signatures. The head is trained on synthetic mixtures of pure spots of the same slide, with $w$ drawn uniformly between 0.15 and 0.85 and both signature types as inputs. Agreement with the true knockout fractions is summarized by the mean absolute error and the Pearson correlation over the 3,775 mixture spots, and Supplementary Fig. 9c shows the per-spot difference in absolute error between NNLS with empirical signatures and the learned head with CONCERT signatures.

**Deconvolution on Perturb-map.** On Perturb-map slide 0, the analyzed spots are the 48 non-knockout, non-normal spots of the patch grown outward from the Jak2-knockout spots until it reaches normal tissue. For spots adjacent to Jak2-knockout spots, the learned head gives the Jak2-knockout weight and the remainder is split among the neighboring states. For the other spots, NNLS on the CONCERT profiles of unperturbed tumor and normal tissue gives the composition. Each spot is expanded into seven sub-spots, one at its center and six on a ring at 0.36 times the spot spacing, and the states are allotted to the sub-spots by largest-remainder rounding of the composition. For each state, the local reference profile is the mean log counts per ten thousand of the pure spots of that state within a radius of 15 coordinate units, and of all pure spots of that state when that radius is empty. Supplementary Fig. 10c reports the Spearman correlation over 1,500 highly variable genes between this reference and either the observed spot or the CONCERT profile of that state at the spot location.

**Patch imputation.** The five most frequent knockouts on slide 2-2 are evaluated. For each, the largest spatial cluster of its cells is found with DBSCAN (radius three times the median nearest-neighbor distance), and the cells on one side of the first principal axis of that cluster are held out. An 800-cell patch of unassigned cells is also held out. The model is trained on the remaining cells of the slide, with the five evaluated knockouts as the perturbed states. At inference, each held-out cell receives a basal embedding obtained by encoding the mean expression of its five nearest retained cells of the same knockout, and the perturbation shift of its knockout is applied at its held-out coordinates. The predicted profiles are compared with the measured profiles of the held-out cells.

**Patch imputation baselines and scoring.** Tangram, run in cell mode with an RNA-count-based density prior, SpatialScope and SpaIM are applied to the same masked slide following their tutorials. Each method is scored on its own gene panel, restricted to the 200 genes of highest variance in the observed profiles. Expression is converted to log counts per ten thousand before scoring. The per-cell score is the Spearman correlation between the imputed and measured profile of a cell, averaged over the held-out cells of a knockout and then over the five knockouts. The measured log-fold change of a knockout is the mean expression of its held-out cells minus the mean expression of the 50 nearest retained unassigned cells. For CONCERT, the predicted log-fold change is its imputed knockout expression minus its imputed unperturbed expression of the same cells. For SpaIM, Tangram and SpatialScope, it is the imputed expression of the held-out knockout cells minus the measured reference. The log-fold-change score is the Spearman correlation between the predicted and measured log-fold changes.

#### Note 7: Tasks designed for spatial perturbation data

For task definition, we refer to each data entry as a cell, which may correspond to a cell or a spot in the real data. With spatial information, the counterfactual prediction of response gene expression (rGEX) becomes a question about a location as well as about a cell: given where a cell sits, what would it look like under a perturbation that was measured somewhere else? We use this to define two benchmark tasks (Fig. 1a), which differ in how far the perturbation effect has to travel to reach the cell being predicted.

**Within-niche prediction.** A spatial patch of unperturbed cells is withheld from training and predicted into a target perturbation state that is carried by measured cells of the same local niche. Prediction is made jointly for the cells of the patch, with at most 500 cells predicted per niche. Cells of one type cluster in tissue, so the cells inside a patch are interdependent. The measured cells of the target state in the same niche provide the reference, so the comparison asks what that perturbation does in that microenvironment.

**Cross-slide prediction.** A perturbation is withheld from an entire target slide while remaining observed on the donor slides, and its response is predicted for a withheld patch of unperturbed cells on the target slide. The model therefore transfers the effect from the donor slides, which differ in composition and in sampling density, to the slide it predicts. This task uses perturbations observed across multiple slides and tests the spatial transfer of a perturbation that is observed elsewhere in the training data.

Both tasks share the same inference contract. The model receives a cell's location, its measured unperturbed expression and a target perturbation, and returns that cell's perturbed expression. Predictions are always made on unperturbed cells and scored against measured perturbed cells.

#### Note 8: Neighbor averaging on spatial perturbation tasks

The benchmark includes neighbor-averaging strategies, which predict a cell from the measured cells around it. This note describes how these strategies and CONCERT draw on the available cells in three settings.

**The target state is far from the predicted cells.** When the cells carrying the target state lie far from the cells being predicted, a neighbor average draws on the same measured cells across the region. CONCERT evaluates its perturbation spatial field at each cell's own coordinates.

**An attribute is confounded with the perturbation.** When a perturbation is observed in one tissue context, as Jak2-KO is in tumor tissue in Perturb-map, the measured cells carrying it also carry that context. CONCERT encodes the perturbation and the tissue state in separate codebooks, which are set independently at inference.

**The perturbation was measured on another slide.** Neighbor averaging draws on the measured cells of the slide being predicted. CONCERT represents a perturbation effect as a function of position and perturbation identity, which is evaluated at coordinates on the target slide.

#### Note 9: Sensitivity analyses of CONCERT

**Design.** We varied ten hyper-parameters of CONCERT's spatial components one at a time from the reported configuration, holding all others at their reported values. Four belong to the perturbation footprint field. They are the band its lengthscale is confined to, its number of inducing points (the reference locations $M_r$ of Note 1), the coordinate scale $\rho$, and the weight of the log-fold-change penalty. Four belong to the spatial smoothness penalty and the basal spatial field. They are the weight of the penalty (where 0 disables it), the radius at which it is evaluated, and the number of random features and the bandwidth of the field. Two belong to the niche-conditioned modulation head, namely the width of the niche code and the size of the neighborhood from which it is computed. Each setting was trained at three random seeds (42, 1 and 2), and each trained model is scored on the log-fold-change rank correlation and the log-fold-change RMSE used in the benchmark.

To judge a change against the variation between runs of the same configuration, we also trained the reported configuration at five random seeds. We call the interval between the lowest and highest of these five scores the seed range. A setting is described as outside the seed range when its mean across its three seeds, plus or minus the standard error, lies entirely outside that interval. This criterion is a descriptive screen, applied to each setting separately. The seed range is estimated from five runs and each setting from three runs.

**Perturb-map.** The analysis uses the four Perturb-map slides in the seen regime, and every model is scored on the same 12 slide $\times$ perturbation evaluations (Supplementary Fig. 3 and Supplementary Fig. 4). The lengthscale band is swept over 1–6, 3–18 and 9–54 spatial units against 60–360 in the reported configuration. The inducing points take 16, 32, 64 and 128, the coordinate scale 10, 20, 40 and 70, the smoothness weight 0, 0.5, 1 and 2, and the smoothness radius 2, 4 and 8 spatial units. The log-fold-change weight takes 0.5, 1 and 2, the random features 64, 128 and 256, the bandwidth 0.05, 0.15 and 0.30, the niche code width 4, 8 and 16, and the neighborhood size 10, 25 and 50 cells. In total 77 models were trained.

Across the five reference seeds, rank correlation ranged from 0.443 to 0.566 and RMSE from 1.93 to 2.19. All settings remained within this seed range for rank correlation. For RMSE, only the 9–54 spatial-unit lengthscale band fell outside the range, with a lower RMSE of 1.87. The largest changes in the three-seed mean relative to the reported configuration were 0.051 in rank correlation (32 inducing points) and 0.14 in RMSE (niche-code width 4 and the 9–54 lengthscale band).

**SPAC-seq.** We repeated the design on the five-slide SPAC-seq model reported in the main text, in the seen regime. Every model is scored on the same 100 slide $\times$ perturbation evaluations spread over the five slides (Supplementary Fig. 5 and Supplementary Fig. 6). The swept values are those listed for Perturb-map, except that the inducing points take 8, 16 and 64, the coordinate scale 10, 20 and 40, and the smoothness weight 0, 1 and 2. In total 65 models were trained.

Across the five reference seeds, rank correlation ranged from 0.399 to 0.405 and RMSE from 3.769 to 3.788. The inducing-point count, coordinate scale, smoothness radius and number of random features remained within the seed range at every setting for both metrics. The largest reductions in rank correlation occurred with a lengthscale band of 1–6 spatial units, a smoothness weight of 2, niche-code widths of 4 or 16, and a neighborhood of 50 cells. The corresponding RMSE analysis identified the same smoothness and niche-code settings, together with the 9–54 spatial-unit lengthscale band, as less favorable than the reference range.

**Comparing the two datasets.** The seed range is about 19 times narrower on SPAC-seq than on Perturb-map, so a change of a given size can lie outside the range on SPAC-seq and inside it on Perturb-map. The two datasets differ in assay resolution, tissue, number of perturbations, and the number of evaluations averaged per model (12 against 100). Each of these can affect run-to-run variation, and this analysis measures their combined effect. We therefore report the results per dataset.

Supplementary Fig. 7 places the two sweeps on a common axis. Each setting is plotted as its difference from the midpoint of that dataset's own seed range, divided by half the width of that range, so the shaded band spans the seed range on both datasets. Supplementary Fig. 8 repeats the comparison on the RMSE.

#### Note 10: Attributes encoded and implementation details per dataset

CONCERT disentangles a cell's expression into a basal embedding and one embedding per encoded attribute, so which attributes are encoded is a per-dataset choice. The main text describes the model using SPAC-seq, where the encoded attribute is the knockout target; slide-to-slide variation is absorbed by a per-slide bias in the decoder. This note lists the attributes and the task-specific settings for the remaining datasets.

**Perturb-map data.** The model is trained on the four Perturb-map slides jointly. The perturbation (Jak2-KO, Tgfbr2-KO, Ifngr2-KO, or unperturbed) and the cell type (tumor, normal, or periphery) are encoded in the disentanglement module, each by its own codebook. The model uses 2,260 genes, the top 2,000 highly variable genes and 260 further panel genes included by name. In the benchmark, a $4 \times 4$ grid per slide defines the niches. The dataset serves two purposes. It is a complementary benchmark at spot resolution, and it carries the case-study counterfactuals, in which a knockout patch is transplanted between niches and each knockout is placed into niches sampled across a slide.

**Mouse colon spatial transcriptomics data.** We train CONCERT on all 52 slices from the 15 mice, collected at the different time points before and after perturbation. The encoded attributes are the perturbation identity, a categorical variable taking the values DSS or control, and the day, a continuous variable taking the values 0, 3, 9 and 21 and encoded through the Fourier-feature path of Note 3. All 942 genes of the panel are used. The task is to predict the expression profile of each mouse's colon at the time points other than its sampling day.

**Mouse stroke data.** We train CONCERT on the eight brain sections, four from a mouse after photothrombotic stroke and four from a sham control, encoding the perturbation label and the tissue region. The perturbation label distinguishes the infarct core and the peri-infarct area in the models used for the core–rim comparison, and is a single stroke label in the 2D cortex-versus-thalamus and lesion-size model. The region labels are those provided with the original study. The registration of the sections and the three-dimensional arm are described in Note 4, and the marker screen and the two counterfactual protocols in Note 11.

**Perturb-FISH data.** The Perturb-FISH tumor section is used for the kernel-parameter analysis in Note 5. The model is trained on the single section with knockout target encoded in the disentanglement module and control cells assigned to the control label, so the action-correction term is active. The 31 knockout targets carried by at least 25 cells are encoded. The spatial components use the SPAC-seq settings: a niche code of width 8 computed from the 25 nearest cells, $g_{\max} = 3$, 16 inducing points, a 3–18 spatial-unit lengthscale band, coordinate scale 20, spatial-smoothness penalty of weight 1 and radius 4 spatial units, and log-fold-change penalty of weight 1. Coordinates are divided by the median nearest-neighbor distance and then rescaled to $[0, 20]$. Training runs for up to 800 epochs with early stopping.

**Spatial hyper-parameters per dataset.** Perturb-map and SPAC-seq differ in the geometry of the perturbation footprint field and share the other spatial settings. Training differs in early stopping, with a patience of 300 epochs on Perturb-map and 50 on SPAC-seq, both within a maximum of 1,500 epochs. Both use a niche code of width 8 computed from the 25 nearest cells, a spatial smoothness penalty of weight 1 evaluated at a radius of 4 spatial units, and a log-fold-change penalty of weight 1. Both also use a basal spatial field of 128 random features at bandwidth 0.15, with cell coordinates rescaled to a range of 20. The footprint field uses 64 inducing points and a lengthscale band of 60–360 spatial units on Perturb-map, against 16 inducing points and a band of 3–18 spatial units on SPAC-seq.

Two further settings follow from the assays. The action-correction term, which is identified by control cells, is active on SPAC-seq, which carries non-targeting guides, and switched off on Perturb-map. A SPAC-seq perturbation must also reach 25 cells before it is encoded, and Perturb-map encodes all three of its knockouts. The results of exchanging the two footprint settings between the datasets are given in Note 9.

#### Note 11: Marker availability and counterfactual protocol in the stroke case study

We screened all 2,000 modelled genes on the four stroke sections against the anatomical labels from the original study, using healthy cortex as the cortical (CTX) reference in the stroke brain. A candidate core marker was required to be detected in more than half of infarct-core (ICA) spots, elevated by more than 0.5 in log space over healthy cortex, and higher in core than in peri-infarct area (PIA) by more than 0.2. Gm42418 showed the strongest core association (core-versus-rim contrast $+2.42$) and was used as the core marker for the across-section comparison. Lesion extent, summarized by a pooled five-gene injury score, is the primary readout; core-marker agreement provides a complementary readout.

The two counterfactual protocols address distinct questions. In the free protocol, the perturbed region is recovered by thresholding the predicted injury field. In the shape-matched protocol, the observed ICA and PIA masks are transferred onto the sham brain, fixing lesion geometry while predicting the transcriptional response. We therefore report the protocols separately.

The 2D cortex-versus-thalamus, lesion-size and multi-focal counterfactuals come from a 2D model with a single stroke label, trained with the paracrine term disabled, and they are read out on the injury genes Lcn2, Vim and Timp1. The 2D and 3D models used for the protocol comparison above are also trained with the paracrine term disabled.

#### Supplementary Tables

Both supplementary tables are provided in a separate file:

`CONCERT_Supplementary_tables.xlsx`

**Supplementary Table 1**. Significance of the comparisons between CONCERT and each competing method on the log-fold-change rank correlation. Each row is one competing method, with the raw and Benjamini–Hochberg adjusted $p$-values for the three benchmark arms: within-niche prediction on SPAC-seq, cross-slide prediction on SPAC-seq, and within-niche prediction on Perturb-map. $P$-values are from two-sided Wilcoxon signed-rank tests over matched evaluation records. A record is a (perturbation, slide, niche) unit in the within-niche arms and a (perturbation, target slide) pair in the cross-slide arm, scored by both methods, and each test uses the records that the competing method shares with CONCERT.

**Supplementary Table 2**. Significance of the same comparisons on the log-fold-change RMSE, in the layout of Supplementary Table 1. A lower RMSE indicates predicted effect sizes closer to the measured ones.

#### Supplementary Figures

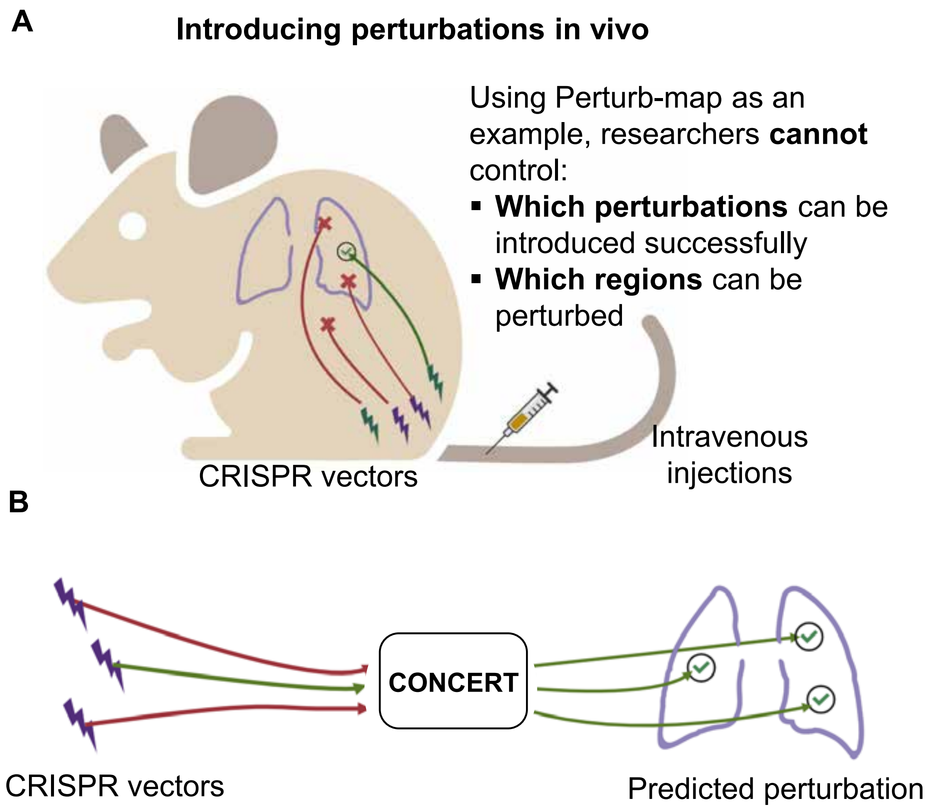


**Supplementary Fig. 1: Overview of spatial perturbation tasks**. Potential issues in current spatial perturbation sequencing technologies (a) and the motivation to develop CONCERT (b).

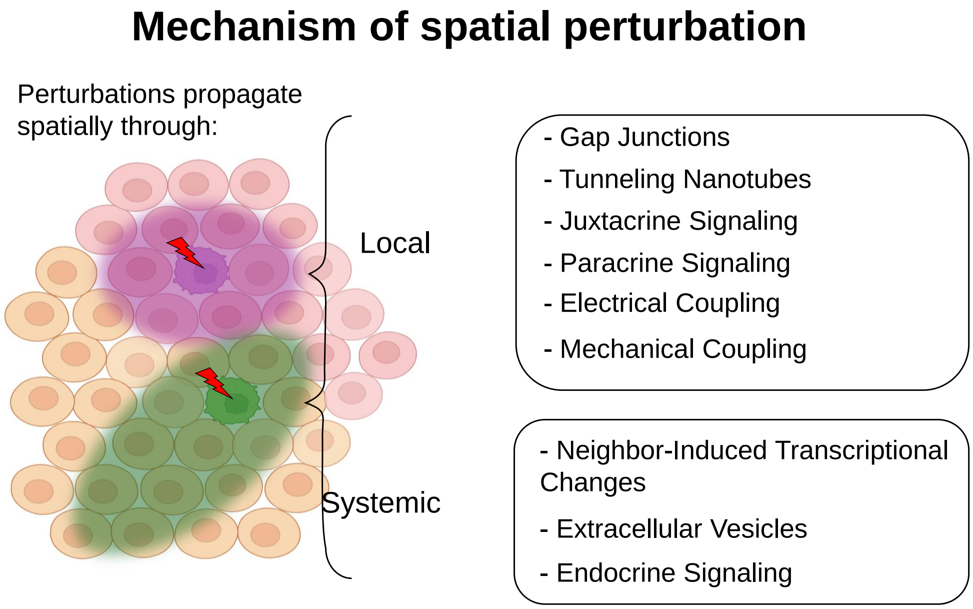


**Supplementary Fig. 2:** Mechanisms of the dispersion of perturbation effects in tissue.

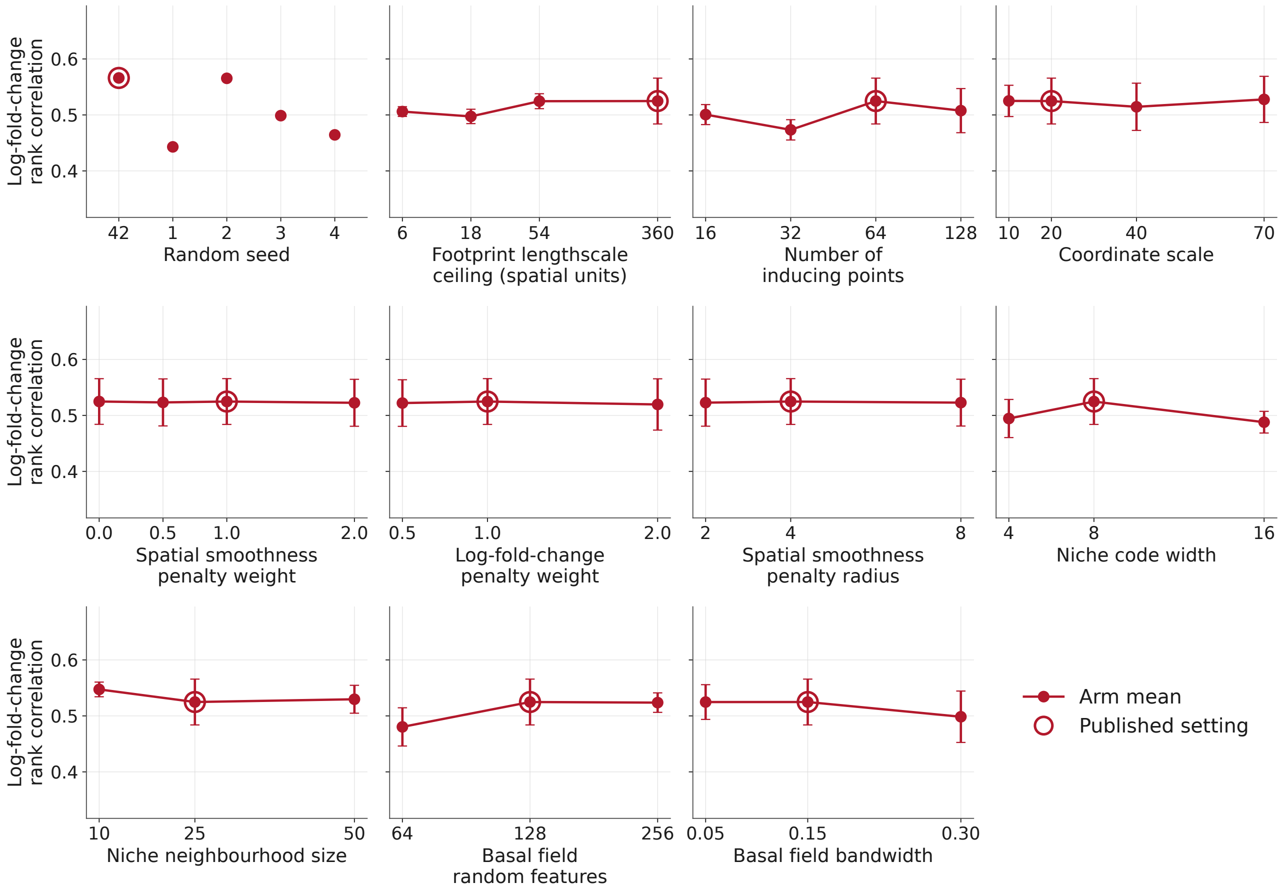


**Supplementary Fig. 3: Sensitivity of CONCERT to the hyperparameters of its spatial components on Perturb-map, measured by log-fold-change rank correlation.** All four slides are used in this test. Each point is the mean over three models trained with different random seeds, and each model is averaged over its 12 combinations (slide $\times$ perturbation); error bars are the standard error across the three seeds. The first panel shows five runs of the reported configuration that differ only in random seed, and sets the scale against which the remaining panels are read. The open ring marks the reported setting. Every other panel adjusts one hyperparameter with all others held at their reported values.

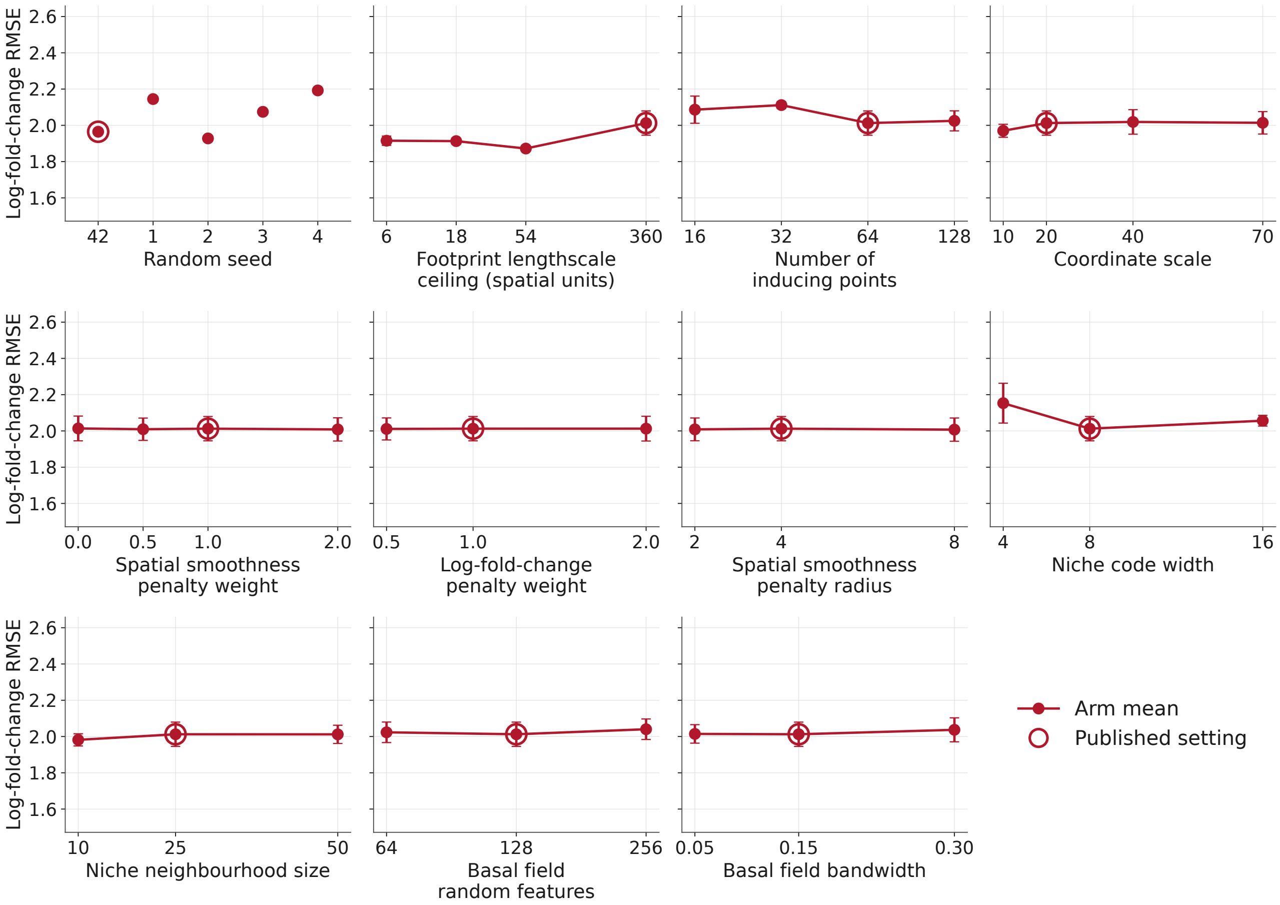


**Supplementary Fig. 4: Sensitivity of CONCERT to the hyperparameters of its spatial components on Perturb-map, measured by log-fold-change RMSE.** Panels, seeds and error bars are the same as in Supplementary Fig. 3. Lower is better.

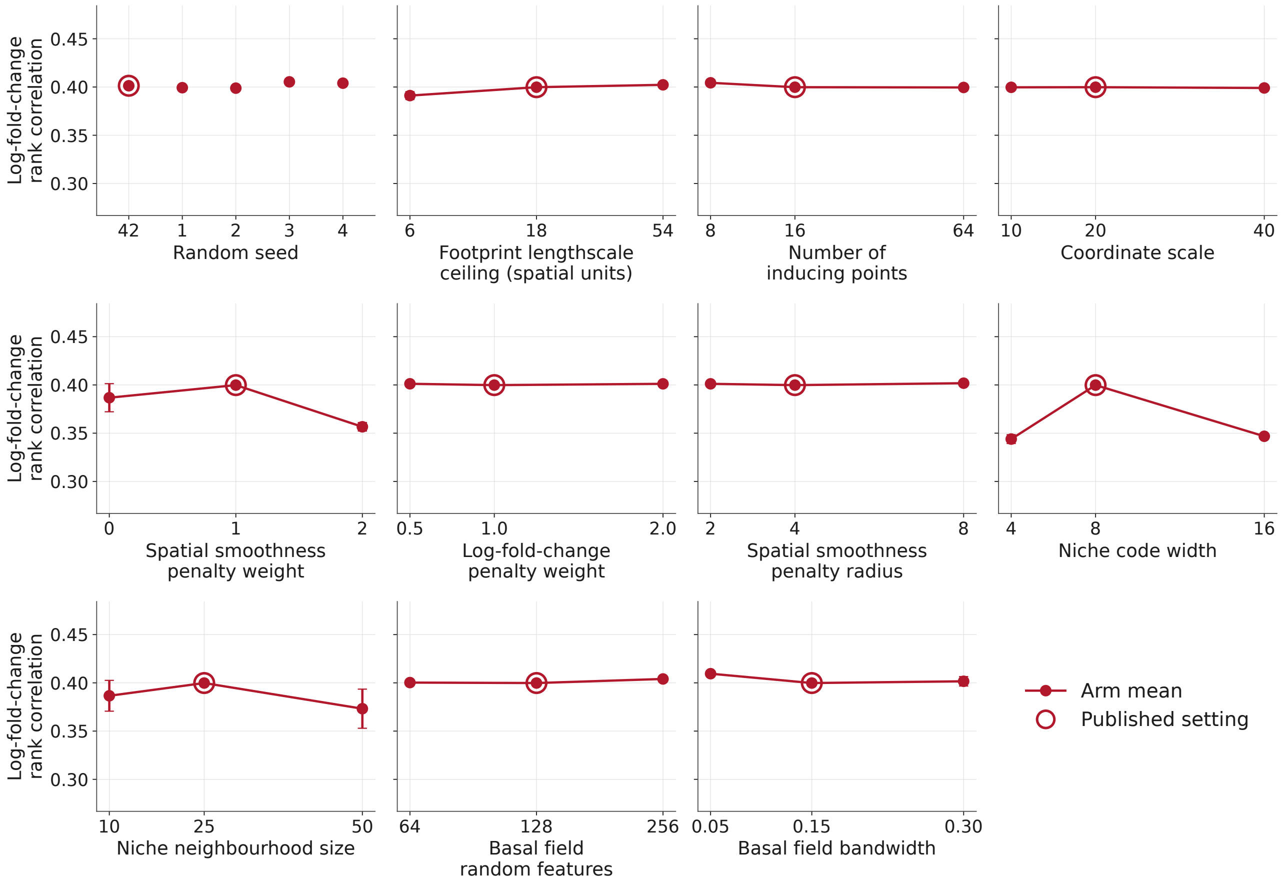


**Supplementary Fig. 5: Sensitivity of CONCERT to the hyperparameters of its spatial components on SPAC-seq, measured by log-fold-change rank correlation.** All five slides are used in this test. Panels, seeds and error bars follow Supplementary Fig. 3, with each model averaged over 100 combinations (slide $\times$ perturbation).

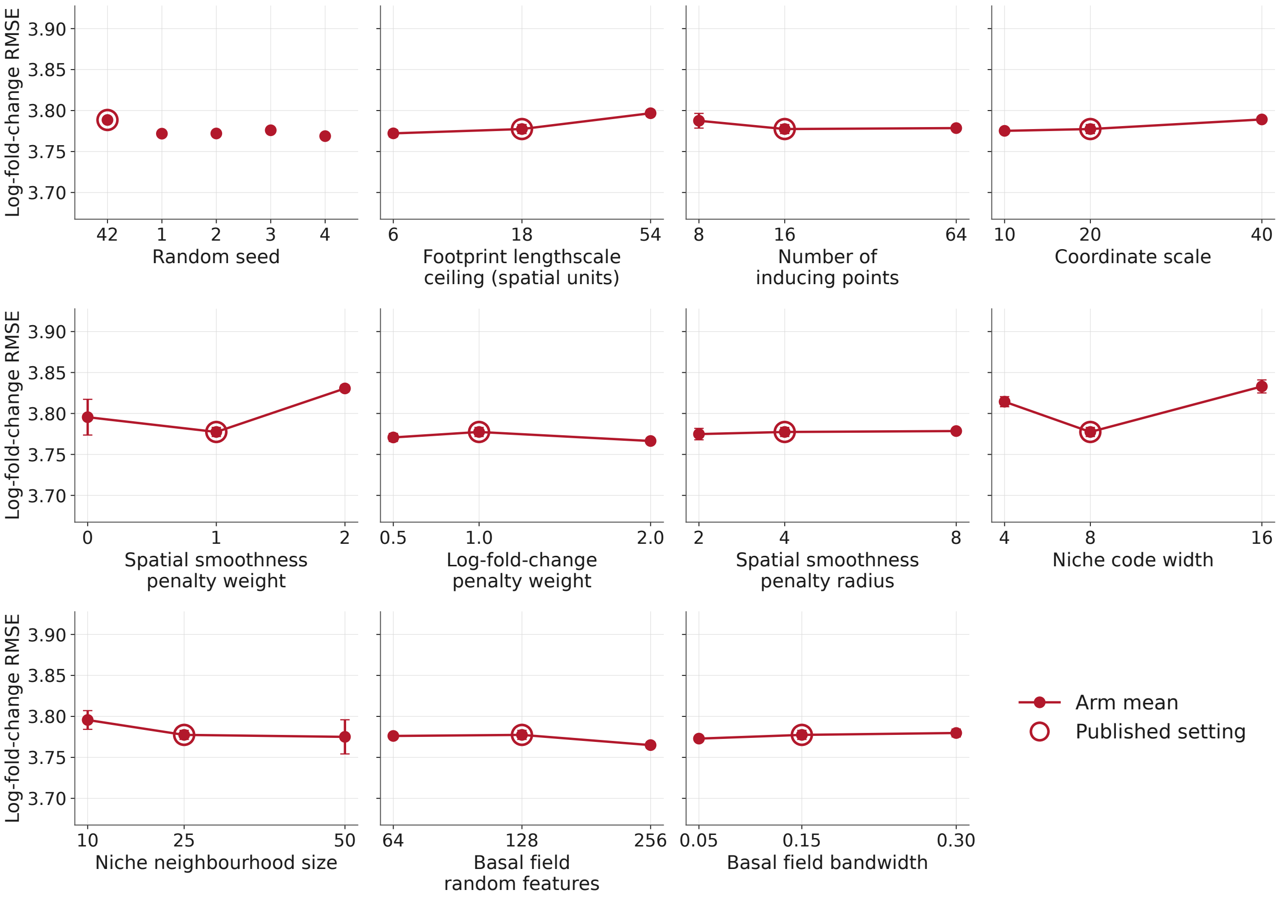


**Supplementary Fig. 6: Sensitivity of CONCERT to the hyperparameters of its spatial components on SPAC-seq, measured by log-fold-change RMSE.** Panels and error bars are the same as in Supplementary Fig. 5. Lower is better.

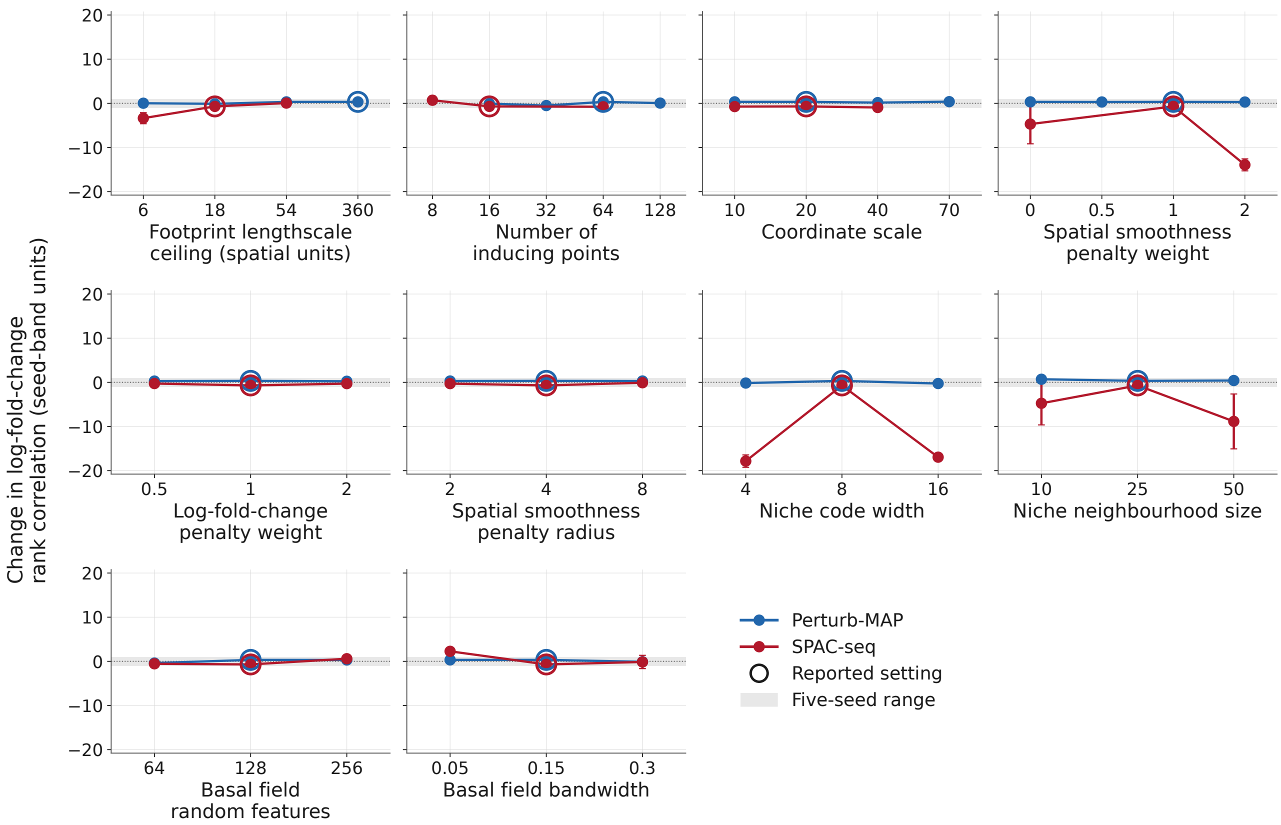


**Supplementary Fig. 7: Sensitivity of CONCERT to the hyperparameters of its spatial components on Perturb-map and SPAC-seq, shown on a common standardized scale of log-fold-change rank correlation.** Each panel varies one hyperparameter while all others are held at that dataset's reported values. Blue, Perturb-map; red, SPAC-seq. Each point is the mean over three models trained with random seeds, and each model's score is averaged over its 12 (Perturb-map) or 100 (SPAC-seq) combinations (slide × perturbation). The two datasets differ in baseline accuracy and in run-to-run variation, so each mean is standardized within its own dataset. For each dataset, five models of the reported configuration that differ only in random seed define a seed range. A point's value is its difference from the midpoint of that range, divided by half the width of the range. On this scale the seed range runs from $-1$ to $+1$ on both datasets (gray band), and positive values indicate a higher rank correlation. A setting is considered to differ from the reported configuration only when its mean ± s.e.m. lies entirely outside the band. Error bars indicate the s.e.m. across the three seeds on the same scale. Open circles show the reported setting of each dataset. Settings are placed at equally spaced positions. Unstandardized values are shown in Supplementary Fig. 3 and Supplementary Fig. 5.

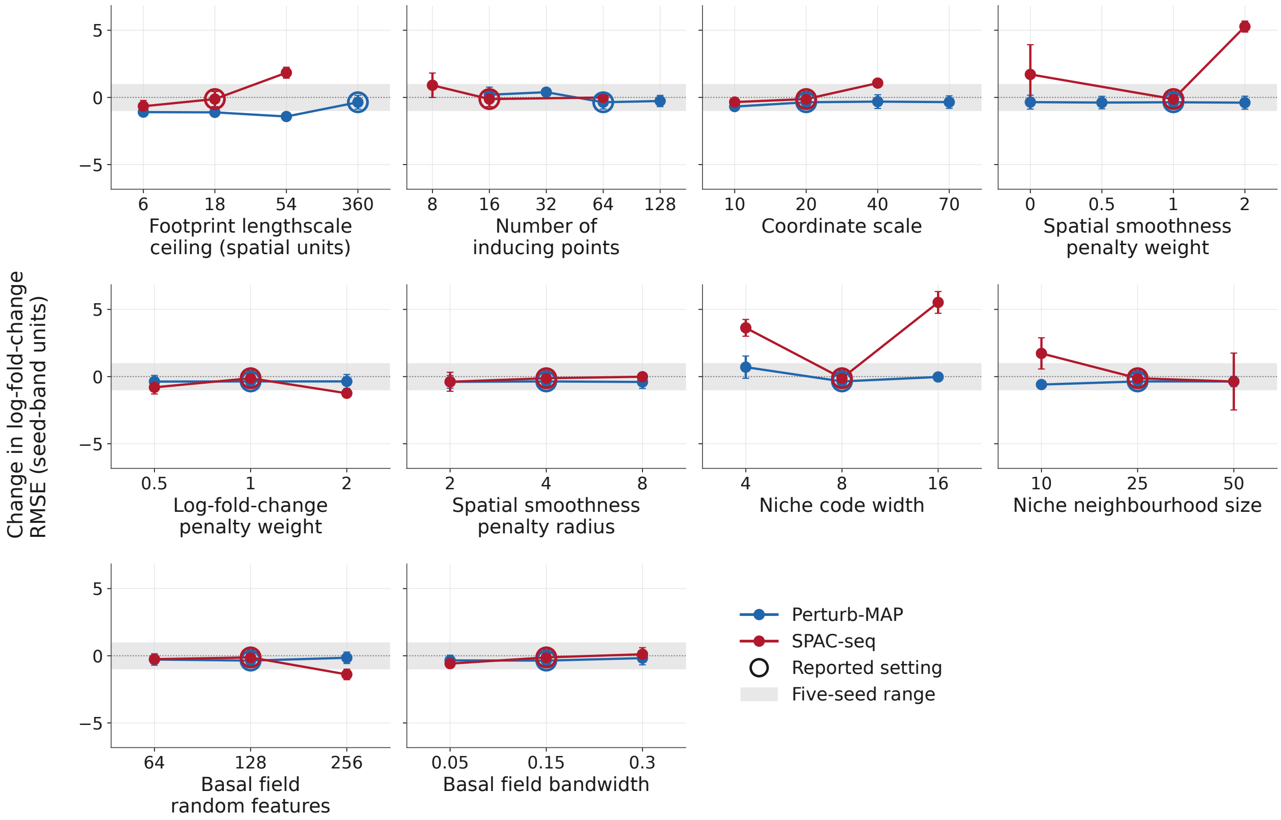


**Supplementary Fig. 8: Sensitivity of CONCERT to the hyperparameters of its spatial components on Perturb-map and SPAC-seq, shown on a common standardized scale (log-fold-change RMSE).** Panels, standardization, grey band, error bars and open circles are the same as in Supplementary Fig. 7. Here positive values indicate a higher RMSE, that is, a lower accuracy. Unstandardized values are shown in Supplementary Fig. 4 and Supplementary Fig. 6.

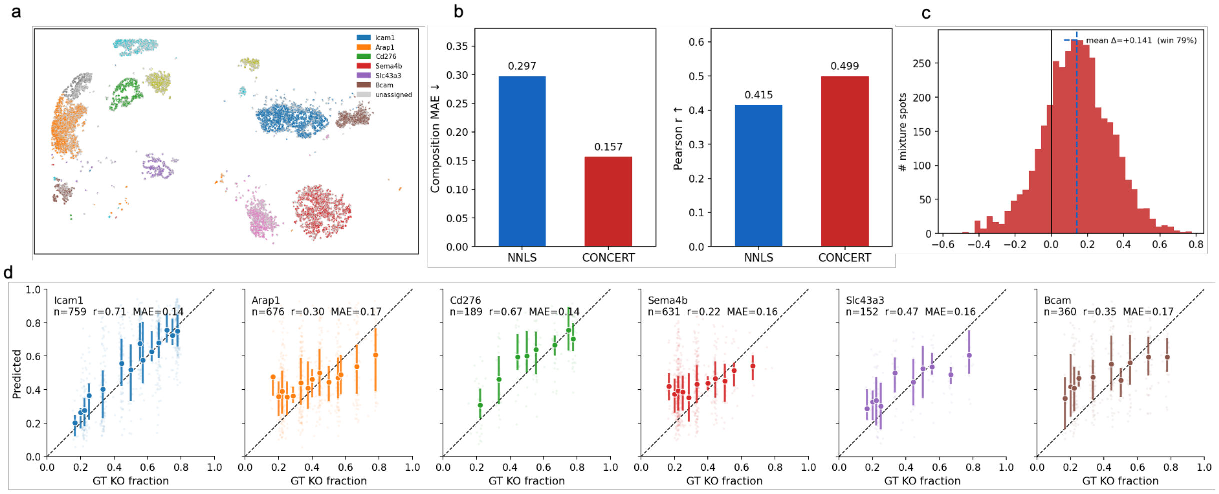


**Supplementary Fig. 9: Validation of CONCERT deconvolution on SPAC-seq.** **a**, SPAC-seq slide 2-1 rendered as pseudo-bulk spots: single cells are aggregated into spots of known composition, so each spot's true perturbation composition is available as ground truth. **b**, Spot composition accuracy for CONCERT and NNLS: mean absolute error (left, lower is better) and Pearson correlation (right) between predicted and true composition. **c**, Per-spot difference in absolute composition error, $|\mathrm{err}_{\mathrm{NNLS}}| - |\mathrm{err}_{CONCERT}|$, across all mixture spots; positive values favour CONCERT. **d**, Predicted knockout fraction against the true fraction for six representative perturbations; points with bars are the binned mean $\pm$ s.d. and the dashed line is the identity.

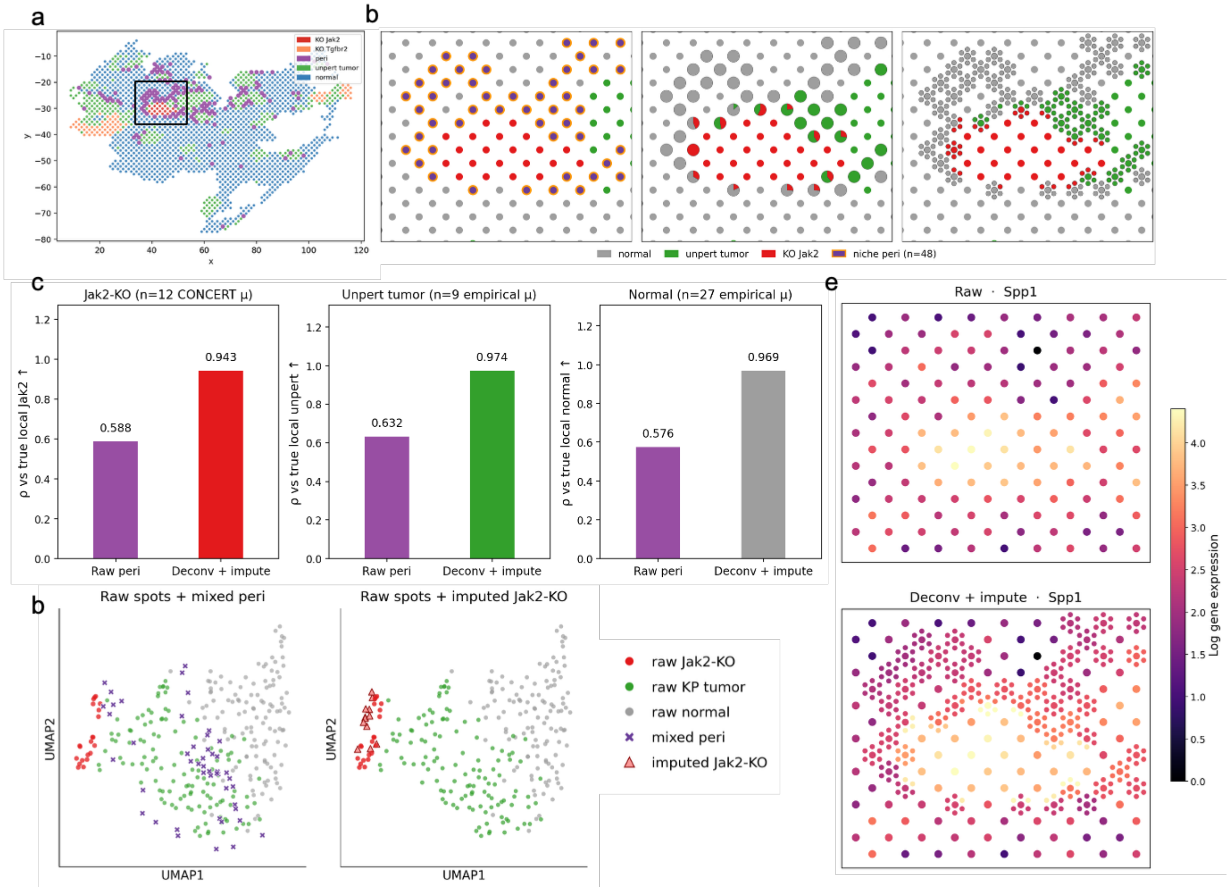


**Supplementary Fig. 10: Deconvolution of periphery spots in Perturb-map.** **a**, The Jak2-knockout tumor patch on Perturb-map slide 0 and the surrounding periphery spots analyzed here. **b**, The same region shown as raw spots (left), as deconvolved cell-type composition (middle), and as deconvolved sub-spots (right). **c**, Agreement with the local profile of each population, for the raw periphery spot and for the profile that CONCERT predicts for that population at the spot location. **d**, UMAP of Perturb-map spots. Left, raw spots colored by identity, with mixed periphery spots (crosses); right, the Jak2-knockout components predicted by CONCERT (triangles). **e**, Expression of Spp1 in the same region at spot level (top) and sub-spot level (bottom).

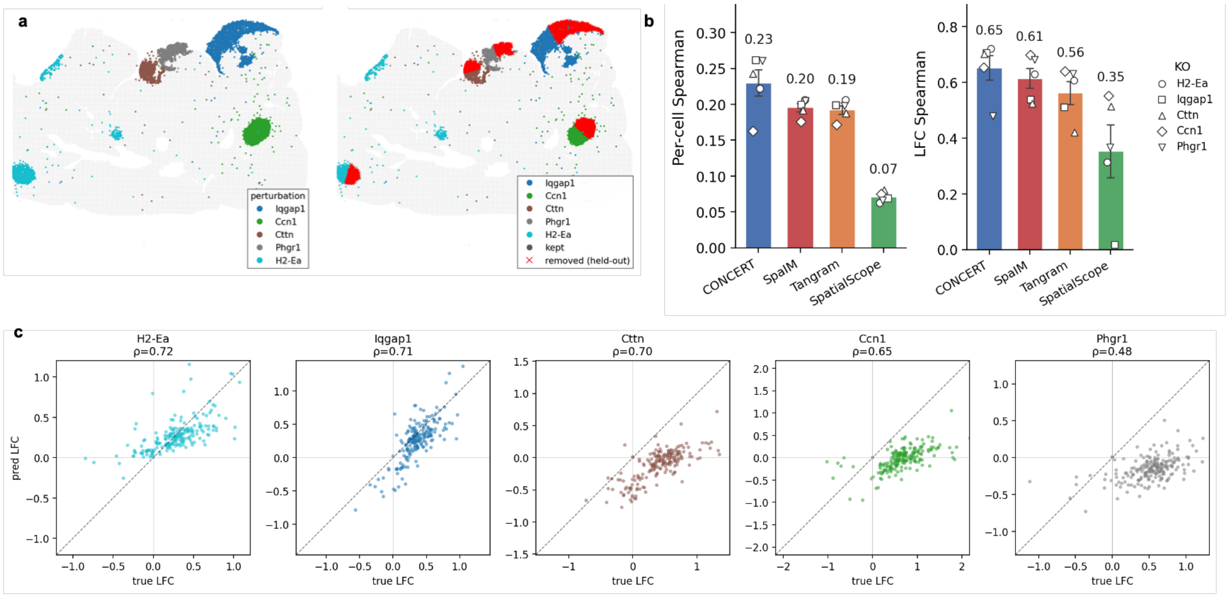

**Supplementary Fig. 11: Imputation of held-out perturbed patches on SPAC-seq. a**, SPAC-seq slide showing the patch layout, with the held-out patches highlighted; all cells inside a held-out patch are withheld from the model and their expression is predicted. **b**, Imputation accuracy over the five held-out knockout patches for CONCERT and three imputation methods. Left, mean per-cell Spearman correlation between imputed and observed expression. Right, Spearman correlation between imputed and observed patch log-fold change. The observed log-fold change of a patch is the mean of its knockout cells minus the mean of nearby unassigned cells. The imputed log-fold change uses the same nearby unassigned cells for the imputation methods, and the imputed unperturbed expression of the same cells for CONCERT. **c**, Observed against CONCERT-imputed patch log-fold change for each held-out patch, on the top-200 highly variable genes; the dashed line is the identity and $\rho$ is the Spearman correlation.

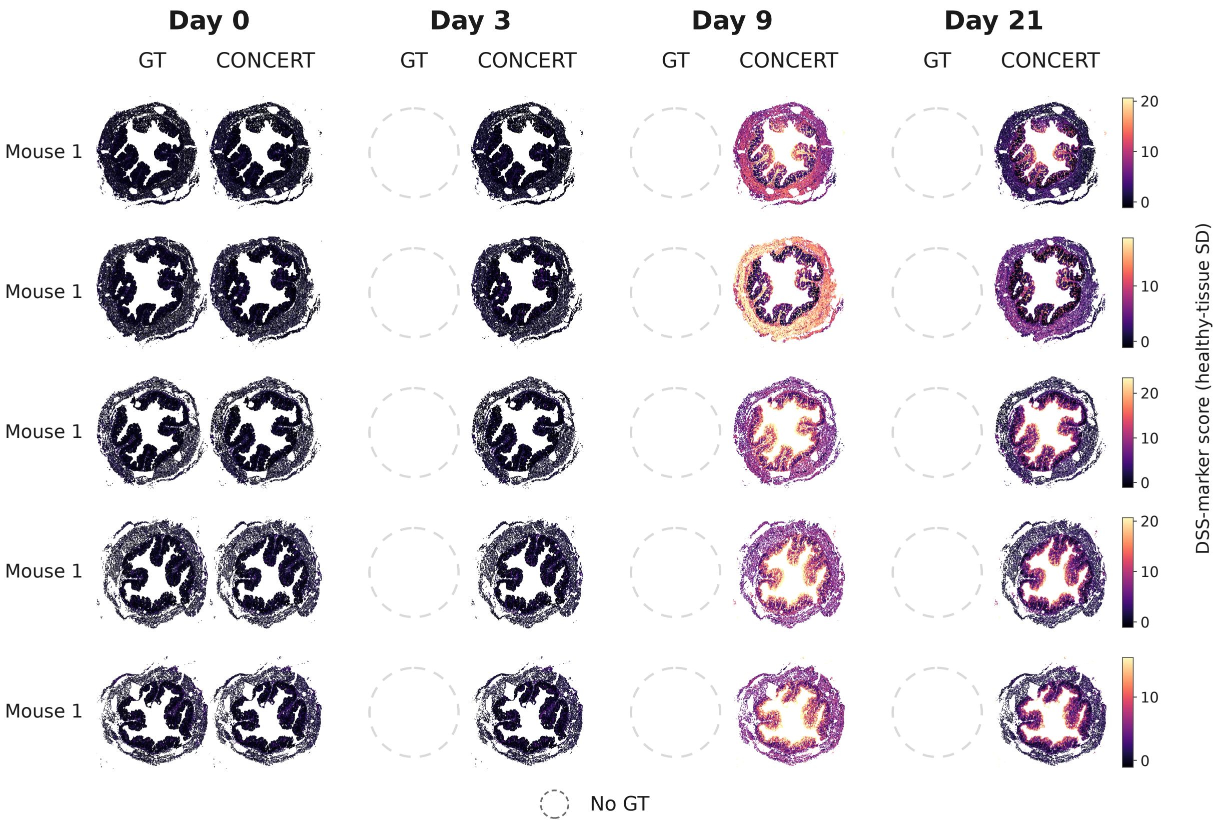


**Supplementary Fig. 12: Predicted and observed inflammation field on the five slides of mouse `062921_D0_m3a`, sampled at day 0.** Color is the continuous DSS-marker score in healthy-tissue s.d. units. Rows are the slides of this animal. Columns are the four sampling days, each split into the observed measurement and the CONCERT prediction. Because sampling is terminal, an observed panel exists only for the day on which this animal was sampled, and the other observed slots are drawn as empty circles.

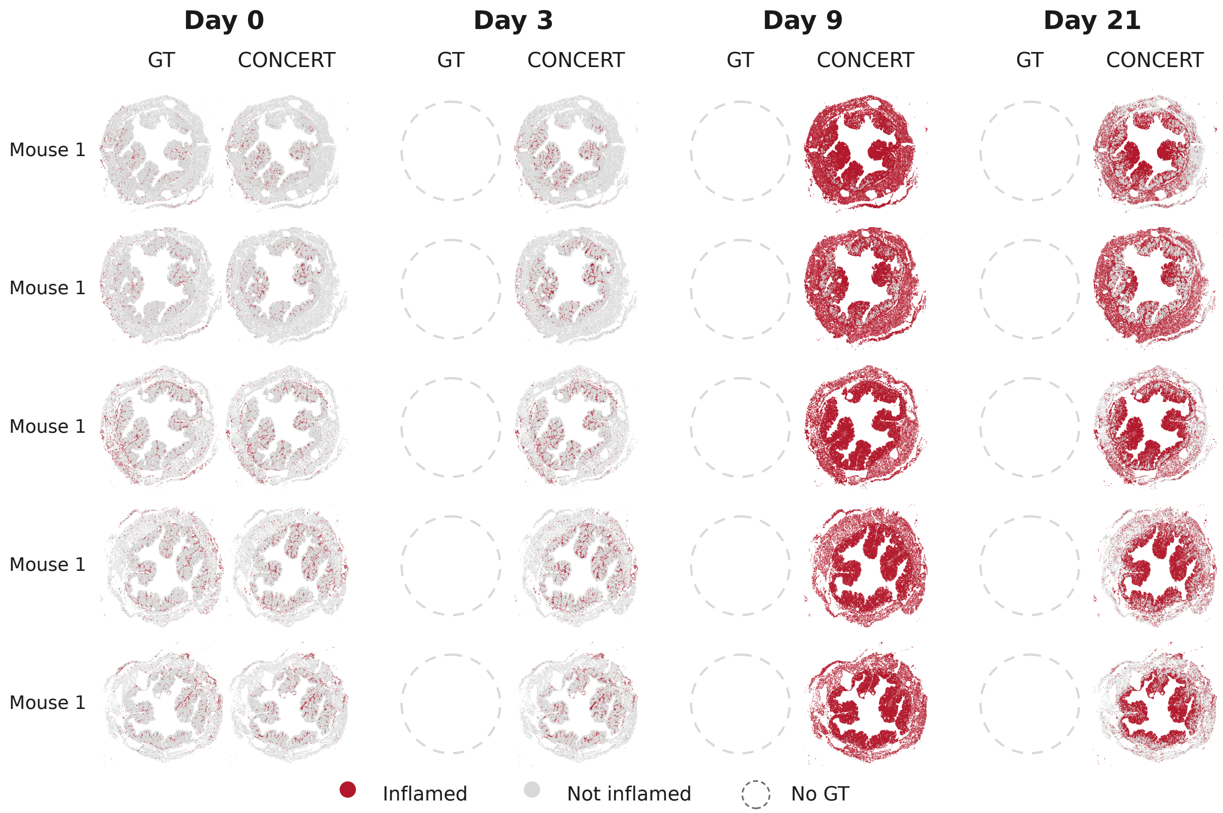


**Supplementary Fig. 13: Predicted and observed inflamed region on the five slides of mouse `062921_D0_m3a`, sampled at day 0.** Binary inflamed region called from the field in Supplementary Fig. 12 with the rate-matched cutoffs used in Fig. 5e. Rows, columns and empty circles are the same as in Supplementary Fig. 12.

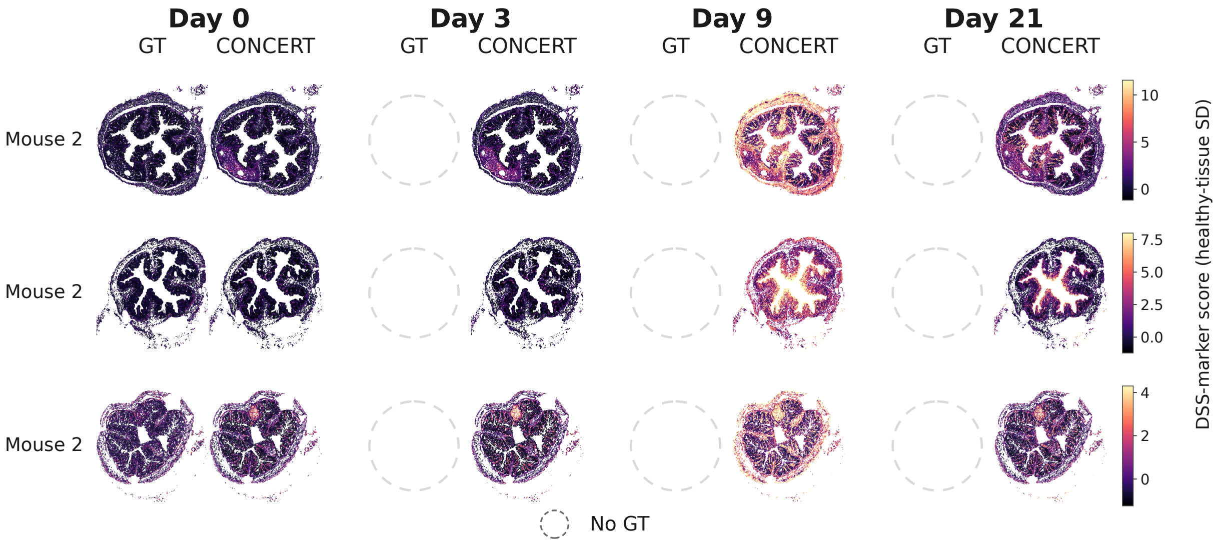


**Supplementary Fig. 14: Predicted and observed inflammation field on the three slides of mouse `082421_D0_m6`, sampled at day 0.** Color is the continuous DSS-marker score in healthy-tissue s.d. units. Rows are the slides of this animal. Columns are the four sampling days, each split into the observed measurement and the CONCERT prediction. Because sampling is terminal, an observed panel exists only for the day on which this animal was sampled, and the other observed slots are drawn as empty circles.

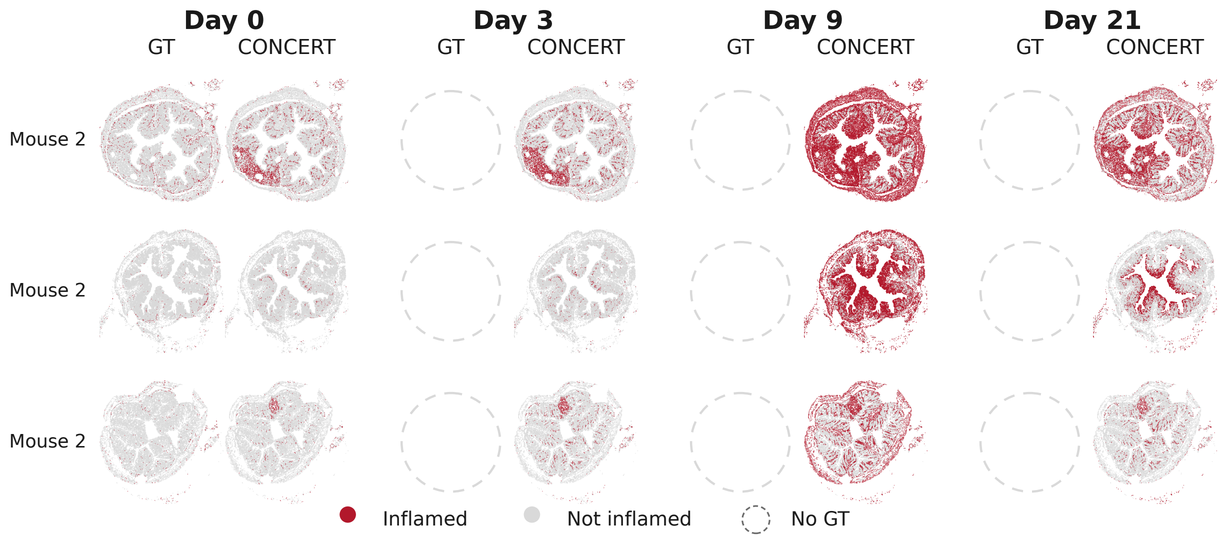


**Supplementary Fig. 15: Predicted and observed inflamed region on the three slides of mouse `082421_D0_m6`, sampled at day 0.** Binary inflamed region called from the field in Supplementary Fig. 14 with the rate-matched cutoffs used in Fig. 5e. Rows, columns and empty circles are the same as in Supplementary Fig. 14.

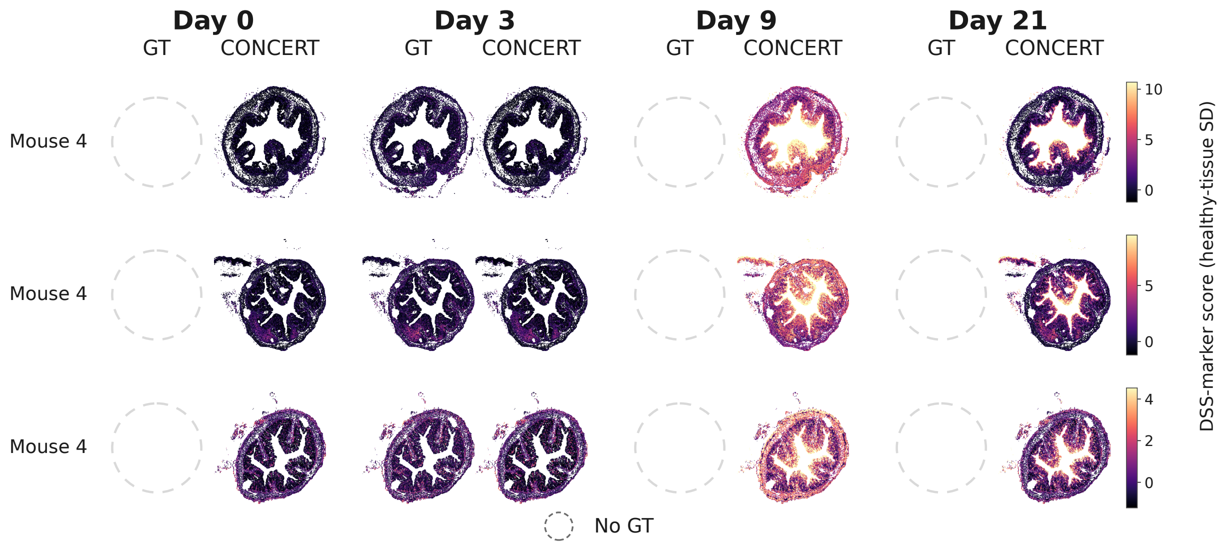


**Supplementary Fig. 16: Predicted and observed inflammation field on the three slides of mouse `092421_D3_m1`, sampled at day 3.** Color is the continuous DSS-marker score in healthy-tissue s.d. units. Rows are the slides of this animal. Columns are the four sampling days, each split into the observed measurement and the CONCERT prediction. Because sampling is terminal, an observed panel exists only for the day on which this animal was sampled, and the other observed slots are drawn as empty circles.

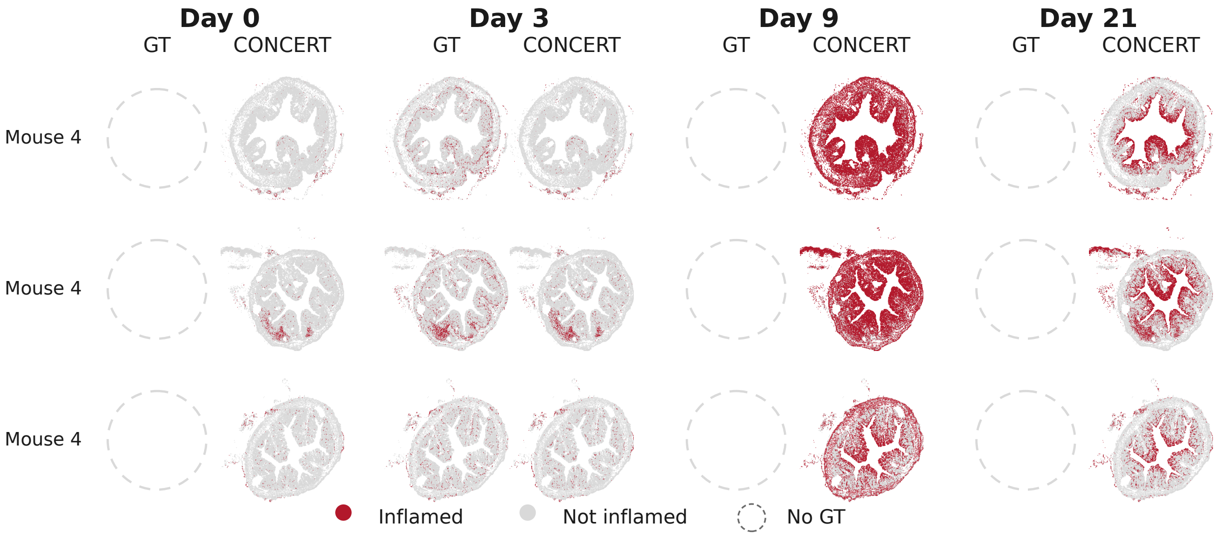


**Supplementary Fig. 17: Predicted and observed inflamed region on the three slides of mouse `092421_D3_m1`, sampled at day 3.** Binary inflamed region called from the field in Supplementary Fig. 16 with the rate-matched cutoffs used in Fig. 5e. Rows, columns and empty circles are the same as in Supplementary Fig. 16.

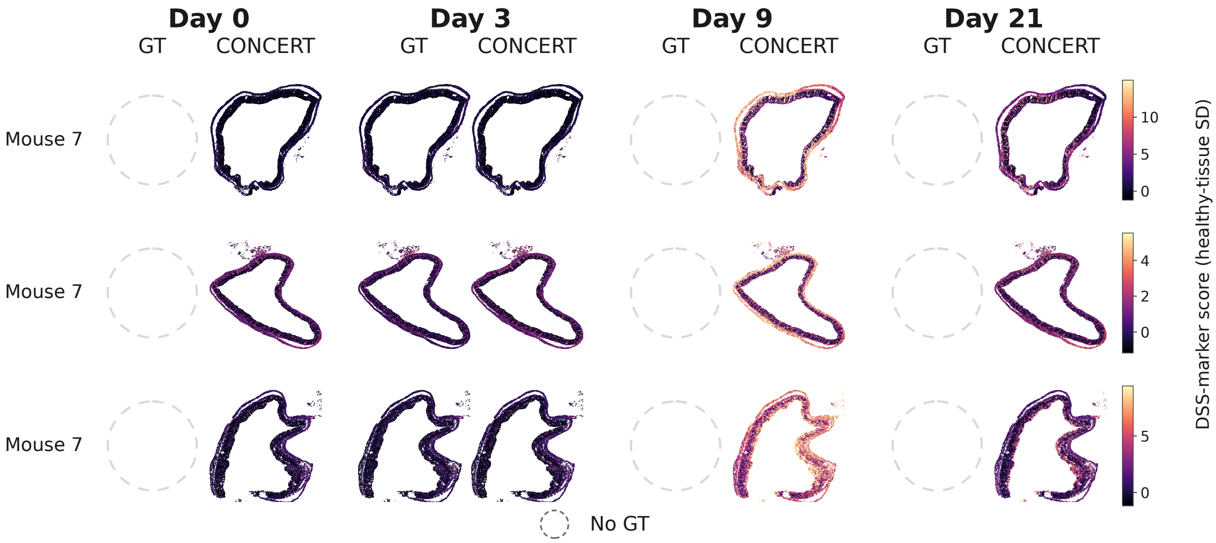


**Supplementary Fig. 18: Predicted and observed inflammation field on the three slides of mouse `092421_D3_m4`, sampled at day 3.** Color is the continuous DSS-marker score in healthy-tissue s.d. units. Rows are the slides of this animal. Columns are the four sampling days, each split into the observed measurement and the CONCERT prediction. Because sampling is terminal, an observed panel exists only for the day on which this animal was sampled, and the other observed slots are drawn as empty circles.

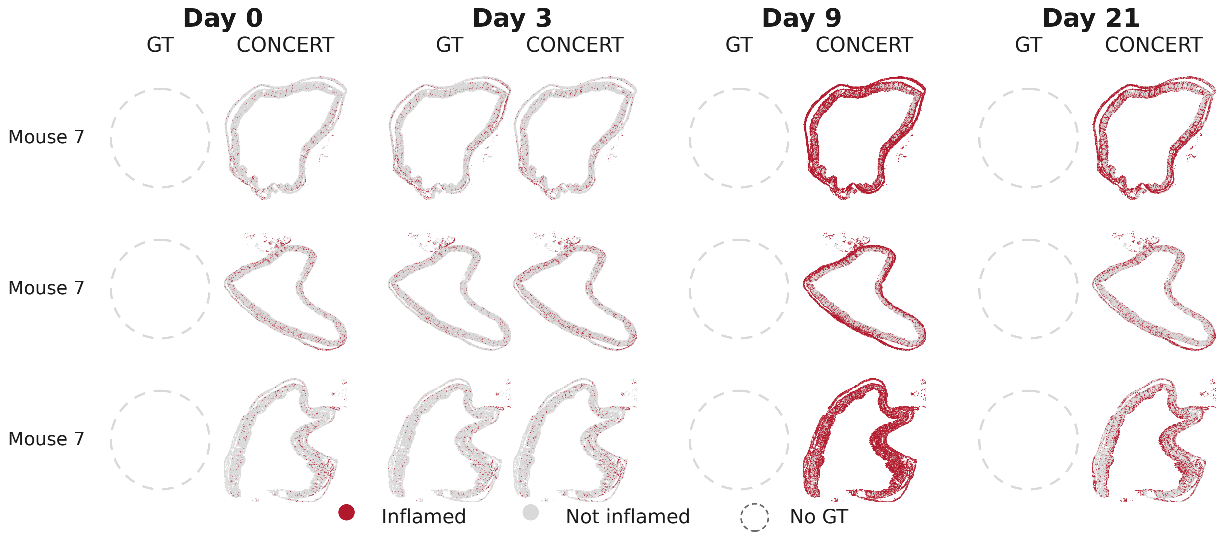


**Supplementary Fig. 19: Predicted and observed inflamed region on the three slides of mouse `092421_D3_m4`, sampled at day 3.** Binary inflamed region called from the field in Supplementary Fig. 18 with the rate-matched cutoffs used in Fig. 5e. Rows, columns and empty circles are the same as in Supplementary Fig. 18.

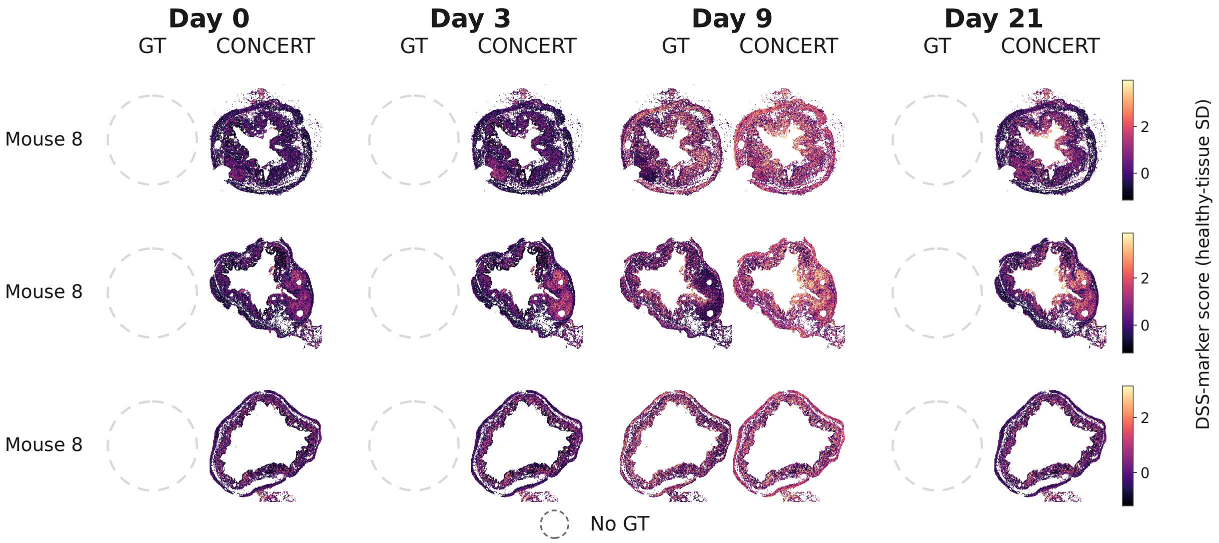


**Supplementary Fig. 20: Predicted and observed inflammation field on the three slides of mouse `062221_D9_m3`, sampled at day 9.** Color is the continuous DSS-marker score in healthy-tissue s.d. units. Rows are the slides of this animal. Columns are the four sampling days, each split into the observed measurement and the CONCERT prediction. Because sampling is terminal, an observed panel exists only for the day on which this animal was sampled, and the other observed slots are drawn as empty circles.

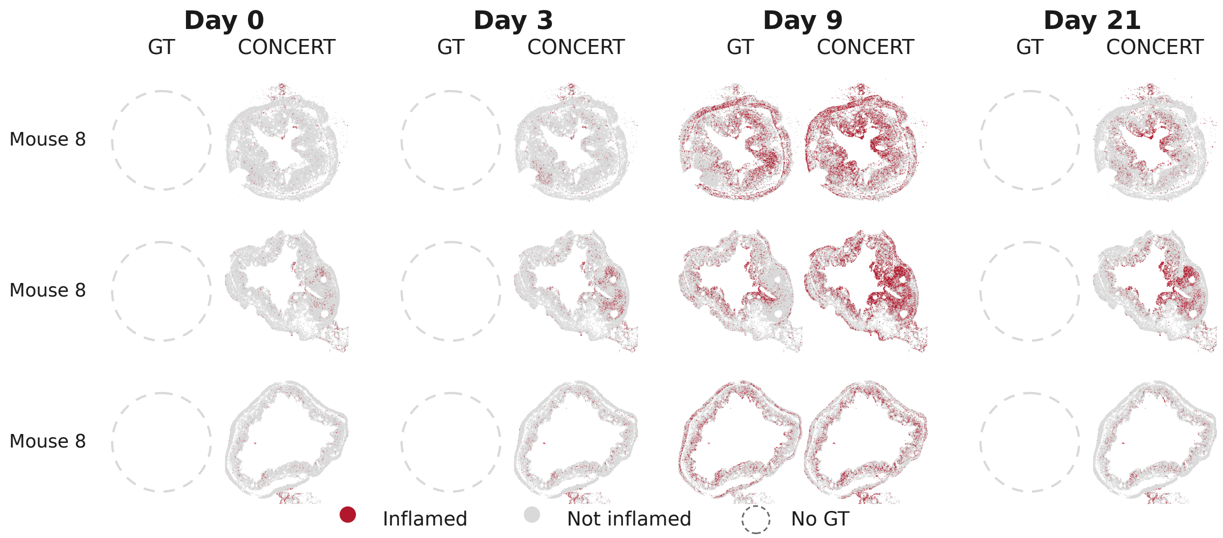


**Supplementary Fig. 21: Predicted and observed inflamed region on the three slides of mouse `062221_D9_m3`, sampled at day 9.** Binary inflamed region called from the field in Supplementary Fig. 20 with the rate-matched cutoffs used in Fig. 5e. Rows, columns and empty circles are the same as in Supplementary Fig. 20.

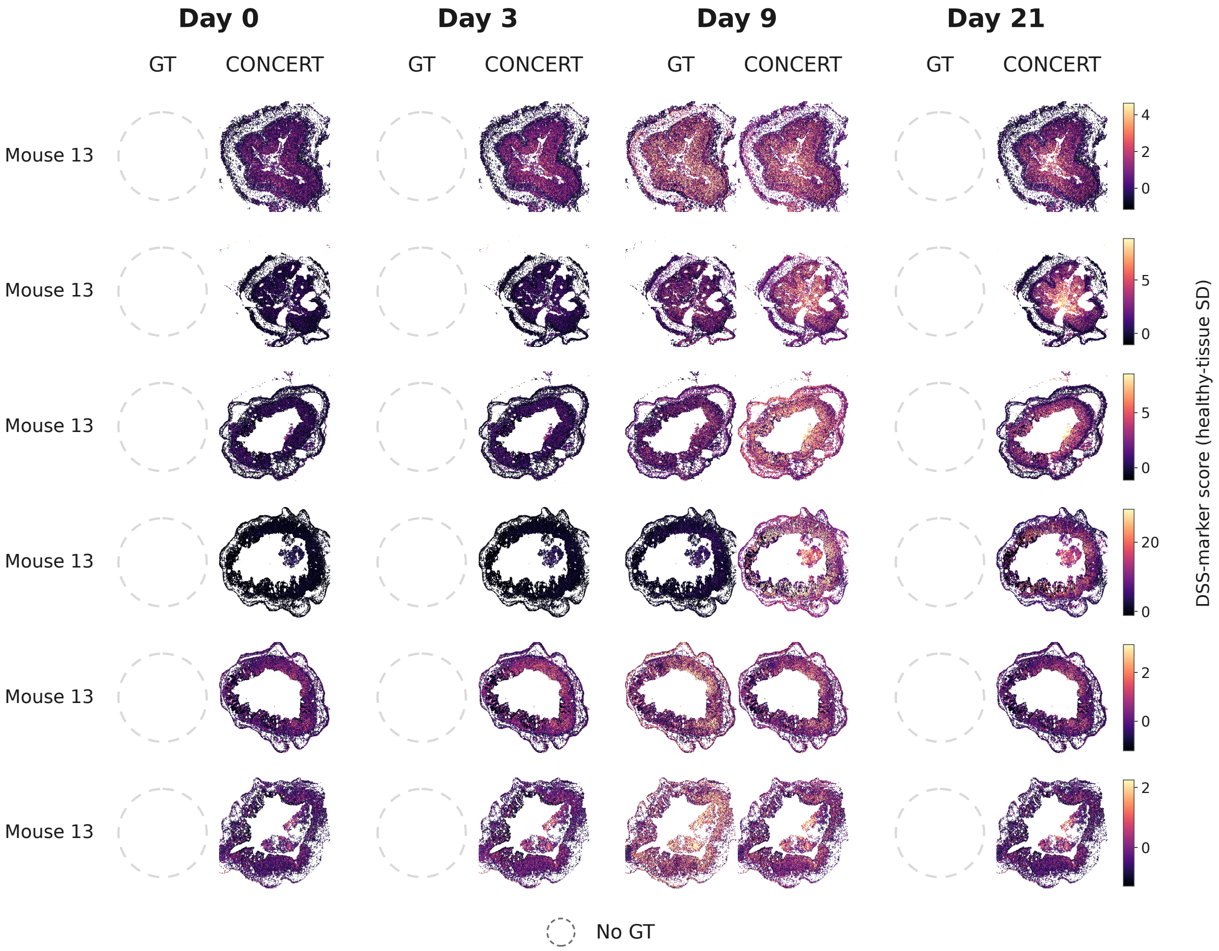


**Supplementary Fig. 22: Predicted and observed inflammation field on the six slides of mouse `100221_D9_m5`, sampled at day 9.** Color is the continuous DSS-marker score in healthy-tissue s.d. units. Rows are the slides of this animal. Columns are the four sampling days, each split into the observed measurement and the CONCERT prediction. Because sampling is terminal, an observed panel exists only for the day on which this animal was sampled, and the other observed slots are drawn as empty circles.

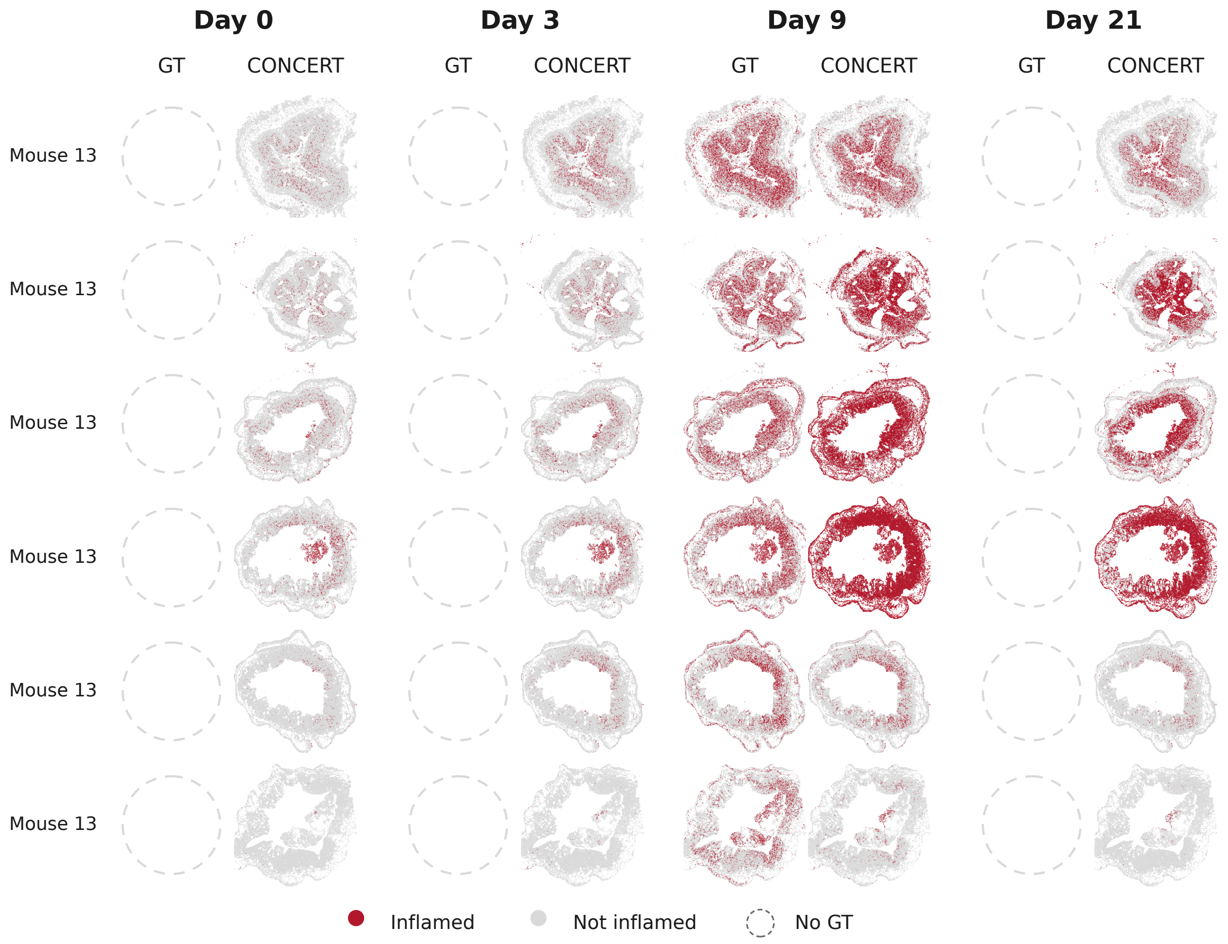


**Supplementary Fig. 23: Predicted and observed inflamed region on the six slides of mouse `100221_D9_m5`, sampled at day 9.** Binary inflamed region called from the field in Supplementary Fig. 22 with the rate-matched cutoffs used in Fig. 5e. Rows, columns and empty circles are the same as in Supplementary Fig. 22.

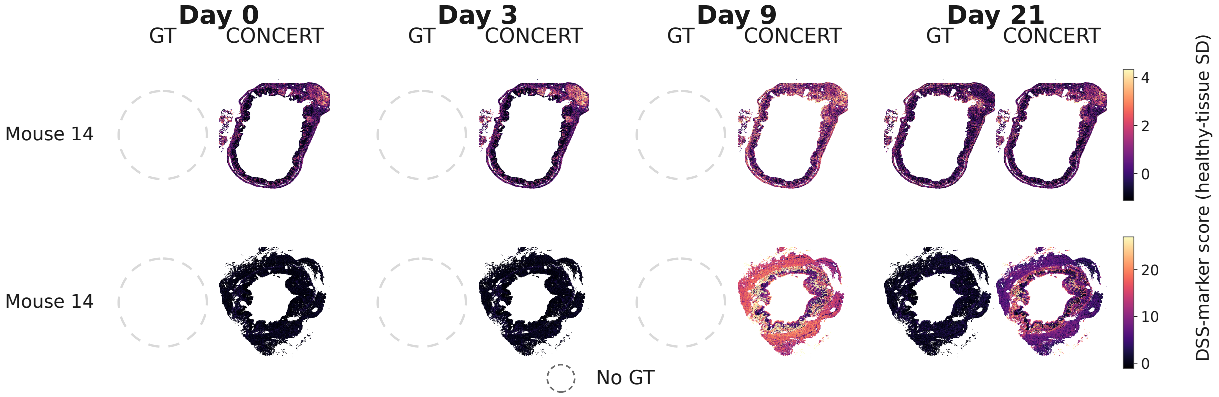


**Supplementary Fig. 24: Predicted and observed inflammation field on both slides of mouse `082421_D21_m1`, sampled at day 21.** Color is the continuous DSS-marker score in healthy-tissue s.d. units. Rows are the slides of this animal. Columns are the four sampling days, each split into the observed measurement and the CONCERT prediction. Because sampling is terminal, an observed panel exists only for the day on which this animal was sampled, and the other observed slots are drawn as empty circles.

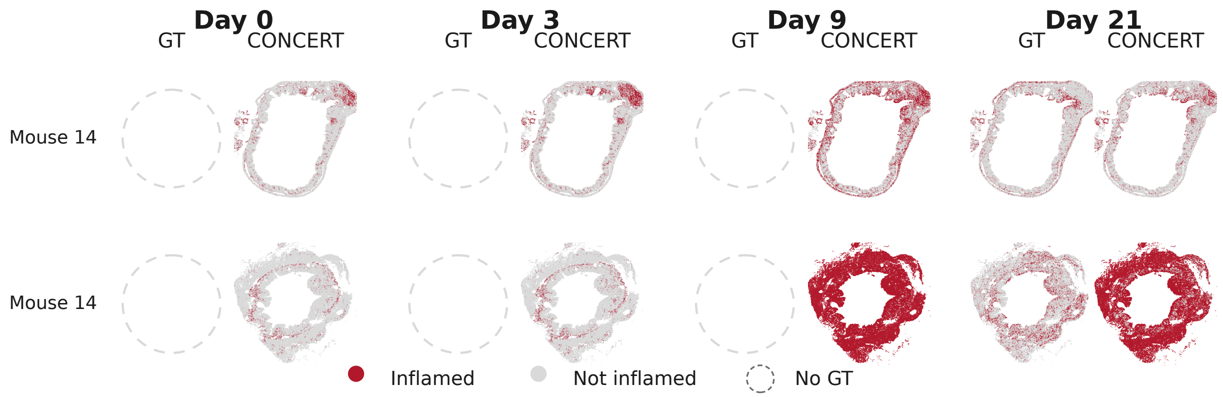


**Supplementary Fig. 25: Predicted and observed inflamed region on both slides of mouse `082421_D21_m1`, sampled at day 21.** Binary inflamed region called from the field in Supplementary Fig. 24 with the rate-matched cutoffs used in Fig. 5e. Rows, columns and empty circles are the same as in Supplementary Fig. 24.

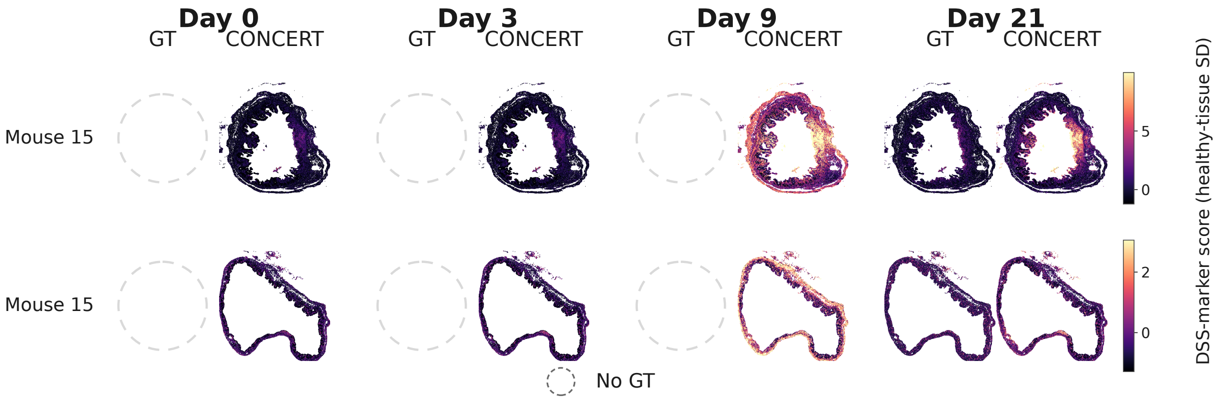


**Supplementary Fig. 26: Predicted and observed inflammation field on both slides of mouse `082421_D21_m2`, sampled at day 21.** Color is the continuous DSS-marker score in healthy-tissue s.d. units. Rows are the slides of this animal. Columns are the four sampling days, each split into the observed measurement and the CONCERT prediction. Because sampling is terminal, an observed panel exists only for the day on which this animal was sampled, and the other observed slots are drawn as empty circles.

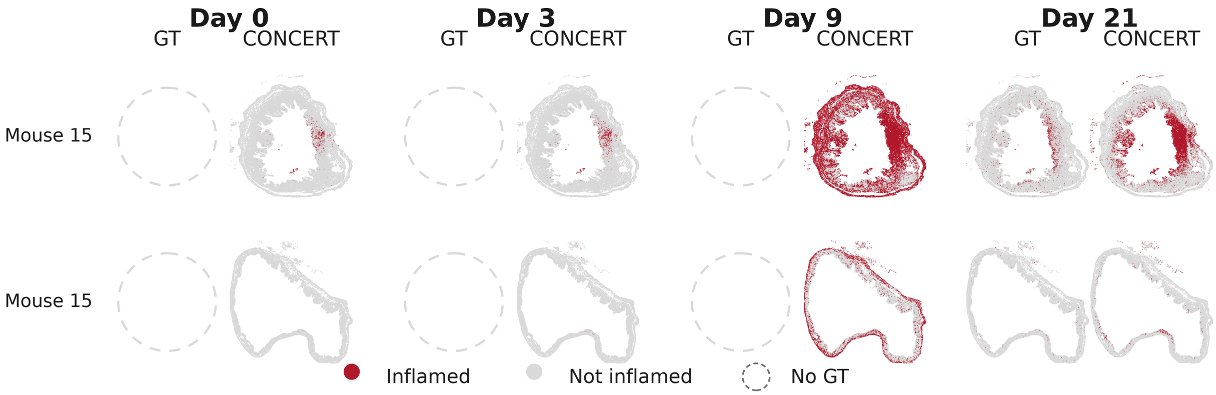


**Supplementary Fig. 27: Predicted and observed inflamed region on both slides of mouse `082421_D21_m2`, sampled at day 21.** Binary inflamed region called from the field in Supplementary Fig. 26 with the rate-matched cutoffs used in Fig. 5e. Rows, columns and empty circles are the same as in Supplementary Fig. 26.

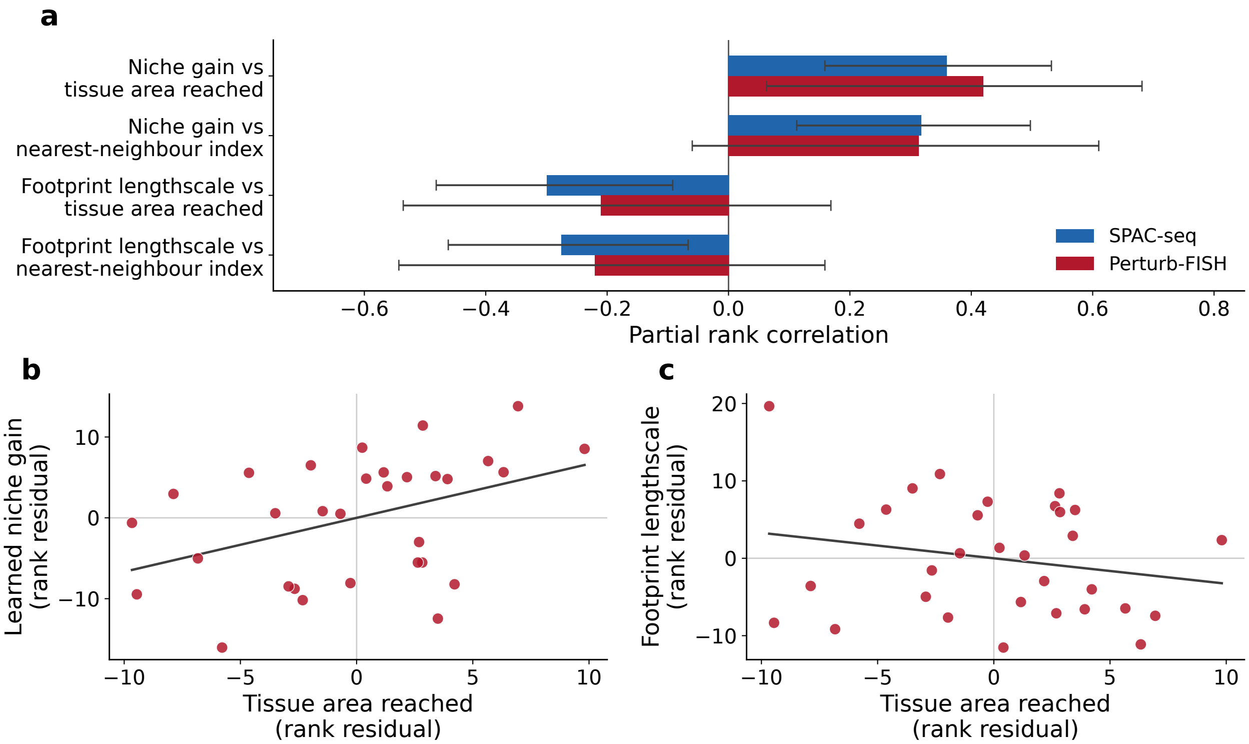


**Supplementary Fig. 28: Learned spatial parameters against the observed layout of each perturbation on Perturb-FISH and SPAC-seq. a**, Partial rank correlation between each of two learned parameters and each of two layout descriptors. Blue, the SPAC-seq within-slide model over the 87 knockouts scored in the benchmark; red, a Perturb-FISH model trained in the same configuration over the 31 knockouts carried by at least 25 cells. The niche gain sets how strongly the local microenvironment may modulate a perturbation's predicted response, and the footprint lengthscale sets how far its spatial effect field extends. The first descriptor is the fraction of section area that the perturbed cells reach. The second is the nearest-neighbor index, the mean distance from a perturbed cell to the nearest cell carrying the same perturbation divided by the distance expected under a random scatter of the same number of cells. Both correlations control for the number of perturbed cells and the norm of the perturbation embedding. Bars are the partial correlation and whiskers its 95% interval on the Fisher scale. **b**, **c**, Perturb-FISH rank residuals, after the same two controls, for the niche gain (**b**) and the footprint lengthscale (**c**) against the fraction of section area reached. One dot per knockout; the line is the least-squares fit. Values and corrected $P$-values are given in Supplementary Note 5.

1. Liu, X., Zeira, R. & Raphael, B. J. Partial alignment of multislice spatially resolved transcriptomics data. *Genome Research* **33**, 1124–1132 (2023).
